# Supramolecular ‘catch-and-release’ strategy for bioorthogonal fluorogenic imaging across the visible spectrum

**DOI:** 10.1101/2023.04.24.538015

**Authors:** Ranjan Sasmal, Arka Som, Pratibha Kumari, Meenakshi Pahwa, Nilanjana Das Saha, Sushma Rao, Sheeba Vasu, Sarit S. Agasti

## Abstract

Fluorogenic probes that unmask fluorescence signals in response to a bioorthogonal reaction are a powerful new addition to biological imaging. They can provide significantly reduced background fluorescence and minimize non-specific signals, potentially allowing real-time high-contrast imaging without washing out excess fluorophores. While diverse classes of highly refined synthetic fluorophores are readily available now, their integration into a bioorthogonal fluorogenic scheme still necessitates another level of extensive design efforts and customized structural alterations to optimize quenching mechanisms for each given fluorophore scaffold. Herein, we present an easy-to-implement and highly generalizable supramolecular ‘catch-and-release’ strategy for generating an efficient bioorthogonal fluorogenic response from essentially any readily available fluorophores without further structural alterations. We designed this distinct strategy based on the macrocyclic cucurbit[7]uril (CB7) host, where a fluorogenic response is achieved by programming a guest displacement reaction from the macrocycle cavity. We used this strategy to rapidly generate fluorogenic probes across the visible spectrum from structurally diverse classes of fluorophore scaffolds, including coumarin, bodipy, rhodamine, and cyanine. These probes were applied to no-wash fluorogenic imaging of various target molecules in live cells and tissue with minimal background and no appreciable non-specific signal. Notably, the orthogonal reactivity profile of the system allowed us to pair this host-guest fluorogenic probe with the covalently clickable fluorogenic probe to achieve high-contrast super-resolution and multiplexed fluorogenic imaging in cells and tissue.

## Introduction

Understanding living systems and unraveling their fundamental biological processes critically relies on the ability to observe specific biomolecules with a high spatiotemporal resolution in cells and tissues. Such efforts are greatly benefited by combining advanced fluorescence microscopy techniques with appropriate labeling strategies.^1–8^ In recent years, with the advent of bioorthogonal reactions that are robust and compatible with living systems, bioorthogonal strategies have emerged as attractive new labeling tools for advanced biological imaging.^9–14^ It offered an exquisite reactivity-based chemical labeling tool to biology, enabling specific tagging of a diverse set of biomolecules in their native environment with highly refined synthetic fluorophores. However, a major challenge that initially held back its use in a variety of advanced imaging applications is attributed to fluorescence background from unbound or nonspecifically bound synthetic fluorophores post-labeling. Although one could utilize rigorous washing steps to remove excess unbound probes, nonspecifically bound fluorescent probes are even harder to eliminate by washing. In addition, in the case of live cell or *in vivo* conditions, it is impossible to implement a washing step that can possibly clear excess or nonspecifically bound probes. Moreover, washing steps are not well tolerated for staining of scant cell populations (e.g., circulating tumor cells) in a point-of-care microfluidic device or in precise pulse-chase experiments where shorter time interval measurements are desired.^15^ These challenges can be elegantly tackled by utilizing a bioorthogonal fluorogenic labeling scheme, which provides an appealing solution for real-time background-free imaging without washing or clearance steps.^16–19^ The fluorescent probes used for such an approach exhibit an increase in fluorescence upon reacting to their specific biorthogonal counterpart, alleviating the problem of a background signal from unbound or nonspecifically bound probes. While vast libraries of highly refined synthetic fluorophore families are readily available, adopting them for a fluorogenic labeling scheme is yet a significant challenge, necessitating another level of customized structural design efforts. Moreover, increasing the spectrum of fluorogenic bioorthogonal transformations by incorporating motifs with unique orthogonal reactivity profiles is highly desirable to expand the scope of fluorogenic imaging for simultaneous probing of multiple cellular targets (i.e., multiplexed imaging).

**Scheme 1.**
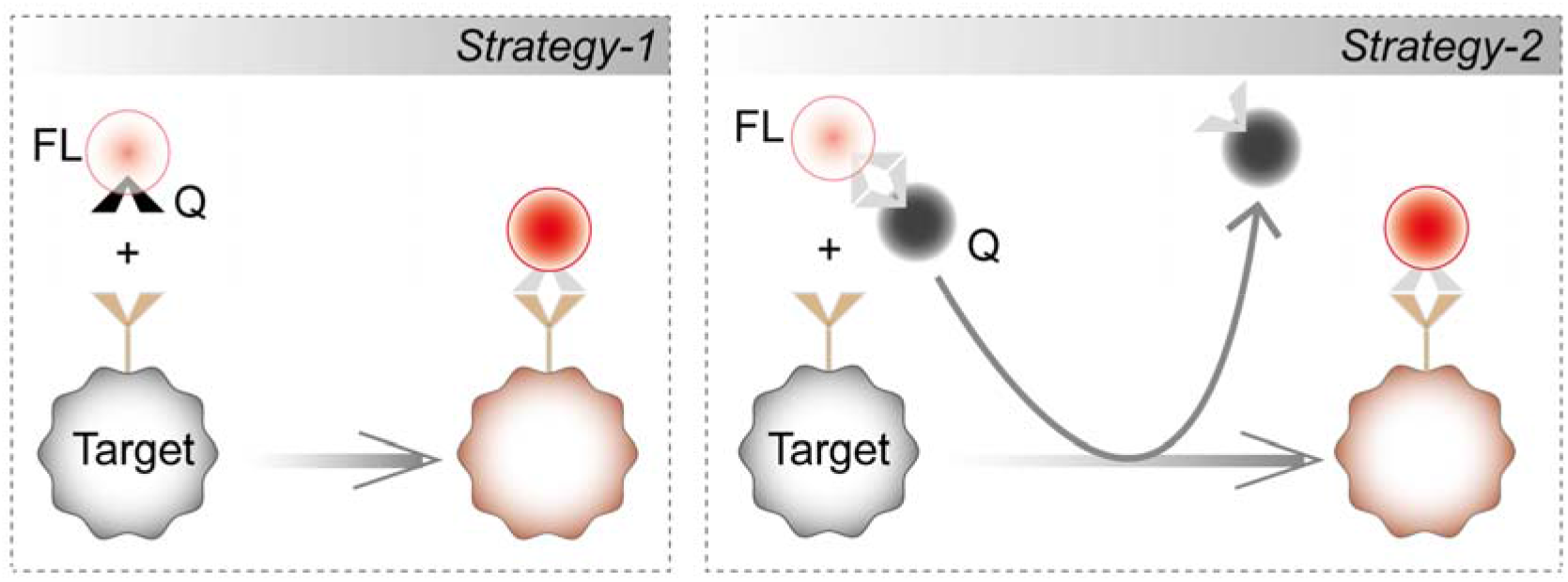
General design strategies for bioorthogonal fluorogenic imaging.

The most common design principle for a bioorthogonal fluorogenic probe is to strategically quench the intrinsic fluorescence of a dye until a specific biorthogonal reaction eliminates this quenching effect, restoring the latent fluorescence.^20^ In rare instances, a dye scaffold has also been generated using a bioorthogonal reaction.^21^ Conceptually quenching of the dyes can be achieved in two ways (Scheme 1): fluorophores can be armed with a suitable quencher moiety that can be either chemically converted to a spectroscopically non-perturbing functionality or eliminated during the labeling process.^20^ In the first concept, bioorthogonal reactive groups (azide/alkyne or tetrazine) are strategically incorporated into the dye skeleton such that they can act as a quencher moiety for the dye.^22–27^ In this case, the quenching effect is eliminated upon bioorthogonal conversion of the quenching moiety. Arguably, the most important example of this concept is tetrazine (Tz) probes, which gained popularity due to their fast reactivity via inverse-electron-demand Diels–Alder cycloaddition reaction.^28–38^ Tz has been shown to quench fluorescence via various photophysical mechanisms, including Forster resonance energy transfer (FRET),^39, 40^ through-bond energy transfer (TBET),^26, 27^ Dexter-type electron exchange,^41^ and photoinduced electron transfer (PET).^42^ Over the last few years, a range of design strategies has been investigated in collaboration with different fluorophore scaffolds to harness various Tz-mediated quenching mechanisms and achieve fluorogenic tetrazine probes extending up to the far-red/NIR range.^26, 27, 30, 33–42^ While such fluorogenic probes have been promising, the quenching strategies are not generalizable to diverse libraries of fluorophore families, requiring extensive design efforts to customize quenching mechanisms for each fluorophore scaffold. In addition, access to these fluorogenic probes often requires direct alteration of the core fluorophore skeleton, demanding re-optimization of the laborious fluorophore synthesis scheme and tackling critical synthetic challenges. As an alternative way of designing fluorogenic probes for labeling, one might consider adapting the second concept where bioorthogonal reaction can lead to the release of the quenching group. This can take advantage of the readily/commercially available and highly optimized dye-quencher pairs to generate a fluorogenic response. This strategy is highly generalizable, and it can essentially incorporate any fluorophores into a fluorogenic labeling scheme by simply pairing them with a suitable quencher molecule. However, designing an appropriate bioorthogonal reaction scheme that can harness the benefit of this concept to achieve fluorogenic labeling of target molecules remains challenging. It has been adapted only in rare instances for fluorogenic labeling; however, even in such cases, it employs a sluggish Staudinger ligation reaction of phosphene probe.^43^ Herein, we demonstrate a fast and readily generalizable supramolecular ‘catch-and-release’ strategy that can easily integrate any highly optimized dye-quencher pairs for rapid fluorogenic labeling of biological targets. In addition, capitalizing on the unique/orthogonal reactivity profile of this supramolecular motif, we bolster efforts to track multiple biomolecule targets simultaneously via multiplexed bioorthogonal fluorogenic labeling.

**Scheme 2.**
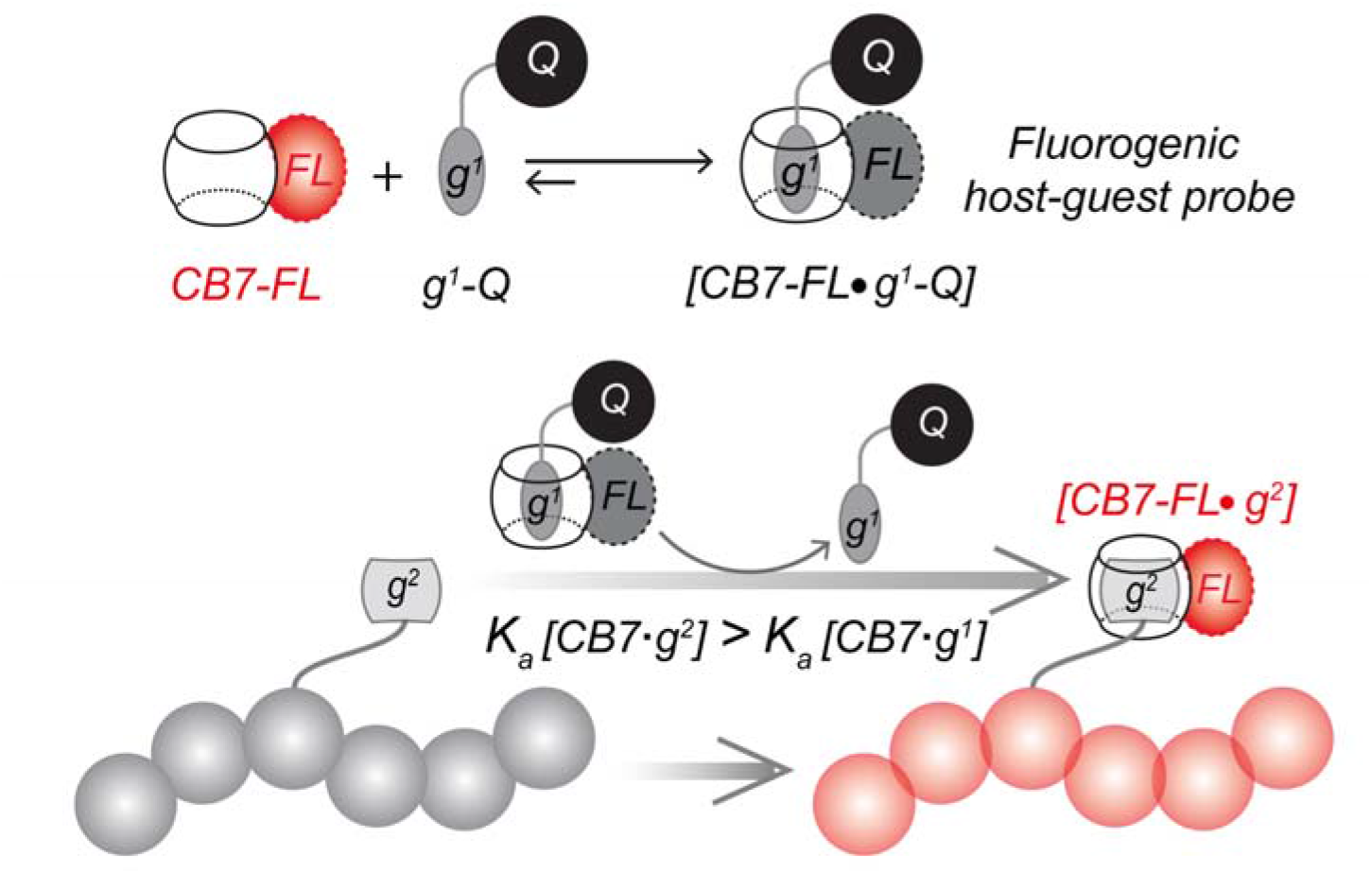
Schematic showing the working principle of supramolecular ‘catch-and-release’ strategy for bioorthogonal fluorogenic imaging. Guest displacement reaction between the quenched CB7-FL·g^1^-Q host-guest complex and a high-affinity guest g^2^ eliminates the quenching effect, restoring the latent fluorescence and forming a fluorescently active reporter complex with the target biomolecule.

Guest displacement reaction between the quenched CB7-FL·g^1^-Q host-guest complex and a high-affinity guest g^2^ eliminates the quenching effect, restoring the latent fluorescence and forming a fluorescently active reporter complex with the target biomolecule. Besides covalent chemistry, an alternative approach towards target labeling is to leverage synthetic host-guest assembly. Most recently, non-covalent host-guest binding pairs based on macrocyclic CB7 host have emerged as a promising biorthogonal imaging tool for visualizing biomolecules both in cells and *in vivo*, often terms as “non-covalent click chemistry”.^44–49^ This is largely fueled by the exceptional molecular recognition property of CB7 with an ability to form ultrastable and highly chemoselective 1:1 host-guest complexes in biological complexities.^50–60^ In addition, their high association kinetics, small size, chemical tractability, and robust chemical structures further boost their potential for advanced imaging applications. Importantly, with the recent demonstrations of the enzymatic,^44^ metabolic,^61^ and genetic^62^ incorporation ability of host-guest elements into biomolecular structures, it is now positioned to offer a powerful alternative to covalent reactions with many complementary benefits. Herein, building on the foundation of the CB7-based host-guest chemistry, we now demonstrate an easily generalizable strategy for designing non-covalent fluorogenic probes across the visible spectrum for bioorthogonal imaging. We designed this strategy, named as supramolecular ‘catch-and-release’ strategy, by programming guest displacement reaction in the synthetic supramolecular host-guest system based on CB7. Scheme 2 illustrates the intrinsic design principle for generating the supramolecular fluorogenic response based on fluorophore-conjugated CB7 reporter probes (CB7-FLs). To suppress the fluorescence of the reporter probe, CB7-FLs are complexed with complimentary guest-conjugated dark quenchers (g1-Qs) via host-guest recognition. Our fluorogenic labeling scheme strategically exploits a guest displacement reaction between the quenched CB7-FL·g1-Q host-guest complex and a high-affinity guest g2 that is directly attached to the target of interest. Upon incubation with the quenched CB7-FL·g1-Q complex, the high-affinity guest g2 binds to the CB7-FL, forming a more thermodynamically stable CB7-FL·g2 complex and displaces the g1-Q out of the CB7 cavity. These binding/association and release events (a outcome similar to the covalent ‘click-to-release’^63^) result in a discrete “turning on” of the CB7-FL fluorescence and the formation of a fluorescently active reporting complex, enabling background-free target visualization under no-wash conditions. To date, CB7-FLs were employed either directly for imaging experiments,^44, 45, 47, 64^ or Forster Resonance Energy Transfer (FRET) pair was developed for monitoring biological processes;^65, 66^ however, to the best of our knowledge, no fluorogenic labeling where fluorescence activation occurs upon specific target engagement has not been demonstrated in cells or tissue. Our current strategy significantly expands the scope of non-covalent chemistry in bioorthogonal labeling, producing a so-called fluorogenic effect that can substantially improve the signal-to-background ratio in fluorescence microscopy and allow no-wash imaging experiments.

## Results and Discussion

The success of this supramolecular fluorogenic imaging strategy was critically dependent on the integration of a suitable host-guest pair that permits selective monovalent complexation (1:1) under physiologically relevant conditions. For this purpose, we relied on CB7 as it stands apart from other synthetic host molecules because of its most striking ability to display exceptionally strong monovalent host-guest molecular recognition (*K_a_*up to 10^17^ M^−1^) towards specific and bioorthogonal guest molecules in biological medium.^50–60, 67, 68^ In addition, depending on the structure of the guest molecules, the affinities can also be tailored from 10^6^ M^−1^ to 10^18^ M^−1^, thereby not only permitting selective complex formation at a biologically viable concentration (typically below μM) but also allowing us to program displacement reactions based on the affinity gradient of various CB7-guest complexes. Specifically, for the design of our displacement-based fluorogenic probes, we choose two guests: *p*-xylylenediamine (XYL, *K_a_*∼10^8^ M^−1^) and 1-adamantylamine (ADA, *K_a_*∼10^14^ M^−1^). ^53^ Given their considerable binding energy difference, we hypothesized that the change in the free energy resulting from the XYL to ADA exchange in the CB7 cavity would be able to drive the guest displacement reaction forward (Figure 1a). To generate fluorogenic probes across the visible spectrum, CB7 is directly attached with a library of well-established fluorophores (CB7-FLs), spanning emissions from 450 nm to 675 nm (Figure 1b). Matching with the fluorophore’s emission, XYL moiety is armed with appropriate dark quenchers (XYL-Qs) to generate quenched host-guest probes (Figure 1c). In addition to Forster Resonance Energy Transfer (FRET) based quenching, these highly optimized quenchers can take advantage of the static quenching, thereby providing excellent fluorescence quenching without the residual background signal. To synthesize CB7-FLs, we first used a photochemical derivatization method, described by Bardelang and Ouari, to synthesize mono-hydroxylated CB7 (CB7(OH)_1_).^69–71^ CB7(OH)_1_ was then converted in two steps to amine derivative (CB7-(NH_2_)_1_), which was subsequently attached to the fluorophores via amide linkage (Figure 1d).^72^ On the other hand, XYL-Qs were synthesized from a nitrobenzyl-protected XYL derivative (Figure 1e). To decorate the protein/biomolecule of interest with high-affinity ADA guest, we prepared two types of ADA conjugated targeting agents (Figure 1f): 1) ADA conjugated antibody and 2) ADA conjugated small molecule binders, phalloidin (for actin targeting) and taxol (for microtubule targeting).

**Figure 1.**
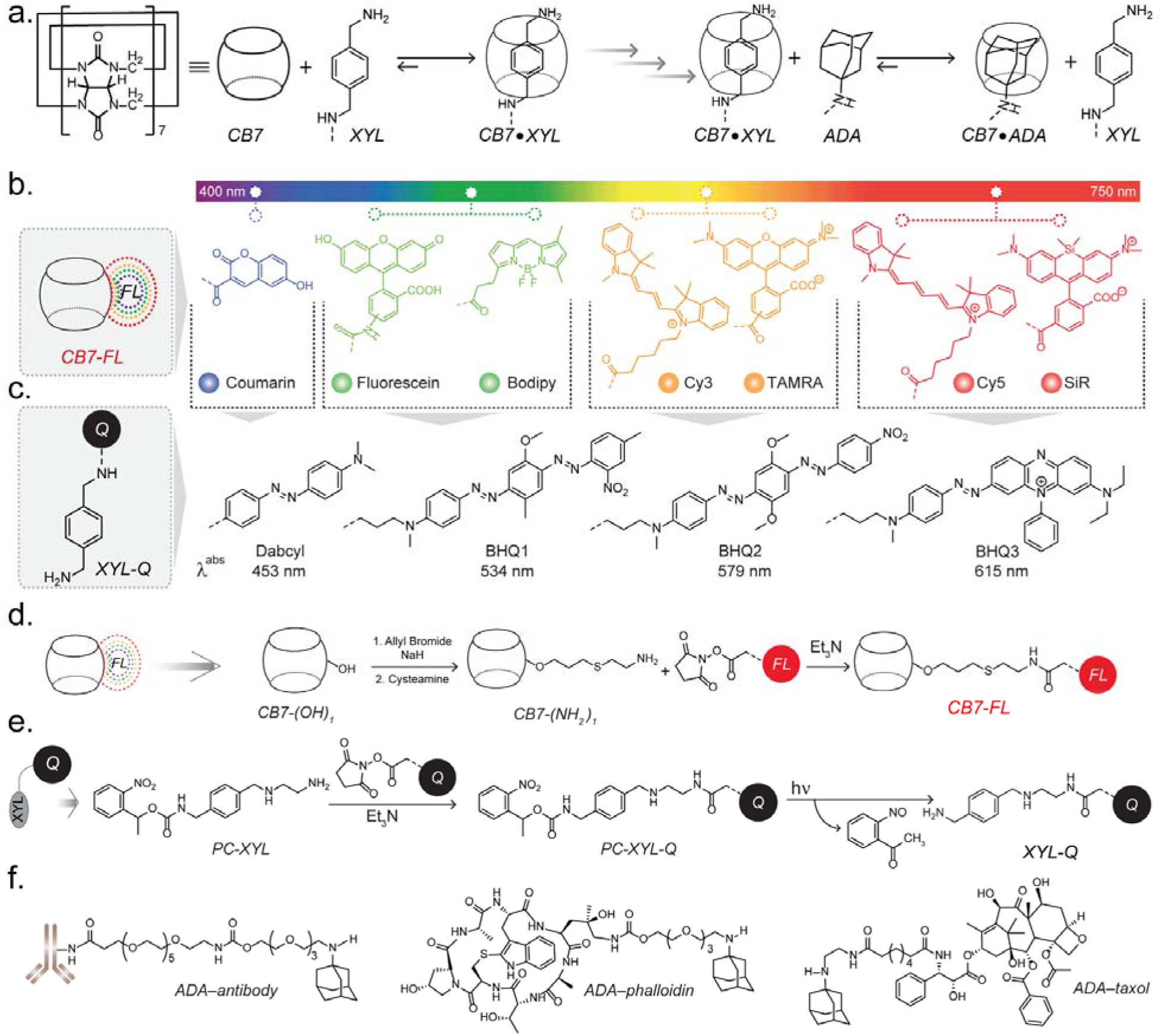
Molecular design for implementing host-guest-based fluorogenic imaging. (a) Molecular structures of host (CB7) and guests (XYL and ADA). (b) Molecular structures of CB7 conjugated fluorophores (CB7-FLs) and (c) XYL conjugated quenchers (XYL-Qs), which are converted to fluorogenic probes. XYL-Qs are used to complex with the spectrally matching CB-FLs to generate fluorogenic probes. (d) Synthetic scheme for generating CB7-FLs. (e) Functionalization scheme of XYL with quenchers. (f) ADA conjugated targeting agents used in the fluorogenic imaging study for specific targeting of biomolecules in cells and tissue.

We first studied the quenching of CB7-FLs by titrating 1 μM solution of CB7-FLs with XYL-Qs. Seven different CB7-FLs were assayed against their spectrally matching XYL-Q molecules (Figure 2a). Notably, all the CB7-FLs exhibited excellent quenching upon the addition of equimolar or slightly higher stoichiometry of respective quencher conjugated XYL guests. The slight variation can be attributed to the structural diversity of the fluorophores, which can have a moderate influence on the CB7·XYL binding affinity. To establish the role of host-guest recognition in the observed fluorescence quenching, we titrated CB7-FLs with a control quencher derivative (EtA-Q) that does not contain any recognition motif for CB7. As shown in Supporting Figure S1, we did not observe any significant quenching upon equivalent addition of EtA-Qs, indicating that the recognition-mediated CB7·XYL complexation is the critical mechanism that brings the fluorophore near to the quencher moiety for effective suppression of fluorescence. Additionally, the formation of a 1:1 complex between CB7-FL and XYL-Q was confirmed via MALDI-MS analysis (Supporting Figure S2), where monovalent complexes are detected in the mass signature. Next, we evaluated the fluorescence activation characteristics of the quenched CB7-FL·XYL-Q complex upon applying a relatively higher binding affinity ADA guest (Figure 1b). The introduction of ADA into a solution of the quenched complex resulted in an immediate increase in fluorescence intensity, a consequence that can be directly attributed to the displacement of XYL-Q by ADA from the CB7 cavity (Figure 1b). MALDI-MS analysis also supported the formation of the CB7-FL·ADA complex upon addition of ADA to the quenched complex (Supporting Figure S3). We also observed a stepwise fluorescence recovery from the quenched complex upon addition of sub-stoichiometric equivalent of ADA, indicating a free energy-driven displacement reaction rather than being a concentration-driven one. Overall, these spectroscopic studies clearly demonstrate a fluorogenic nature of the host-guest complex, which is efficiently triggered by the displacement reaction with a high-affinity bioorthogonal guest molecule.

**Figure 2.**
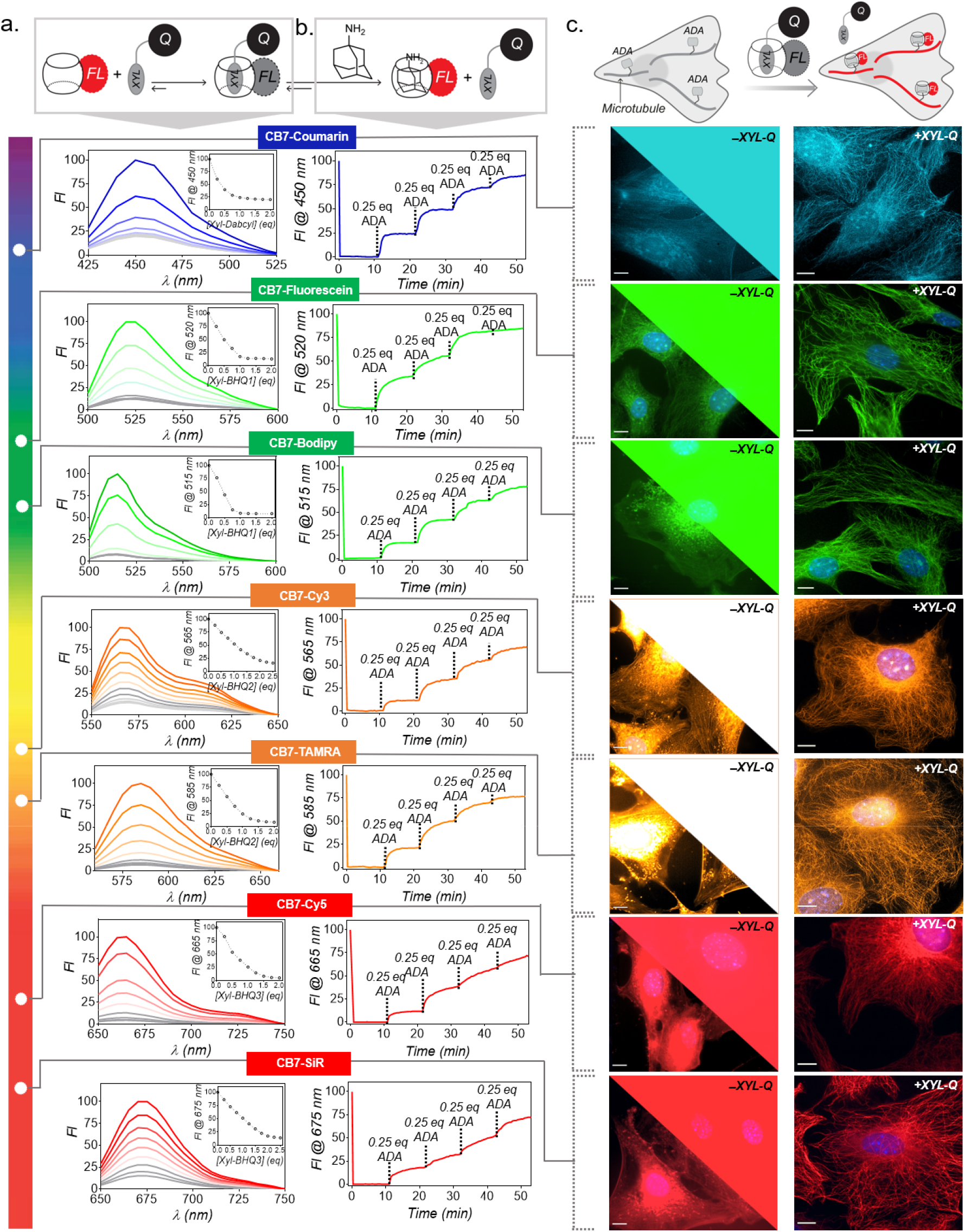
Characterization of guest displacement reaction and demonstration of fluorogenic imaging in the fixed cells. (a) Fluorescence titration of CB7-FL with XYL-Q shows fluorescence quenching upon host-guest complex formation. All the CB7-FLs exhibited excellent quenching upon the addition of respective XYL-Q. (b) Fluorescence recovery upon addition of ADA to the quenched probes. Stepwise addition of the sub-stoichiometric equivalent of ADA showed an efficient fluorescence recovery from the quenched probes. (c) Fluorogenic imaging of microtubules from fixed cells using ADA-Abs and CB7-FL·XYL-Q quenched probes. Right panel: High-contrast microtubules images were observed from fluorogenic probes without removing excess unbound probes. Left panel (top right corner + bottom left corner): Control studies with only CB7-FLs (without XYL-Qs) did not lead to an appreciable microtubule visualization due to high background fluorescence from the unbound fluorophores. The top right corner shows images that are represented with the same brightness contrast as that of the fluorogenic images. The bottom left corner presents images with adjusted brightness contrast, best suited to visualize signals from cells. Scale bar: 10 μm (c).

We next evaluated the ability of this non-covalent ‘catch-and-release’ strategy for fluorogenic imaging of a specific biological target in cellular settings. Before target imaging, we first probed the stability of the quenched CB7-FL·XYL-Q complex in the cellular environment. For this purpose, quenched CB7-TAMRA·XYL-BHQ2 complex was incubated with the mouse embryonic fibroblast (MEF) cells and time-lapse fluorescence microscopy images were recorded from the surrounding media to observe any cell mediated dissociation of the quenched complex. Quantification of these time-lapse microscopy images showed a negligible increase in fluorescence intensity over time (Supporting Figure S4), indicating minimal non-specific activation/dissociation of CB7-FL·XYL-Q quenched complex by native cellular components. Additionally, no appreciable fluorescence signal or non-specific staining was observed from the cells upon incubation with the CB7-FL·XYL-Q quenched complex (Supporting Figure S5). We next investigated whether the higher affinity ADA guest directed to a biological target molecule could be utilized for target-specific fluorogenic activation and visualization of the cellular entities under no-wash imaging conditions. To evaluate our hypothesis, we conjugated ADA with antibodies (ADA-Ab) and targeted microtubules in MEF cells. After immunolabeling using ADA-Ab, MEF cells were incubated with the quenched CB7-FL·XYL-Q probes and subsequently imaged without performing additional washing steps required to remove unbound probes. A set of traditional experiments were also conducted where immunolabelled cells were stained with only CB7-FLs under identical conditions to compare the effect of unbound fluorescent probes in target visualization. In all cases, cells treated with the fluorogenic probes, consisting of complexed CB7-FL·XYL-Q probe, showed high-contrast staining of microtubules, whereas cells treated with only CB7-FL under identical conditions only led to barely visible microtubule structures (Figure 2c). Notably, with the host-guest fluorogenic probes, microtubules were clearly visible even under the widefield epi-fluorescence mode of microscopy, where the presence of any background fluorescence is known to significantly impact the image quality (Figure 2c). This indicates suppressed fluorescence from the unbound host-guest probes played a critical role in high-contrast target visualization. Notably, specific, and background-free staining of microtubules were consistently observed from all the spectrally different host-guest quenched probes, demonstrating the advantage of this easily generalizable non-covalent ‘catch-and-release’ based strategy for rapid generation of multicolor fluorogenic probes. A quantitative estimation from fluorescence intensity profiling also indicated dramatically enhanced signal-to-noise from the CB7-FL·XYL-Q probes compared to CB7-FL alone (Supporting Figure S7). The specificity of the host-guest labeling approach was also established via co-localization experiment with a microtubule targeted direct-fluorophore antibody conjugate (Supporting Figure S12). In order to gain a quantitative view of the fluorogenic labeling kinetics, time-lapse imaging was performed on ADA-targeted cells using a quenched CB7-TAMRA·XYL-BHQ2 probe (Supporting Figure S6). Intensity profile over time showed completion of the fluorogenic labeling within minutes, highlighting the suitability of these probes for a fast pulse-chase labeling experiment or labeling under high flow *in vivo* conditions. Fluorescence intensity was also found to be stable over time after reaching saturation which could be potentially useful for fluorescence tracking experiments. Overall, these results clearly establish the rapid fluorogenic imaging capability of the host-guest probe across the visible spectrum with the advantages of no-wash imaging and substantially improved signal-to-noise ratio for fluorescence microscopy.

In addition to the suppressed background from unbound probes, we probed the reduction of non-specific signals by imaging microtubules labeled cells after removing excess unbound probes. Given that host-guest quenched probe exhibits fluorescence signal only upon encountering ADA counterpart, nonspecifically bound probes are expected to remain ‘invisible’ due to their non-fluorescent nature. The fluorogenic host-guest probe (+XYL-Q) exhibited microtubule visualization with a minimal off-target fluorescence signal, whereas CB7-FLs, when used alone (-XYL-Q), showed strong off-target binding even after repetitive washing (Figure 3 and Supporting Figure S8). This contrasting effect additionally highlight the advantage of our fluorogenic probes over direct use CB7-FLs in bioorthogonal imaging.

**Figure 3.**
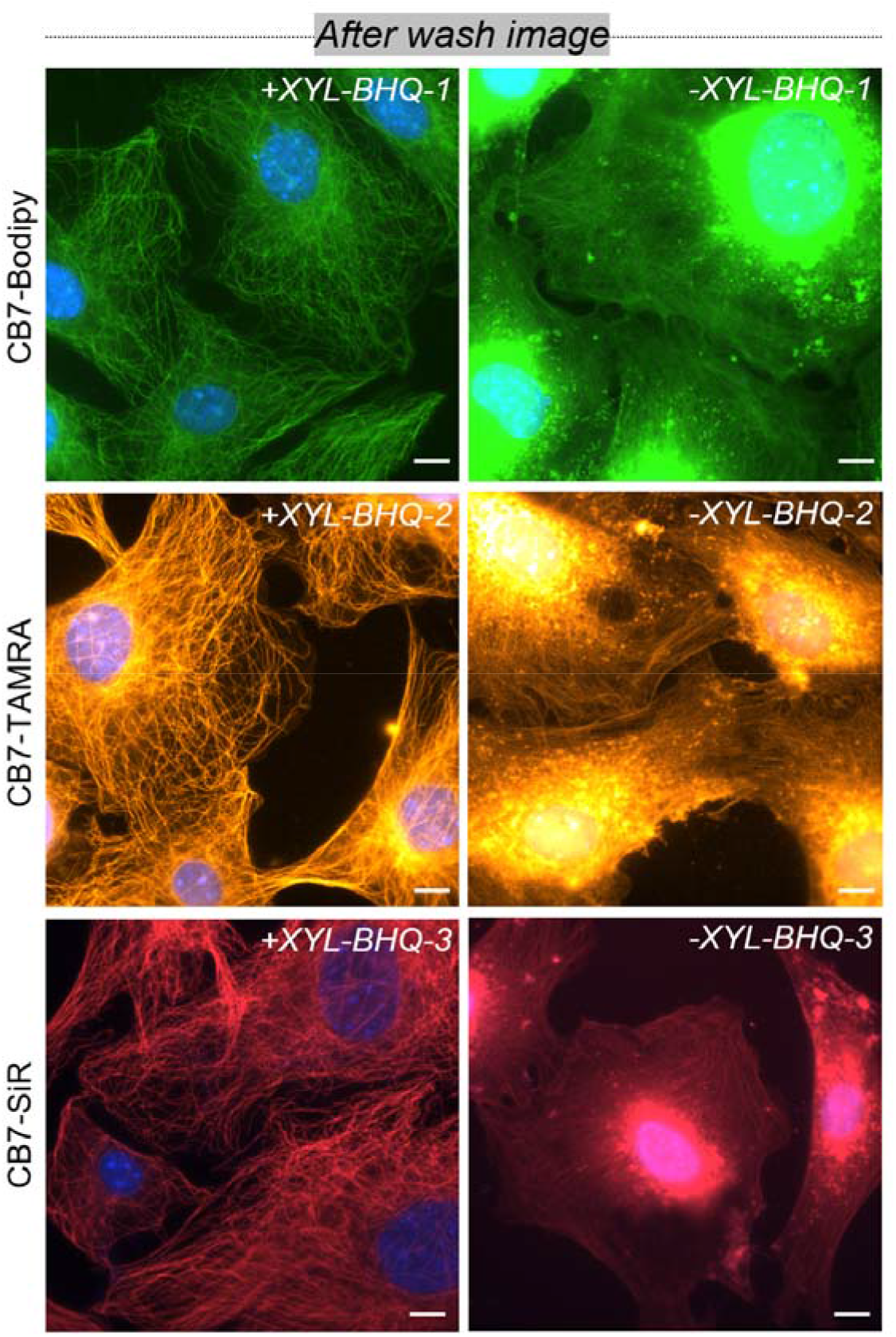
After wash images from labeling with fluorogenic CB7 probes (+XYL-Q) and CB-FLs alone (-XYL-Q). A dramatically reduced off-target signal from the host-guest quenched probes (+XYL-Q) as compared to the CB7-FL (-XYL-Q) alone was observed due to the conditionally activable nature of the quenched probes. Scale bar: 10 μm.

Compared to cells cultured on coverslips, labeling and imaging specific targets in tissue samples usually harbor a more significant challenge. Tissues possess a complex architecture, and with their inherently slow diffusion kinetics, it is even more challenging to rapidly remove unbound fluorophores or eliminate nonspecifically bound probes. We tested whether our non-covalent ‘catch-and-release’ strategy could be applicable for fluorogenic imaging in tissue samples to tackle these challenges. To test this, we attempted to label actin filaments in the thoracic muscle of *Drosophila melanogaster* using the fluorogenic probes. A small molecule binder of actin, namely, phalloidin, is attached with the ADA guest molecule and used for targeting the actin structure (Figure 4a). First, the target specificity of the phalloidin molecule after attachment with ADA guest was tested in cells, where actin filaments were simultaneously visualized with a traditional alexa488-phalloidin stain along with the ADA-phalloidin probe. Excellent colocalization of the two probes indicated the specificity of the ADA-phalloidin towards the intended actin target (Figure 4a). Next, we used ADA-phalloidin to target actin structures in the thoracic muscle of Drosophila melanogaster for fluorogenic imaging. Incubation of phalloidin-ADA conjugates with the tissue sample, and the subsequent addition of CB7-FL·XYL-Q fluorogenic probe resulted in specific and distinct visualization of a two-dimensional (2D) actin pattern in the muscle sample (Figure 4b). A repeating band patterned fluorescence, resembling intrinsic spatial organization actin in thoracic muscle, was prominent from the epi-fluorescence microscopy images. Notably, the visualization of actin patterns from the tissue section remained impressive when tested with the range of fluorogenic host-guest probes. We also performed 3D imaging of the actin structure from the tissue samples to understand fluorogenic imaging capability inside a thick tissue sample. The 3D distribution of actin patterns in the ovary of *Drosophila melanogaster* was clearly visualized with minimal background over ∼80 μm axial direction (Supporting Figure S11). These results highlight the applicability of the host-guest fluorogenic probes in labeling and imaging complex tissue samples.

**Figure 4.**
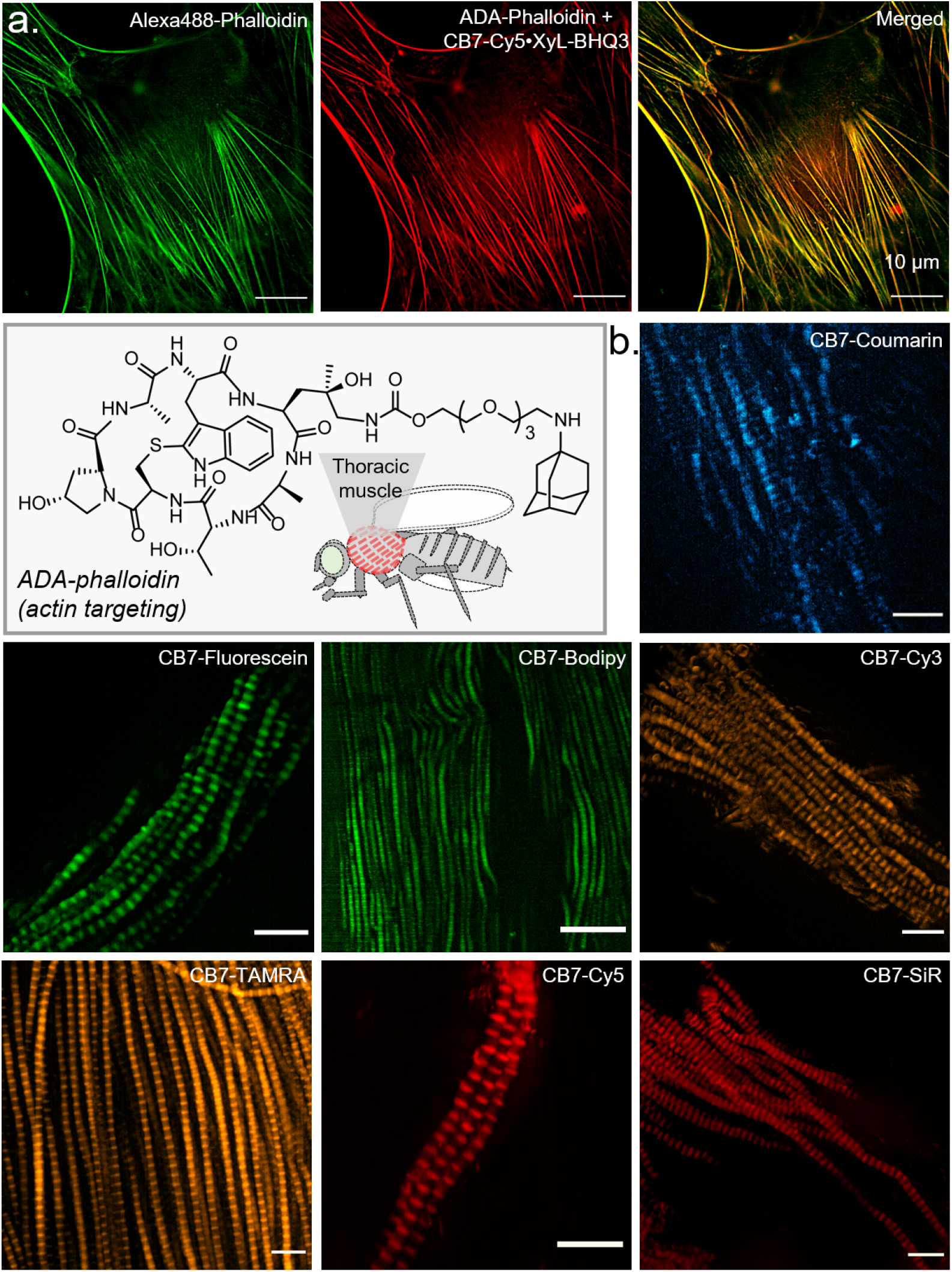
Host-guest fluorogenic imaging of the actin structures in fixed MEF cells and fixed thoracic muscle tissue of Drosophila melanogaster. (a) Two color imaging of actin, comparing guest-modified targeting ligands to direct fluorophore-conjugated imaging agent. Additionally, the molecular structure of ADA-phalloidin that is used for targeting actin in muscle tissue is shown. (b) Epi-fluorescence images of the muscle tissue after incubation with spectrally diverse fluorogenic probes. We observed repeating band patterned actin distribution using host-guest-based fluorogenic imaging strategy. Scale bar: 10 µm (a) and (b).

We subsequently employed our non-covalent ‘catch-and-release’ strategy for fluorogenic imaging of biological targets in living specimens. Towards this goal, we first intended to image the overexpression of epidermal growth factor receptor (EGFR) on the live A431 cell membrane (Figure 5a). After targeting EGFR with ADA-Ab, we incubated the A431 cells with the fluorogenic probes for imaging. Live A431 cells imaged without the washing, or excess probe removal step showed a strong fluorescence signal emanating from the membrane, indicating fluorogenic labeling of the EGFR on the live-cell surface (Figure 5b and Supporting Figure S9). Control experiments that are performed without adding ADA-Ab to the A431 cells or with an EGFR negative cell line (3T3 cells) showed negligible membrane fluorescence, indicating specificity of the host-guest probes for labeling targets in the living system (Supporting Figure S10). Next, we utilized our non-covalent ‘catch-and-release’ strategy for fluorogenic imaging of biological targets inside living cells. We used a small molecule-based microtubule binder, docetaxel, to specifically target polymeric microtubules in live cells. To afford fluorogenic microtubule imaging, we prepared an ADA derivative of docetaxel (ADA-taxol), where ADA is conjugated with docetaxel *via* a C6–linker (Figure 5c). First, the specificity of the docetaxel towards microtubule after derivatization with ADA was tested by an *in vitro* experiment. In this experiment with *in vitro* polymerized microtubules, ADA-taxol·CB7-Bodipy complex was found to be colocalized with the directly alexa568 labeled microtubules, confirming the target specificity of the ADA-taxol (Figure 5d). However, poor cell membrane permeability of these dyes or the dye-quencher pairs limits their application in efficient target labeling inside a live cell. As a result, to expand this supramolecular fluorogenic strategy for live intracellular target imaging, we considered evaluating a new system where an efficient delivery vector could serve the dual purpose of intracellular transporter of dyes as well as act as a guest quencher. For this purpose, we explored the use of gold nanoparticles (AuNPs), which are known to be highly efficient cytosolic delivery platforms with the ability to quench proximal dye fluorescence across a broad spectral range.^73–76^ In order to transform the AuNP into a guest quencher, we surface-functionalized it with XYL guest conjugated thiolated ligand and promoted the formation of quenched CB7-FL·XYL complex at the AuNP surface (Figure 5). Fluorescence quenching of the CB7-FL dyes by XYL decorated AuNPs (XYL-AuNPs) was investigated by titrating 1 μM solution of CB7-TAMRA (CB7-FL) with an increasing concentration of XYL-AuNP. Notably, we observed an excellent quenching of CB7-TAMRA fluorescence upon the addition of 0.2 equivalent of XYL-AuNP, arising from the multivalent nature of the guest XYL-AuNP coupled with its efficient quenching ability (Figure 5e). Furthermore, to establish the role of host-guest recognition in the observed fluorescence quenching, we titrated TAMRA-EtA, which does not have any CB7 recognition motif with XYL-AuNP. In this case, we did not observe any significant quenching upon the addition of 0.2 equivalent of XYL-AuNP (Supporting Figure S13 a), indicating that the recognition-mediated CB7·XYL complexation plays a crucial role in bringing the fluorophore close to the AuNP surface for effective quenching of fluorescence. Also, we examined the fluorescence activation characteristics of the quenched CB7-TAMRA·XYL-AuNP complex upon applying a relatively higher binding affinity ADA guest (Figure 5f). The introduction of ADA to the quenched complex resulted in an immediate recovery in fluorescence intensity, a consequence that can be directly attributed to the displacement of CB7-FL dye by ADA from the XYL-AuNP surface (Figure 5f). Overall, these spectroscopic studies demonstrate the fluorogenic nature of the AuNP host-guest complex, which is efficiently triggered by the displacement reaction with a high-affinity bioorthogonal guest molecule. Next, we used this quenched CB7-TAMRA·XYL-AuNP complex to image the microtubule network inside live HeLa cells (Figure 5g). After targeting the microtubule with ADA-taxol, we incubated the HeLa cells with the CB7-TAMRA·XYL-AuNP fluorogenic probe for target imaging. Live HeLa cells imaged without the inclusion of a washing step showed strong fluorescence signals originating from the microtubule filaments, indicating fluorogenic labeling of the microtubules inside the live cell (Figure 5g and Supporting Figure S13c). High-contrast microtubule images with minimal background fluorescence indicate that XYL-AuNP not only efficiently transported CB7-FLs inside the cell but also kept the unbound FLs in a quenched state in the intracellular environment. In addition, the control experiment that was performed without adding ADA-taxol to the HeLa cells showed negligible microtubule fluorescence, indicating the specificity of the host-guest probes for intra-cellular labeling targets in the living system (Supporting Figure S13 b). Next, we evaluated whether the host-guest fluorogenic probes could be translated into even more complex settings, like live tissue imaging. In this regard, we chose to image the distribution of microtubules in intestinal tissue from *Drosophila melanogaster*. ADA-taxol was used to stain microtubules in live intestinal tissue. Subsequently, CB7-FL·XYL-Q fluorogenic probe in Schneider’s medium was incubated with the ADA-labeled tissues, and imaging was performed via super-resolution structured illumination microscopy (SIM). The SIM images that are acquired with the range of host-guest fluorogenic probes clearly showed the distribution of microtubules in the live intestine tissue (Figure 5h). Notably, all the spectrally different fluorogenic probes performed efficiently to label and visualize the distribution of microtubules from the live intestine tissues. Overall, these results prove the applicability of our non-covalent ‘catch-and-release’ strategy-based fluorogenic probes for imaging complex living specimens.

**Figure 5.**
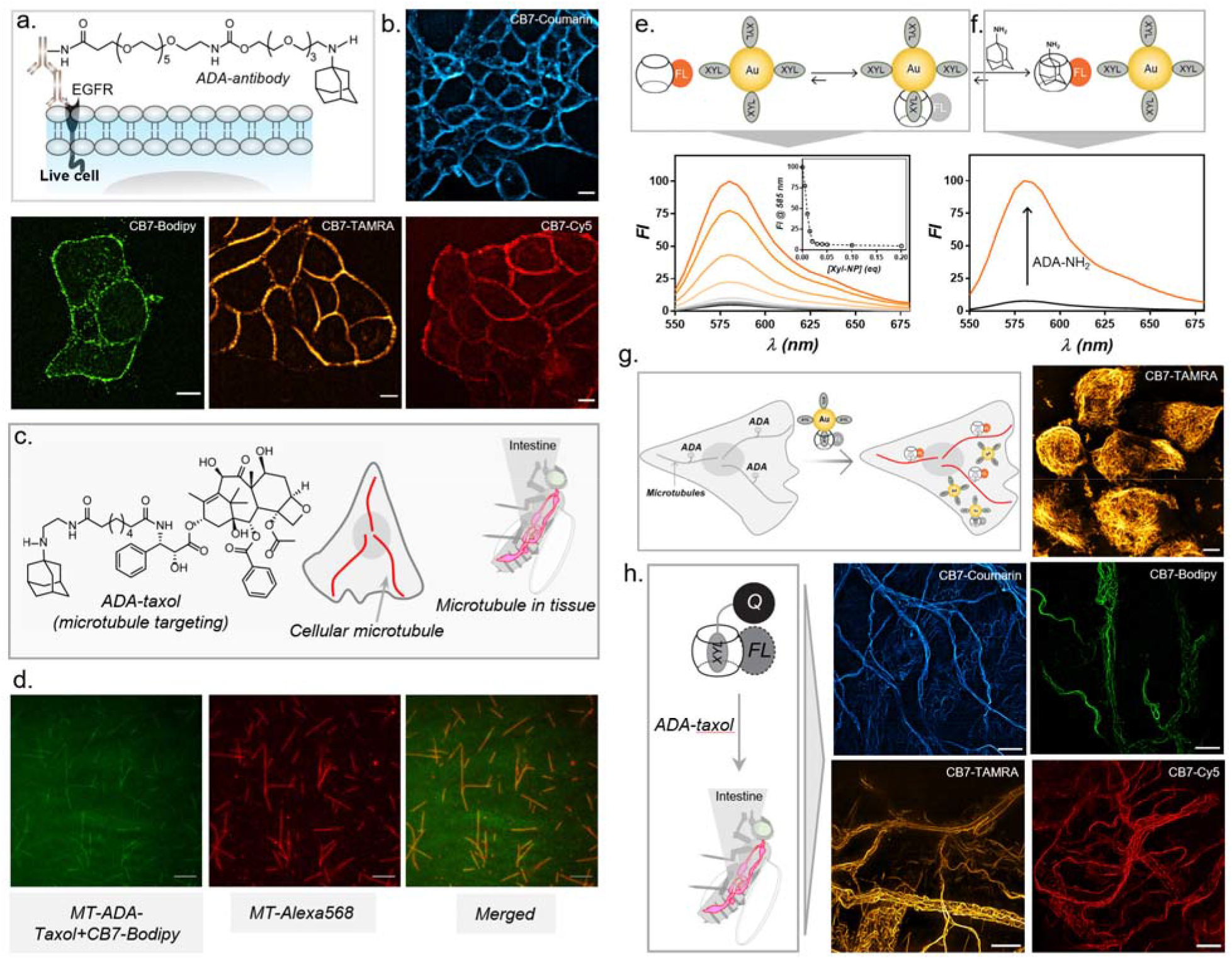
Host-guest fluorogenic imaging in live cells and tissues (a) Targeting strategy for EGFR on the live cell membrane. (b) Fluorescence images from the EGFR overexpressing live A431 cells show an intense membrane staining, indicating fluorogenic labeling of the EGFR on the live-cell surface. (c) Molecular structure of ADA conjugated taxol derivative that is used for the specific targeting of microtubules in the live cells and live intestine tissue. (d) Two-color imaging of in-vitro microtubules, comparing guest-modified targeting ligand (ADA-taxol) to direct fluorophore-conjugated microtubules. (e) Fluorescence titration of CB7-TAMRA with XYL-AuNP shows fluorescence quenching upon host-guest complex formation. (f) Fluorescence recovery upon addition of ADA to the quenched probes. (g) Fluorogenic imaging of microtubules in live Hela cells using ADA-Taxol and CB7-FL·XYL-AuNP quenched probes. (h) No-wash, fluorescence SIM images from the live intestine sample show the microtubule distribution within the tissue specimen. Scale bar: 10 µm (b, g, and h), 5 µm (d).

Simultaneous imaging of multiple biomolecules is an important aspect of modern biological investigations, requiring fluorogenic pairs with orthogonal reactivity. The development of our host-guest-based quenched probe presents a new opportunity for multiplexed fluorogenic labeling, as it can be easily paired with the covalently clickable fluorogenic probe. In addition to its orthogonal reactivity, the flexible choice of fluorophores that can be used in the host-guest-based fluorogenic mechanism makes it an ideal choice for multicolor bioorthogonal fluorogenic imaging. To demonstrate this, we used CB7-TAMRA·XYL-BHQ2 based non-covalent fluorogenic probe and combined it with a tetrazine (Tz)-Bodipy-based fluorogenic probe for multiplexed imaging (Figure 6a).^39^ We first aimed to simultaneously image the distribution of microtubule and actin in cells, which were accordingly targeted using ADA-Ab and TCO-phalloidin, respectively. After incubating with a mixture of quenched probes, we acquired dual-color SIM images from the cells. As demonstrated in Figure 6b, the SIM images clearly revealed specific staining of the microtubule network in the TAMRA channel via CB7·ADA displacement, whereas actin filaments showed a strong signal in the bodipy channel via Tz-TCO activation.

**Figure 6.**
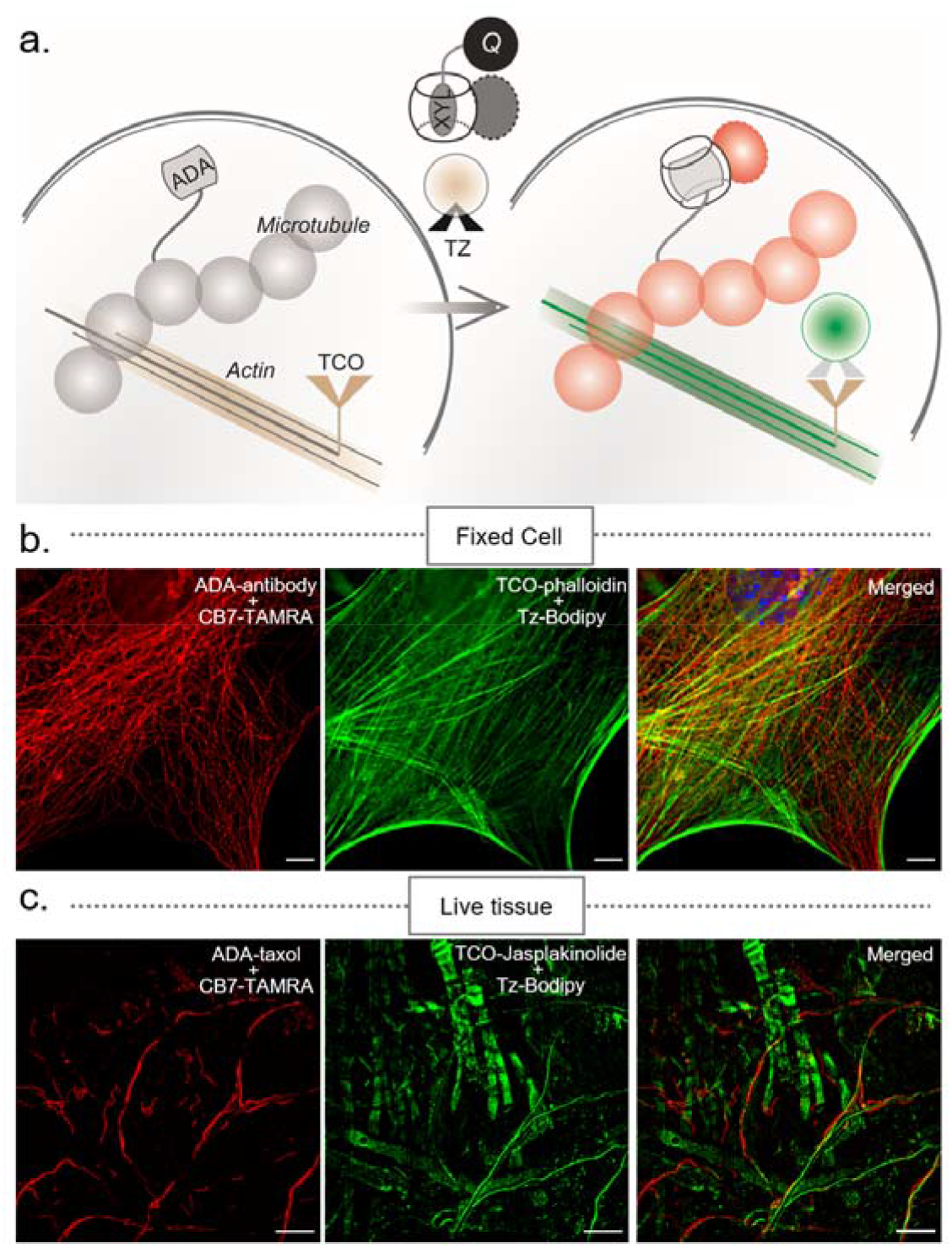
Fluorogenic multiplexed imaging in fixed cells and live tissues using orthogonally reactive fluorogenic probes. (a) Strategy for multiplexed fluorogenic imaging by combining the host-guest fluorogenic system with a covalently clickable (TCO–Tz) fluorogenic probe. (b-c) Multiplexed fluorogenic imaging of microtubules and actins in fixed cells and live tissue using orthogonally reactive CB7-TAMRA·XYL-BHQ2 and Tz-Bodipy probes. Scale bar: 10 µm (b–c).

This suggests that orthogonal reactivity of CB7 and Tz based fluorogenic probes can be easily adopted for multicolor imaging experiments without using a clearing step. We further performed dual-color fluorogenic imaging of microtubule and actin in live intestine tissue of *Drosophila melanogaster*. Microtubules and actin were targeted using living system compatible small molecule-based targeting agents, docetaxel-ADA and jasplakinolide-TCO conjugates. SIM microscopy images acquired after incubation with the pair of orthogonally reactive fluorogenic probes confirmed distinct staining of the microtubule and actin in live intestine tissue (Figure 6c), highlighting the importance of the host-guest fluorogenic probe in a multiplexed assay.

## Conclusion

In conclusion, we introduced a distinct supramolecular ‘catch-and-release’ strategy based on CB7 host-guest interaction for generating highly efficient bioorthogonal fluorogenic probes. This strategy is highly generalizable and can be easily extended to any readily available fluorophore scaffold without any further core structural alternations, providing straightforward synthesis and appealing optical flexibility for imaging applications. Utilizing this supramolecular ‘catch-and-release’ strategy, we transformed a library of highly optimized dye molecules, spanning the entire visible-light spectrum, into highly efficient fluorogenic probes, validating our approach’s generalizability and robust mechanism. We demonstrated no-wash fluorogenic imaging of diverse target molecules in the biological complexities of live cells and tissue sections, where high contrast images of the target molecules were achieved with minimal background fluorescence and negligible non-specific signal. Further design optimization will focus on developing fluorogenic single-component CB7-dye conjugate. Some attractive CB7-dye conjugates were reported for single component analyte sensing;^77, 78^ however, critical design changes would be necessary to reverse their ON-to-OFF response to OFF-to-ON response for fluorogenic imaging applications. Nevertheless, the fast and catalyst-free reactivity, straightforward synthesis, mutual orthogonality to the preexisting bioorthogonal reactions, and appealing optical flexibility of our current host-guest fluorogenic probes should synergize with emerging microscopic methods to open up new opportunities in multiplexed investigations *in vitro*, *in vivo*, and in diagnostic settings. Moreover, because of its highly flexible choice of fluorophores unmatched by other techniques, the current strategy can easily accommodate already existing highly specialized dye molecules optimized for super-resolution imaging, providing an important addition to the toolkit of fluorogenic nanoscopic imaging. ^2,3, 79, 80^

## Code Availability

Research papers using custom computer code will also be asked to fill out a <u>code and software submission checklist</u> that will be made available to editors and reviewers during manuscript assessment. The aim is to make studies that use such code more reliable by ensuring that all relevant documentation is available and by facilitating testing of software by the reviewers. Further detailed guidance and required documentation at submission and acceptance of the manuscript can be found <u>here</u>.

## ASSOCIATED CONTENT

The Supporting Information is available free of charge.

Synthesis and characterization data of the host-guest probes and targeting ligands. Experimental protocols for imaging, microscopy, analysis and other supporting data (PDF).

## AUTHOR INFORMATION

Corresponding Author Sarit S. Agasti

## Contributions

R.S., A.S., and S.S.A. conceived the study, designed and performed the experiments, analyzed and interpreted the data, and wrote the manuscript. P.K. and N.D.S. prepared cells for the imaging experiment and reviewed the manuscript. M.P. performed a few titration experiments.

S.R. and S.V. prepared the tissue sample for imaging. S.S.A. supervised the overall study.

## Acknowledgements

We acknowledge the support from Dr. Minhaj Sirajuddin lab (inStem) for generous gift of microtubule monomers (native, labelled, and biotinylated). We thank Akshay Saroha for help with the in vitro microtubule experiment and Dr. Jerrin Thomas George for critical comment on the manuscript. We thank Athira MP, K. Palani Ganesh and V.Ramjayakumar for help with the nanoparticle synthesis.

