## Supplementary information for "Supramolecular ‘catch-and-release’ strategy for bioorthogonal fluorogenic imaging across the visible spectrum"

**for**

**Contents**

1. General information………………………………………………………………………………………..S3

2. Synthesis protocol for amine derivatives of *p*-xylenediamine (XYL) and 1-adamantylamine (ADA)………………………………………………………………………………………………………..S5

3. Synthesis protocol for XYL conjugated quenchers (XYL-Q)……………………………………………...S7

4. Synthesis protocol for EtA conjugated quenchers (EtA-Q)……………………………………………….S10

5. Synthesis protocol for tetrazine conjugated BODIPY fluorophore……………………………………….S11

6. Synthesis of NHS ester derivatives of ADA………………………………………………………………S12

7. Conjugation of ADA with the antibody…………………………………………………………………..S12

8. Conjugation of ADA with phalloidin……………………………………………………………………..S13

9. Conjugation of ADA with taxol…………………………………………………………………………..S13

10. Conjugation of TCO with phalloidin…………………………………………………………………….S14

11. Conjugation of TCO with jasplakinolide…………………………………………………………….….S15

12. Synthesis scheme for XYL-AuNP……..…………………………………………………………….….S16

13. Other experimental protocols and supporting results……………………………………………….…..S17

13.1. Protocol for fluorescence quenching study of CB7–FLs with XYL–Qs……………………..S17

13.2. Protocol for fluorogenic response study from CB7–XYL quenched complex……………….S17

13.3. Protocol for MALDI-MS analysis…………………………………………………………….S19

13.4	̶ 13.5. Protocol for fluorescence quenching study of CB7–FL with XYL–AuNP……………..S21

13.6 ̶ 13.16. *In vitro* microtubule labeling, cell culture, tissue dissection, cell/tissue target labeling, and fluorogenic imaging protocols……………................................................................................S22

14. Microscopy set up…………………………………………………………………………………..…..S28

15. Image processing and data analysis……………………………………………………………….……S30

16. HPLC, MALDI-MS, and NMR characterization data………………………………………………….S37

17. References………………………………………………………………………………………………S48

**1. General information**

All the chemicals were purchased from either of the following companies: Sigma Aldrich, Alfa Aesar, Thermo Fischer Scientific, TCI Chemicals, Merck, SD fine chemicals, and Spectrochem, unless mentioned specifically. Amino Phalloidin (Product No. 92-1-10) was purchased from the American Peptide Company. Lysine derivative of jasplakinolide (Boc-Lys-jasplakinolide, SKU – SC005) was purchased from Spirochrome Ltd. Phosphate buffered saline powder pH 7.4 (Product No. P3813-10PAK) was purchased from Sigma-Aldrich. Zeba^TM^ spin desalting columns (Product No. 89883) were purchased from Thermo Fisher Scientific. Antibodies, fluorophores, and quenchers were purchased from commercial sources as listed below. Whenever necessary, solvents were dried by using standard solvent drying methods and then used for reactions. ^1^H NMR spectrum was recorded using Bruker AVANCE III 400 MHz and JEOL Delta 600 MHz instrument, and data analysis was done using Spinworks_4.0 and JEOL delta v5.0.5.1 software. High-Resolution Mass Spectrometry (HRMS) was carried out using Agilent 6538 Ultra High Definition (UHD) Accurate-Mass Q-TOF LC/MS. Liquid chromatography-mass spectrometry (LCMS) experiments were carried out using a Waters Alliance High-Performance Liquid Chromatography (HPLC) system attached a SQD2 mass detector. HPLC purification was carried out using Agilent 1260 infinity quaternary HPLC system equipped with analytical ZORBAX Eclipse plus C18 column (4.6 mm × 100 mm, 3.5 microns) and semipreparative ZORBAX Eclipse plus C18 column (9.4 mm × 250 mm, 5 microns). The solvents used as eluent in HPLC purification were solvent A (water containing 0.1% TFA) and solvent B (acetonitrile containing 0.1% TFA). In general, gradient elution of solvent B in solvent A from 5-100% was used for the purification. Absorbance measurement was carried out to check the concentration of fluorophores in an Eppendorf BioSpectrometer. The microscopic studies were carried out using three optical set-ups: a) A custom-built inverted epi–fluorescence microscope (Olympus) equipped with Cool–LED light source, 2) A Zeiss ELYRA PS1 set up for Structured Illumination Microscopy (SIM), and 3) A Leica SP8 confocal microscope.

**Table S1**: List of primary antibodies used for immunostaining

| **Target** | **Antibody commercial sources** | **Species** |
| --- | --- | --- |
| Microtubule (anti–α tubulin) | Thermo Fischer Scientific (MA1-80017) | Rat |
| Human EGFR (Cetuximab) MAb (Clone Hu1) | R&D Systems, Inc (MAB9577-100) | Human |
| Human EGFR (Cetuximab) | Merck (IL/BIO-000085-FF-373) | Human |

**Table S2:** List of secondary antibodies used for immunostaining

| **Target** | **Host** | **Specification and commercial source** |
| --- | --- | --- |
| Rat | Donkey | Donkey Anti-Rat IgG (H+L)  (min X Bov, Ck, Gt, GP, SyHms, Hrs, Hu, Ms, Rb, Shp Sr Prot)  Jackson ImmunoResearch Laboratories (Cat. No. 712-005-153) |
| Human | Donkey | Donkey Anti-Human IgG (H+L)  (min X Bov, Ck, Gt, GP, Sy Hms, Hrs, Ms, Rb, Rat, Shp Sr Prot)  Jackson ImmunoResearch Laboratories (Cat. No. 709-005-149) |

**Table S3**: List of fluorophores used for the fluorogenic imaging study.

| **Fluorophores** | **Commercial source** |
| --- | --- |
| Coumarin NHS ester | TCI Chemicals (Cat No. S0866) |
| Fluorescein Isothiocyanate | TCI Chemicals (Cat No. F0026) |
| BODIPY NHS ester | TOCRIS Bioscience (Cat No. 5465) |
| TAMRA NHS ester | Sigma Aldrich (Cat No. 53048) |
| Cy3 NHS ester | Lumiprobe (Cat. No. 21020) |
| Cy5 NHS ester | Lumiprobe (Cat. No. 23020) |
| Silicon rhodamine (SiR) NHS ester | Spirochrome (SC003) |
| Alexa488-Phalloidin | Thermo Fisher Scientific (A12379) |
| Alexa568-NHS | Thermo Fisher Scientific (A20103) |

**Table S4**: List of Quenchers used for the fluorogenic imaging study.

| **Quenchers** | **Commercial source** |
| --- | --- |
| Dabcyl NHS ester | Sigma Aldrich (Cat No. 09278) |
| BHQ1 NHS ester | LGC Biosearch Technologies (Cat No. BHQ–1000S) |
| BHQ2 NHS ester | LGC Biosearch Technologies (Cat No. BHQ–2000S) |
| BHQ3 NHS ester | LGC Biosearch Technologies (Cat No. BHQ–3000S) |

**2. Synthesis protocol for amine derivatives of p-xylenediamine (XYL) and 1-adamantylamine (ADA)**

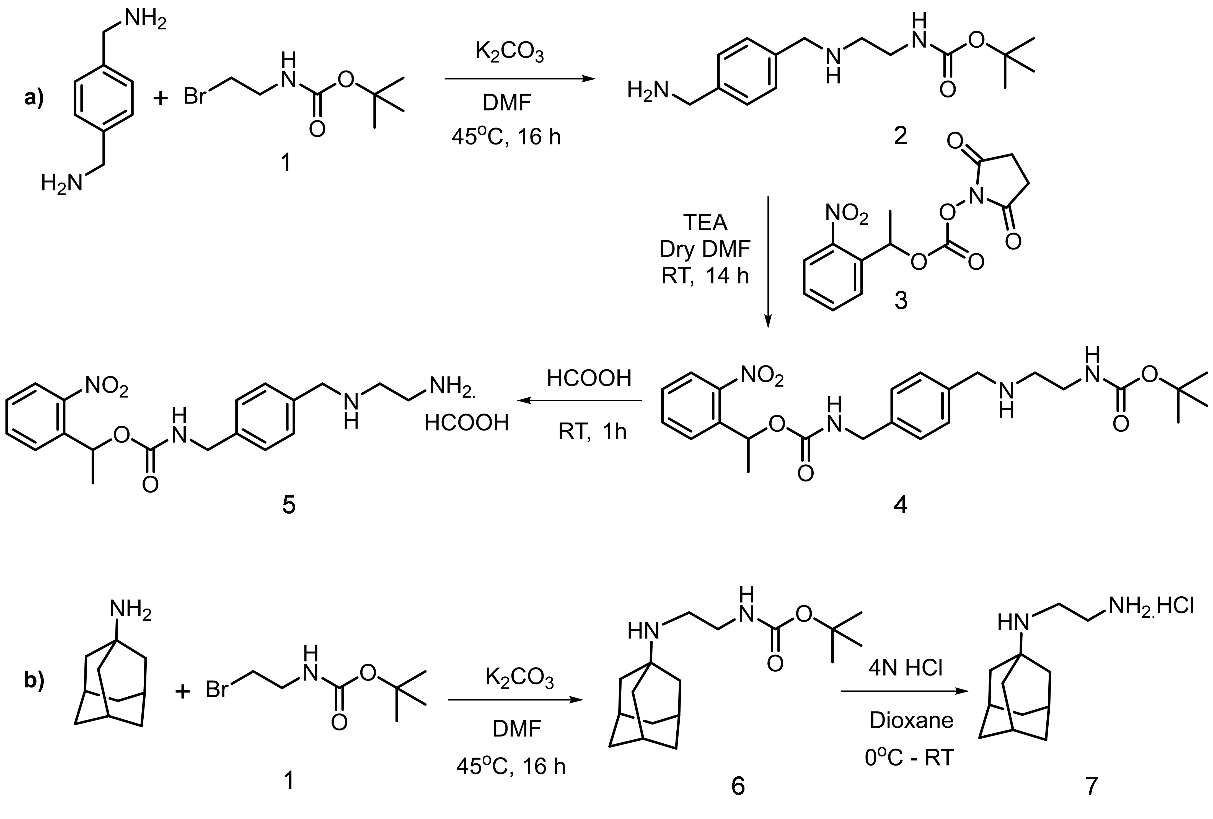

**Scheme S1:** Synthetic scheme for preparing amine derivatives of a) XYL and b) ADA guests.

**2.1. Synthesis of Compound 2**

Compound 1 has been synthesized according to the literature procedure^[1]^. In a 50 ml RB flask, p–xylenediamine (3 g, 22.027 mmol) was dissolved in 15 ml dry DMF followed by the addition of K_2_CO_3_ (1.5 g, 10.85 mmol) to the solution. In a separate flask compound 1 (0.8 g, 3.587 mmol) was dissolved in 5 ml dry DMF. The solution was added dropwise to p-xylenediamine solution over 10 min. The reaction mixture was stirred at 45^o^C for 16 h and then cooled down to room temperature. The TLC was checked in 20% MeOH in DCM (R_f_ = 0.09) to ensure the completion of the reaction. After that, the reaction mixture was filtered to remove solids, and the filtrate was extracted in DCM by washing with water (2 times), brine solution (1 time), and concentrated under reduced pressure. The crude sample was charged on silica gel and purified by column chromatography (eluent: 20% MeOH, 4% NH_3_ in DCM). Compound 2 was thus obtained as a yellow color liquid (370 mg, 1.324 mmol, Yield 37%). LCMS (ESI–MS): calculated 280.20 [M+H] ^+^, found 280.11 [M+H] ^+^. ^1^H–NMR (400 MHz, CD_3_OD): δ 7.31 (d, 4H, ArH), 3.79 (s, 2H, benzyl–CH_2_–), 3.73 ((s, 2H, benzyl–CH_2_–), 3.18 (t, 2H, CO–NH–CH_2_), 2.65 (t, 2H, NH–CH_2_), 1.42 (s, 9H, –C(CH_3_)_3_–). ^1^H–NMR of compound 2 has been shown in figure S27.

**2.2. Synthesis of Compound 4**

Compound 3 has been synthesized according to the literature procedure^[2]^. In a 5 ml pear-shaped flask, compound 2 (100 mg, 0.357 mmol) was taken, and an inert atmosphere was created using N_2_ gas. To this, 2 ml of dry DMF was added, followed by subsequent addition of Et_3_N (0.120 ml, 0.892 mmol). At last, compound 3 (121 mg, 0.393 mmol) was added. The reaction mixture was stirred at RT for 14 h. The TLC was checked in 5% MeOH in DCM (R_f_ = 0.21). The solvent was evaporated under reduced pressure and dissolved in DCM. The organic layer was washed with water (2 times), brine solution (1 time), and dried over Na_2_SO_4_. The crude sample was charged on silica gel and purified by column chromatography (eluent: 5% MeOH in DCM). Compound 4 was thus obtained as a colorless oily liquid (82 mg, 0.173 mmol, Yield 49%). LCMS (ESI–MS): calculated 473.24 [M+H] ^+^, found 473.67 [M+H] ^+^. ^1^H–NMR (400 MHz, CDCl_3_): δ 7.92 (d, 1H, ArH), 7.63 (m, 2H, ArH), 7.43 (m, 1H, ArH), 7.29 ( d, 2H, ArH), 7.20 (d, 2H, ArH), 6.29 (q, 1H, O–CH–), 5.13 (br, 1H, NH–CO), 5.05 (br, 1H, NH–BOC), 4.30 (s, 2H, benzyl–CH_2_– NH–CO), 3.79 (s, 2H, benzyl–CH_2_– NH), 3.24 (t, 2H, CO–NH– CH_2_–), 2.77 (t, 2H, NH– CH_2_–),1.64 (d, 3H, –C–CH_3_), 1.44 (s, 9H, –C(CH_3_)_3_–). ^1^H–NMR of compound 4 has been shown in figure S28.

**2.3. Synthesis of Compound 5**

In a 5 ml pear-shaped flask, compound 4 (54.6 mg, 0.116 mmol) was dissolved in formic acid (5.9 ml, 156.42 mmol) and incubated at room temperature for 1 h. The solvent was evaporated under reduced pressure and thoroughly dried under a high vacuum. The product was dissolved in a minimum amount of ACN: water (1:1) and lyophilized. Compound 5 was thus obtained as a yellow color liquid (17.1 mg, 0.0459 mmol, Yield 87%). HRMS (ESI–MS): calculated 373.1870 [M + H^+^], found 373.1854 [M + H^+^]. ^1^H–NMR (400 MHz, CDCl_3_): δ 8.26 (br, 1H, formate) 7.86 (d, 1H, ArH), 7.60 (m, 2H, ArH), 7.36 (m, 1H, ArH), 7.22 ( d, 2H, ArH), 7.10 (d, 2H, ArH), 6.19 (q, 1H, O–CH–), 6.10 (br, 1H, NH–CO), 4.13 (s, 2H, benzyl–CH_2_– NH–CO), 3.79 (s, 2H, benzyl–CH_2_– NH), 2.97 (t, 4H, –CH_2_–CH_2_–),1.56 (d, 3H, –C–CH_3_). ^1^H–NMR of compound 5 has been shown in figure S29.

**2.4. Synthesis of Compound 6**

In a 25 ml RB flask, 1–adamantylamine (1.0 g, 6.60 mmol) was dissolved in 5 ml dry DMF followed by the addition of K_2_CO_3_ (1.11 g, 8.031 mmol) to the solution. In a separate flask compound 1 (1.48 g, 6.60 mmol) was dissolved in 3 ml dry DMF and added to 1–adamantylamine solution. The reaction mixture was stirred at 45^o^C for 16 h and then cooled down to room temperature. The TLC was checked in 20% MeOH in DCM (R_f_ = 0.2). After that, the reaction mixture was filtered to remove the solids, and the filtrate was extracted in DCM by washing with water (2 times), brine solution (1 time), and concentrated under reduced pressure. The crude sample was charged on silica gel and purified by column chromatography (eluent: 20% MeOH, 1% NH_3_ in DCM). Compound 6 was thus obtained as a yellow color liquid (675 mg, 2.310 mmol, Yield 35%). ^1^H–NMR (400 MHz, CDCl_3_): δ 3.50 (Br, 2H, –CH_2_– NHCO–), 3.27 (Br, 2H, –NH–CH_2_–), 2.83 (s, 1H, –NH–CO–), 2.10 (t, 3H, –CH–), 1.64­–1.72 (m, 12H, –CH_2_–), 1.44 (s, 9H, –(CH_3_)_3_). ^1^H–NMR of compound 6 has been shown in figure S30.

**2.5. Synthesis of Compound 7**

In a 5 ml pear-shaped flask, 4N HCl in dioxane (2 ml, 156.42 mmol) was added to compound 6 (400 mg, 1.36 mmol) at 0^o^C. The reaction mixture was incubated at room temperature for 1 h. The solvent was evaporated under reduced pressure and thoroughly dried under a high vacuum. Afterward, the product was dissolved in a minimum amount of ACN: water (1:1) and lyophilized. Compound 7 was thus obtained as yellow color solid (229 mg, 1.18 mmol, Yield 87%). ^1^H–NMR (400 MHz, D_2_O): δ 3.40 (Br, 4H, –CH_2_– NH–), 2.98 (s, 1H, – NH–), 2.27 (Br, 3H, –CH–), 1.97 (Br, 6H, –CH_2_–), 1.80 (m, 6H, –CH_2_–). ^1^H–NMR of compound 7 has been shown in figure S31.

**3. Synthesis protocol for XYL conjugated quenchers (XYL-Q)**

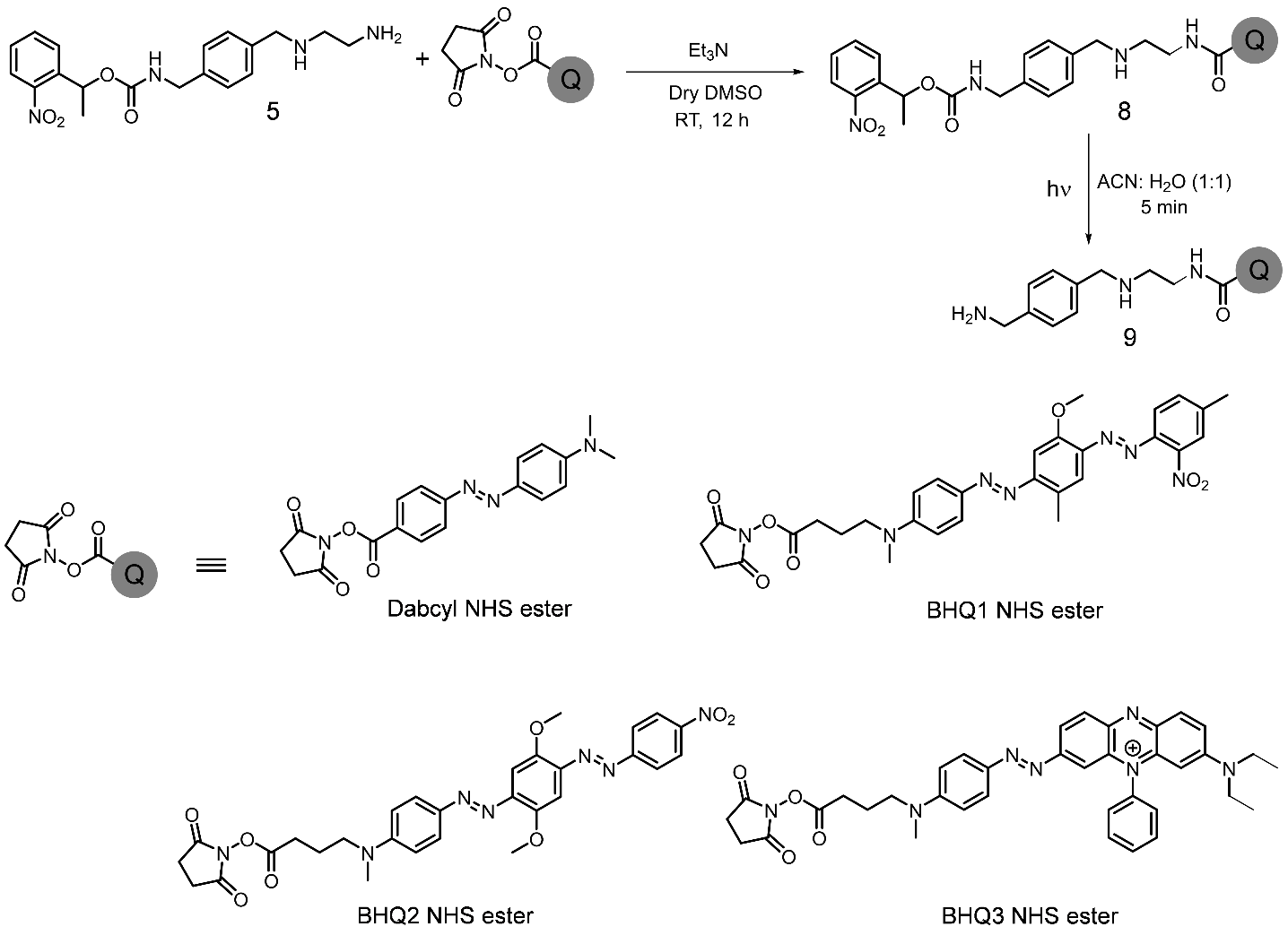

**Scheme S2:** Synthetic scheme for the preparation of XYL–Q.

**3.1. Synthesis of XYL conjugated Dabcyl**

Compound 5 (305 μg, 30.5 μl from 10 mg. ml^–1^ stock in dry DMSO, 0.819 μmol) was taken in a microcentrifuge tube and triethylamine (138 μg, 1.9 μl from 100 μl.ml^–1^ stock in dry DMSO, 1.365 μmol) was added to it. Dabcyl NHS ester (100 μg, 10 μl from 10 mg. ml^–1^ stock in dry DMSO, 0.273 μmol) was added to the above mixture, and the reaction was stirred at room temperature for 12 h. Next, the reaction mixture was made into a 1:1 mixture in water and directly injected into HPLC for purification using water/acetonitrile as eluent. The purified product was characterized by LCMS and dried by lyophilization. The yield calculated is ~75 % (from HPLC). LCMS (ESI–MS): calculated 624.29 [M+H]^+^, 312.65 [M+2H]^2+^; found 624.38 [M+H]^+^, 312.85 [M+2H]^2+^.

The photocleavable group (PCG) protected XYL conjugated product was further dissolved in 1:1 mixture of water/acetonitrile and irradiated with 365 nm UV lamp (50 mW) for 5 min to photocleave the PCG group followed by purified by HPLC using water/acetonitrile as eluent. The purified product was characterized by ^1^H–NMR and HRMS. The yield calculated is ~90% (from HPLC). HRMS (ESI–MS): calculated 431.2554 [M+H]^+^; found 431.2565 [M+H]^+^. ^1^H–NMR (DMSO–d^6^, 600 MHz) δ 8.88 (s, 1H), 8.80 (t, J = 5.7 Hz, 1H), 8.15 (s, 2H), 7.99 (d, J = 8.6 Hz, 2H), 7.85 (d, J = 8.6 Hz, 2H), 7.81 (d, J = 8.9 Hz, 2H), 7.54 (d, J = 7.9 Hz, 2H), 7.50 (d, J = 7.9 Hz, 2H), 6.85 (d, J = 9.3 Hz, 2H), 4.24 (t, J = 5.7 Hz, 2H), 4.06 (q, J = 5.8 Hz, 2H), 3.61 (q, J = 5.9 Hz, 2H), 3.13 (t, J = 6.0 Hz, 2H), 3.08 (s, 6H). The HPLC chromatogram and ^1^H–NMR of XYL–Dabcyl have been shown in figure S14 and S32 respectively.

**3.2. Synthesis of XYL conjugated BHQ1**

Compound 5 (310 μg, 31 μl from 10 mg. ml^–1^ stock in dry DMSO, 0.8331 μmol) was taken in a microcentrifuge tube and triethylamine (84 μg, 1.2 μl from 100 μl.ml^–1^ stock in dry DMSO, 0.8331 μmol) was added to it. BHQ1 NHS ester (100 μg, 10 μl from 10 mg. ml^–1^ stock in dry DMSO, 0.1662 μmol) was added to the above mixture, and the reaction was stirred at room temperature for 12 h. Next, the reaction mixture was made into a 1:1 mixture in water and directly injected into HPLC for purification using water/acetonitrile as eluent. The purified product was characterized by LCMS and dried by lyophilization. The yield calculated is ~60% (from HPLC). LCMS (ESI–MS): calculated 859.39 [M+H]^+^, 430.19 [M+2H]^2+^; found 859.58 [M+H]^+^, 430.39 [M+2H]^2+^.

The PCG protected XYL conjugated product was then dissolved in 1:1 mixture of water/acetonitrile and irradiated with 365 nm UV lamp (50 mW) for 5 min to photocleave the PCG group followed by purified by HPLC using water/acetonitrile as eluent. The purified product was characterized by ^1^H–NMR and HRMS. The yield calculated is ~95% (from HPLC). HRMS (ESI–MS): calculated 666.3511 [M+H]^+^, 333.6792 [M+2H]^2+^; found 666.3310 [M+H]^+^, 333.6763 [M+2H]^2+^. ^1^H–NMR (DMSO–d^6^, 600 MHz) δ 8.81 (s, 2H), 8.13 (t, J = 5.8 Hz, 3H), 7.95 (s, 1H), 7.81 (d, J = 9.9 Hz, 2H), 7.77 (d, J = 8.2 Hz, 1H), 7.69 (d, J = 7.2 Hz, 1H), 7.51 (dd, J = 15.6, 8.4 Hz, 4H), 7.29 (s, 1H), 6.88 (d, J = 9.3 Hz, 2H), 4.20 (t, J = 5.8 Hz, 2H), 4.05 (q, J = 5.8 Hz, 2H), 3.92 (s, 3H), 3.43-3.66 (m, 4H merged with H_2_O peak), 3.06 (s, 3H), 2.97 (t, J = 6.0 Hz, 2H), 2.63 (s, 3H), 2.55-2.38 (m, 3H merged with DMSO peak), 2.20 (t, J = 7.4 Hz, 2H), 1.78-1.83 (m, 2H). The HPLC chromatogram and ^1^H–NMR of XYL–BHQ1 have been shown in figure S15 and S33 respectively.

**3.3. Synthesis of XYL conjugated BHQ2**

Compound 5 (308 μg, 30.8 μl from 10 mg. ml^–1^ stock in dry DMSO, 0.8283 μmol) was taken in a microcentrifuge tube and triethylamine (83.65 μg, 1.2 μl from 100 μl.ml^–1^ stock in dry DMSO, 0.8283 μmol) was added to it. BHQ2 NHS ester (100 μg, 10 μl from 10 mg. ml^–1^ stock in dry DMSO, 0.1657 μmol) was added to the above mixture, and the reaction was stirred at room temperature for 12 h. Next, the reaction mixture was made into a 1:1 mixture in water and directly injected into HPLC for purification using water/acetonitrile as eluent. The purified product was characterized by LCMS and dried by lyophilization. The yield calculated is ~90% (from HPLC). LCMS (ESI–MS): calculated 861.36 [M+H]^+^, 431.19 [M+2H]^2+^; found 861.54 [M+H]^+^, 431.31 [M+2H]^2+^.

The PCG protected XYL conjugated product was then dissolved in 1:1 mixture of water/acetonitrile and irradiated with 365 nm UV lamp (50 mW) for 5 min to photocleave the PCG group followed by purified by HPLC using water/acetonitrile as eluent. The purified product was characterized by ^1^H–NMR and HRMS. The yield calculated is ~100% (from HPLC). HRMS (ESI–MS): calculated 668.3303 [M+H]^+^, 334.6688 [M+2H]^2+^; found 668.3310 [M+H]^+^, 334.6690 [M+2H]^2+^. ^1^H–NMR (DMSO–d^6^, 600 MHz) δ 8.78 (s, 2H), 8.44 (d, J = 8.9 Hz, 2H), 8.11 (t, J = 5.7 Hz, 3H), 8.06 (d, J = 8.9 Hz, 2H), 7.82 (d, J = 8.9 Hz, 2H), 7.49 (q, J = 8.0 Hz, 4H), 7.37 (s, 1H), 6.88 (d, J = 9.3 Hz, 2H), 4.18 (t, J = 5.8 Hz, 2H), 4.04 (q, J = 5.8 Hz, 2H), 3.99 (s, 3H), 3.94 (s, 3H), 3.51-3.39 (m, 2H merged with H_2_O peak), 3.06 (s, 3H), 2.96 (t, J = 6.2 Hz, 2H), 2.59-2.39 (m, 2H merged with DMSO peak), 2.19 (t, J = 7.4 Hz, 2H), 1.80 (t, J = 7.2 Hz, 2H). The HPLC chromatogram and ^1^H–NMR of XYL–BHQ2 have been shown in figure S16 and S34 respectively.

**3.4. Synthesis of Xyl conjugated BHQ3**

Compound 5 (288.84 μg, 28.9 μl from 10 mg. ml^–1^ stock in dry DMSO, 0.7754 μmol) was taken in a microcentrifuge tube and triethylamine (78.3 μg, 1.1 μl from 100 μl.ml^–1^ stock in dry DMSO, 0.7754 μmol) was added to it. BHQ3 NHS ester (100 μg, 10 μl from 10 mg. ml^–1^ stock in dry DMSO, 0.1550 μmol) was added to the above mixture, and the reaction was stirred at room temperature for 12 h. Next, the reaction mixture was made into a 1:1 mixture in water and directly injected into HPLC for purification using water/acetonitrile as eluent. The purified product was characterized by LCMS and dried by lyophilization. The yield calculated is ~40% (from HPLC). LCMS (ESI–MS): calculated 901.45 [M]^+^, 451.23 [M+H]^2+^; found 901.64 [M]^+^, 451.38 [M+H]^2+^.

The PCG protected Xyl conjugated product was then dissolved in 1:1 mixture of water/acetonitrile and irradiated with 365 nm UV lamp (50 mW) for 5 min to photocleave the PCG group followed by purified by HPLC using water/acetonitrile as eluent. The purified product was characterized by ^1^H–NMR and HRMS. The yield calculated is ~99% (from HPLC). HRMS (ESI–MS): calculated 708.4133 [M]^+^, 354.7103 [M+H]^2+^; found 708.4136 [M]^+^, 354.7105 [M+H]^2+^. ^1^H–NMR (DMSO–d^6^, 600 MHz) δ 8.84-8.76 (2H), 8.43 (d, J = 8.9 Hz, 2H), 8.19-8.21 (m, 2H), 8.15 (s, 3H), 8.00 (d, J = 12.3 Hz, 1H), 7.90-7.95 (m, 2H), 7.77-7.80 (m, 3H), 7.50 (d, J = 5.5 Hz, 4H), 7.23 (s, 1H), 6.88 (d, J = 9.3 Hz, 2H), 5.70 (s, 1H), 4.18-4.20 (m, 2H), 4.04-4.07 (m, 2H), 3.56-3.82 (m, 6H merged with H_2_O peak), 3.09 (s, 3H), 2.95-2.99 (m, 2H), 2.41-2.65 (m, 2H merged with DMSO peak), 2.19 (t, J = 6.7 Hz, 2H), 1.77-1.81 (m, 2H), 1.23 (s, 6H). The HPLC chromatogram and ^1^H–NMR of XYL–BHQ3 have been shown in figure S17 and S35 respectively.

**4. Synthesis protocol for EtA conjugated quenchers (EtA-Q)**

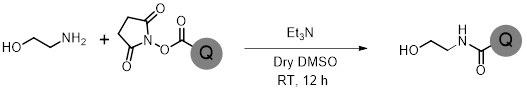

**Scheme S3:** Synthetic scheme for the preparation of EtA–Q.

**4.1. Synthesis of EtA–Dabcyl conjugate**

Ethanolamine (166.67 μg, 1.67 μl from 1:10 diluted stock in dry DMSO, 2.7295 μmol) was taken in a microcentrifuge tube and triethylamine (1.9 μl from 1:10 diluted stock in dry DMSO, 1.3647 μmol) was added to it and mixed by vortex mixture. Dabcyl NHS ester (100 μg, 0.2729 μmol) was added to the above mixture and the reaction was stirred at room temperature for 12 h. Next, the reaction mixture was diluted to a 1:1 mixture in water and directly injected into HPLC for purification (figure S18) using water/acetonitrile as eluent. The purified compound was characterized by HRMS. The yield calculated is ~95% (from HPLC). HRMS (ESI–MS): calculated 313.1659 [M+H]^+^; found 313.1649 [M+H]^+^.

**4.2. Synthesis of EtA–BHQ1 conjugate**

Ethanolamine (101.20 μg, 1.02 μl from 1:10 diluted stock in dry DMSO, 1.6620 μmol) was taken in a microcentrifuge tube and triethylamine (1.16 μl from 1:10 diluted stock in dry DMSO, 0.8283 μmol) was added to it and mixed by vortex mixture. BHQ1 NHS ester (100 μg, 0.1662 μmol) was added to the above mixture and the reaction was stirred at room temperature for 12 h. Next, the reaction mixture was diluted to a 1:1 mixture in water and directly injected into HPLC for purification (figure S19) using water/acetonitrile as eluent. The purified compound was characterized by HRMS. The yield calculated is ~95% (from HPLC). HRMS (ESI–MS): calculated 548.2616 [M+H]^+^; found 548.2604 [M+H]^+^.

**4.3. Synthesis of EtA–BHQ2 conjugate**

Ethanolamine (101.20 μg, 1.02 μl from 1:10 diluted stock in dry DMSO, 1.657 μmol) was taken in a microcentrifuge tube and triethylamine (1.16 μl from 1:10 diluted stock in dry DMSO, 0.8283 μmol) was added to it and mixed by vortex mixture. BHQ2 NHS ester (100 μg, 0.1657 μmol) was added to the above mixture and the reaction was stirred at room temperature for 12 h. Next, the reaction mixture was diluted to a 1:1 mixture in water and directly injected into HPLC for purification (figure S20) using water/acetonitrile as eluent. The purified compound was characterized by HRMS. The yield calculated is ~95% (from HPLC). HRMS (ESI–MS): calculated 550.2409 [M+H]^+^; found 550.2388 [M+H]^+^.

**4.4. Synthesis of EtA–BHQ3 conjugate**

Ethanolamine (77.8 μg, 7.78 μl from 1:100 diluted stock in dry DMSO, 1.2660 μmol) was taken in a microcentrifuge tube and triethylamine (8.8 μl from 1:100 diluted stock in dry DMSO, 0.6333μmol) was added to it and mixed by vortex mixture. BHQ3 NHS ester (100 μg, 0.1551 μmol) was added to the above mixture and the reaction was stirred at room temperature for 12 h. Next, the reaction mixture was diluted to a 1:1 mixture in water and directly injected into HPLC for purification (figure S21) using water/acetonitrile as eluent. The purified compound was characterized by HRMS. The yield calculated is ~60% (from HPLC). HRMS (ESI–MS): calculated 590.3238 [M]^+^; found 590.3214 [M]^+^.

**5. Synthesis protocol for tetrazine conjugated BODIPY fluorophore**

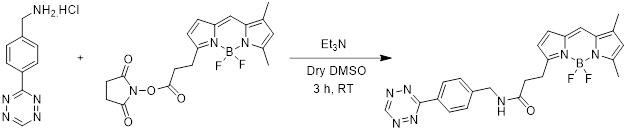

**Scheme S4:** Synthetic scheme for preparing tetrazine–BODIPY fluor.

Tetrazine-BODIPY was synthesized using the previously reported protocol^[3]^. Benzyl amine derivative of tetrazine (241.2 μg, 24.12 μl from 10 mg. ml^–1^ stock in DMSO, 1.2848 μmol) was taken in a microcentrifuge tube. Triethylamine (259.57 μg, 3.6 μl from 1:10 (v/v) stock in DMSO, 2.5700 μmol) was added to it. BODIPY NHS ester (100 μg, 5 μl from 10 mg. ml^–1^ stock in DMSO, 0.2570 μmol) was added to the reaction mixture and stirred at room temperature for 3 h. After that, the reaction mixture was diluted using water and directly injected into HPLC for purification. The purified product was dried under vacuum and characterized by LCMS. The yield calculated is ~50% (from HPLC). LCMS (ESI–MS): calculated 462.20 [M+H]^+^; found 462.47 [M+H]^+^.

**6. Synthesis of NHS ester derivatives of ADA**

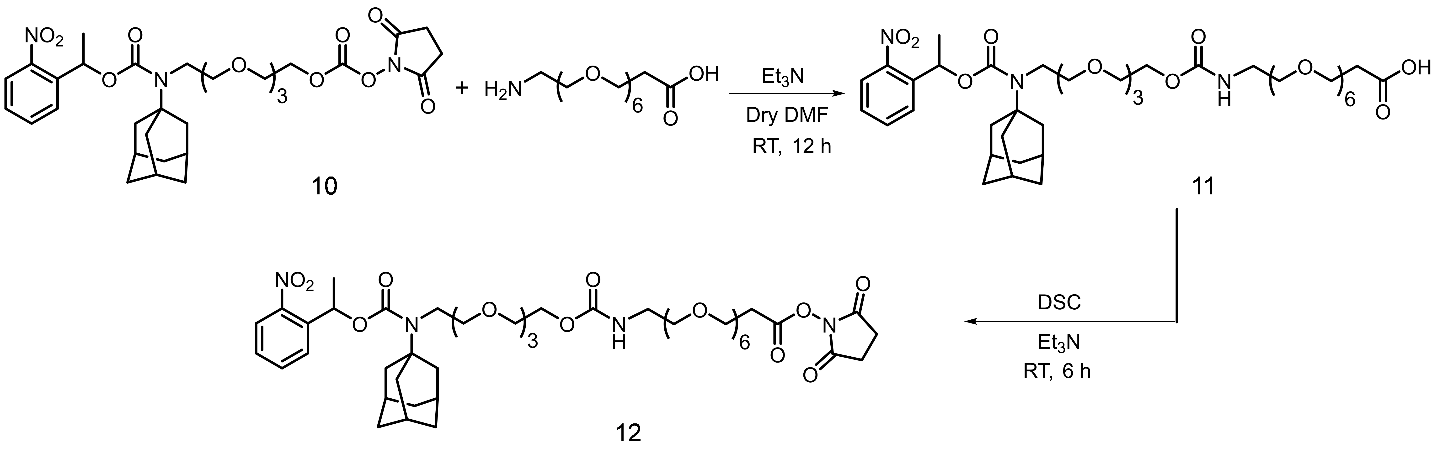

**Scheme S5:** Synthetic scheme for the preparation of nitrobenzyl-protected ADA–PEG–NHS ester

**Note –** Compounds 11 and 12 were synthesized using our previously reported protocol.^[4]^

**7. Conjugation of ADA with the antibody**

1. Secondary antibodies (Donkey anti–Rat and Donkey anti-human) were buffer exchanged by Zeba^TM^ spin column pre-equilibrated with PBS containing 10% 1M NaHCO_3_.
2. Buffer exchanged secondary antibody (50 µg, 3.33x10^-4^ µmol) was first taken in a 1.5 ml microcentrifuge tube.
3. Compound 12 (3.32 µg, 3.33x10^-3^ µmol, 4.5 µl solution from of 0.75 mg ml^−1^ in DMF) were added to the antibody solutions in two portions (2.25 µl each time) to the buffer exchanged secondary antibodies.
4. The reaction was kept for gentle shaking at 4^o^C for 12 h.
5. After that nitrobenzyl-protected ADA conjugated antibody was purified by Zeba spin column (pre-equilibrated three times with PBS).
6. The purified antibody was then subjected to irradiation using 365 nm UV light (50 mW.cm^–2^) for 5 min to generate ADA conjugated antibody.
7. The ADA conjugated antibody was then characterized by MALDI mass spectrometry (figure S26) and proceeded with the fluorogenic microscopy experiment.

**8. Conjugation of ADA with phalloidin**

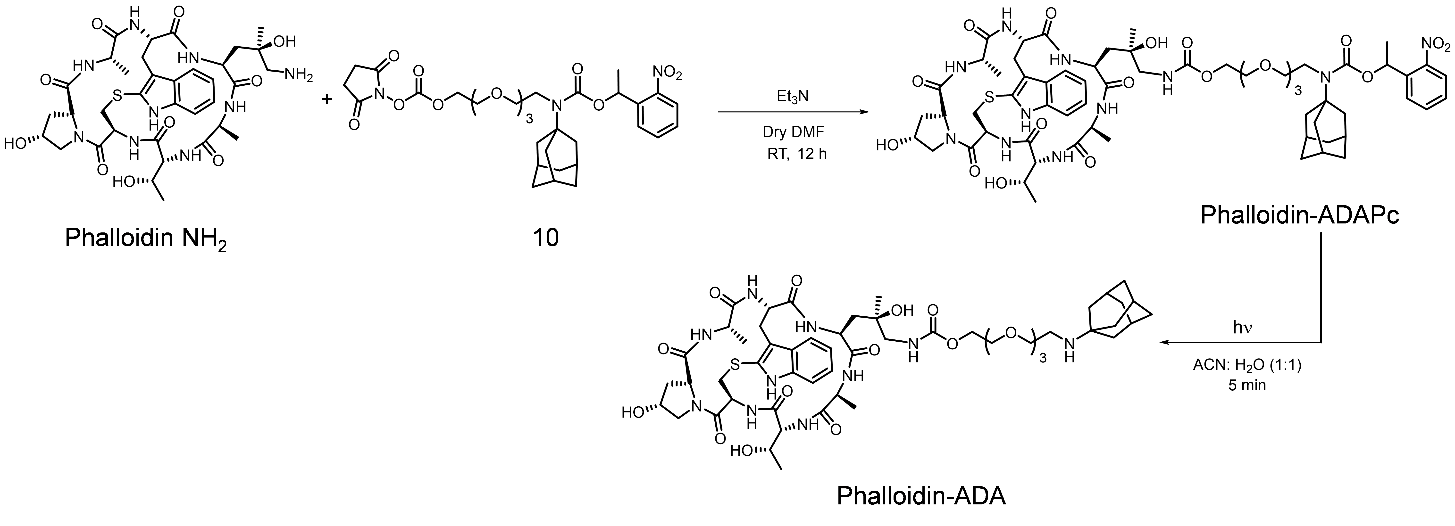

**Scheme S6:** Synthetic scheme for the preparation of ADA conjugated phalloidin**.**

1. Phalloidin–ADAPc was synthesized using previously reported protocols by our group^4^.
2. The ADAPc conjugated phalloidin was then dissolved in a 1:1 mixture of water/acetonitrile and irradiated with a 365 nm UV lamp (50 mW) for 5 min to photocleave the Pc group followed by purified by HPLC using water/acetonitrile as eluent.
3. The ADA conjugated phalloidin was then characterized by HRMS and proceeded for the fluorogenic microscopy experiment. Yield: quantitative (from HPLC). HRMS (ESI-MS): Calculated m/z 1141.5598 [M+H]^+^; found 1141.5598 [M+H]^+^.

**9. Conjugation of ADA with taxol**

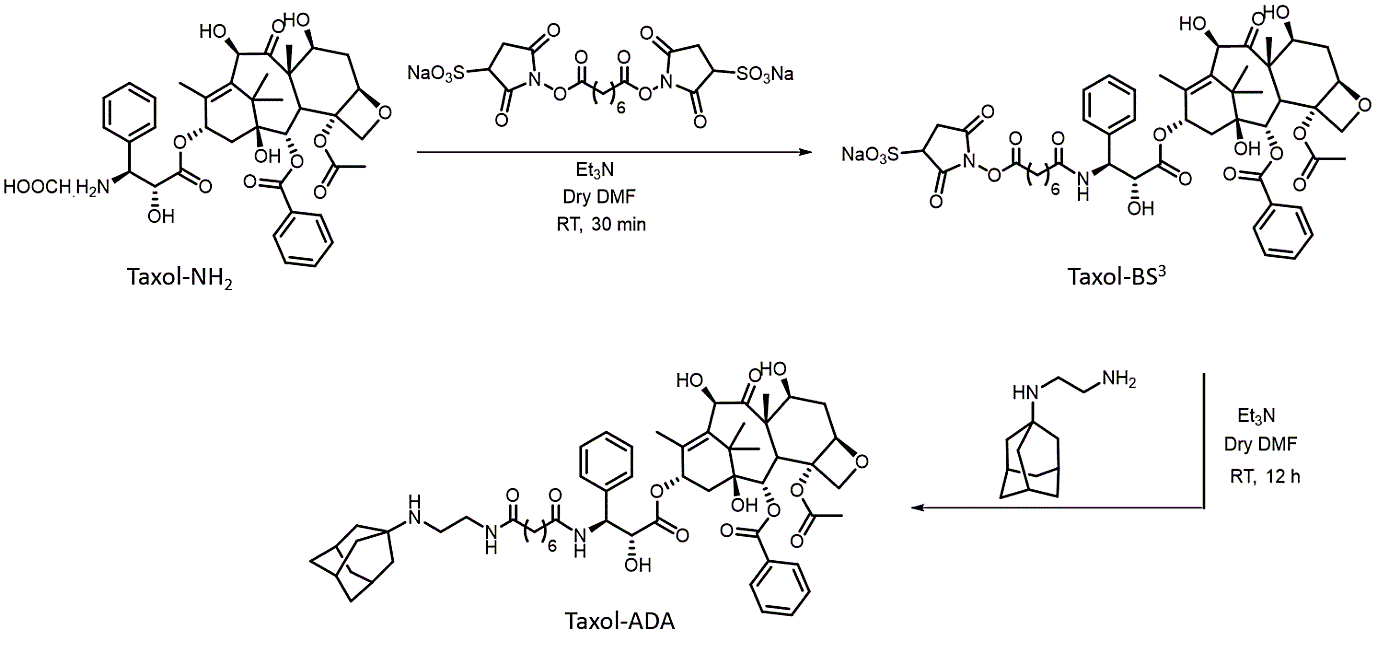

**Scheme S7:** Synthetic scheme for the preparation of ADA conjugated taxol**.**

1. Amine derivative of docetaxel (80 µg, 16 µl from 5 mg ml^−1^ stock solution in dry DMF, 0.1061 µmol) was taken in a 0.5 ml microcentrifuge tube.
2. 4.2 µl DMF solution containing 0.042 µl Et_3_N was added to it.
3. Bis(sulfosuccinimidyl) suberate (BS^3^) (1.5 mg, 2.84 µmol, dissolved in 30 µl DMF) was added to it and the reaction was stirred at room temperature for 30 min.
4. After that, the reaction mixture was diluted using 50 µl of Mili Q water and injected into HPLC for purification using water/acetonitrile containing 0.1% TFA as eluent.
5. The taxol-BS3 conjugate was characterized by using LCMS. LCMS (ESI-MS): Calculated m/z 1041.35 [M+H]^+^; found 1041.56 [M+H]^+^.
6. The purified compound (Yield: quantitative from HPLC chromatogram) was then dissolved in dry DMF for conjugation with ADA.
7. In a separate microcentrifuge tube, compound 7 (0.187 mg, 1.016 µmol, dissolved in 20 µl DMF) was taken and triethylamine (2.82 µl DMF solution containing 0.141 µl Et_3_N) was added to it.
8. Next, the solutions were mixed together and allowed to stir at room temperature for 12 h.
9. After the completion of the reactions, the reaction mixture was diluted using 50 µl of Mili Q water and injected into HPLC for purification using water/acetonitrile containing 0.1% TFA as eluent.
10. The ADA conjugated taxol was then characterized by LCMS and ^1^H NMR. LCMS (ESI-MS): Calculated m/z 1040.55 [M+H]^+^; found 1040.73 [M+H]^+^.

^1^H NMR (DMSO–d^6^, 600 MHz) δ 8.36 (d, J = 8.0 Hz, 1H), 8.00 (t, J = 7.8 Hz, 2H), 7.84 (d, J = 8.4 Hz, 1H), 7.71 – 7.59 (m, 2H), 7.48 – 7.18 (m, 5H), 5.95 – 5.88 (m, 2H), 5.43 (d, J = 7.2 Hz, 1H), 5.28 (dd, J = 6.0, 3.0Hz, 1H), 5.17 – 4.90 (m, 4H), 4.59 (s, 1H), 4.42 (t, J = 6.6 Hz, 1H), 4.12 – 3.98 (m, 3H), 3.76 (q, J = 6.6 Hz, 1H), 2.99 (s, 1H), 2.93 – 2.87 (m, 2H), 2.29 (s, 1H), 2.24 (s, 3H), 2.20 – 2.15 (m, 2H), 2.12 (br, 3H), 2.09 – 1.80 (m, 6H), 1.79 – 1.74 (m, 8H), 1.69 – 1.58 (m, 6H), 1.53 (s, 3H), 1.46 (q, J = 6.6 Hz, 4H), 1.21 – 1.09 (m, 4H), 1.06 – 0.99 (m, 6H). The HPLC and ^1^H–NMR of ADA-taxol conjugate have been shown in figure S23 and S36 respectively.

**10. Conjugation of TCO with phalloidin**

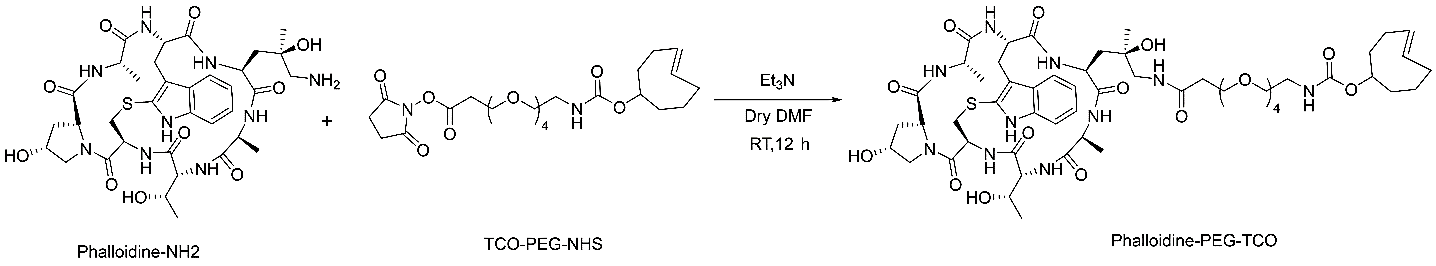

**Scheme S8:** Synthetic scheme for the preparation of TCO conjugated phalloidin.

1. Phalloidin amine (10 μg, 1 μl from 10 mg. ml^–1^ stock in DMSO, 0.0127 μmol) was taken in a microcentrifuge tube and diluted to 10 μl using dry DMSO.
2. Triethylamine (6.09 μg, 8.5 μl from 1:1000 (v/v) stock in DMSO, 0.6029 μmol) was added.
3. TCO NHS ester (31.2 μg, 3.12μl from 10 mg. ml^–1^ stock in DMSO, 0.6029 μmol) was added to the reaction mixture and stirred at room temperature for 12 h.
4. After the reaction, the reaction mixture was diluted to a 1:1 water/DMSO mixture and directly injected into HPLC for purification (figure S24).
5. The purified product was dried under vacuum and characterized by LCMS. The yield calculated is ~80% (from HPLC). LCMS (ESI–MS): calculated 1187.56 [M+H]^+^, 1209.54 [M+Na]^+^; found 1187.81 [M+H]^+^, 1209.78 [M+Na]^+^.

**11. Conjugation of TCO with jasplakinolide**

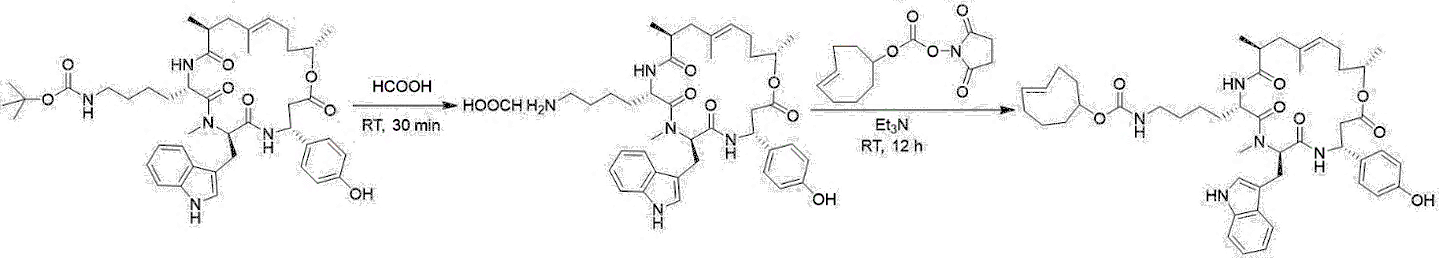

**Scheme S9:** Synthetic scheme for the preparation of TCO–jasplakinolide conjugate.

**11.1. Jasplakinolide–lysine amine preparation**

Boc protected lysine derivative of jasplakinolide (100 μg, 0.1291 μmol) was dissolved in 100 μl formic acid and incubated for 30 min at room temperature. The formic acid solution was diluted using water and dried under a vacuum for 12 h to remove the acid traces completely. The crude jasplakinolide lysine derivative (considering quantitative yield) was dissolved in dry DMSO and characterized by LCMS. LCMS (ESI–MS): calculated 674.39 [M+H]^+^ ; found 674.57 [M+H]^+^.

**11.2. Jasplakinolide–TCO conjugate**

1. Jasplakinolide–lysine amine (10 μg, 1 μl from 10 mg.ml^–1^ stock in DMSO, 0.015 μmol) was taken in a microcentrifuge tube and diluted to 10 μl using dry DMSO.
2. Triethylamine (4.54 μg, 6.3 μl from 0.1% (v/v) stock in DMSO, 0.045 μmol) was added.
3. TCO NHS ester (61.5 μg, 6.15 μl from 10 mg. ml^–1^ stock in DMSO, 0.23 μmol) was added into the reaction mixture and stirred at room temperature for 12 h.
4. After the reaction, the reaction mixture was diluted to a 1:1 water/DMSO mixture and directly injected into HPLC for purification (figure S25).
5. The purified product was dried under vacuum and characterized by HRMS. The yield calculated is ~70% (from HPLC). HRMS (ESI–MS): calculated 826.4749 [M+H]^+^, 848.4569 [M+Na]^+^; found 826.4676 [M+H]^+^, 848.4496 [M+Na]^+^.

**12. Synthetic Scheme for XYL–AuNP**

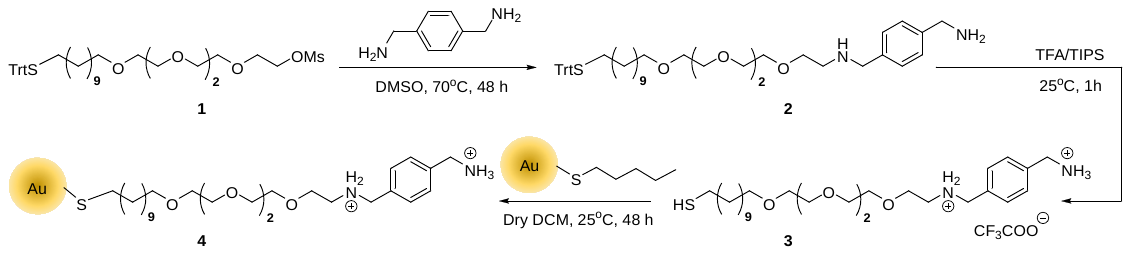

**Scheme S10**: Synthetic scheme for the preparation of XYL–AuNP.

**12.1. Synthesis of Compound 2 (****Trt-C11-TEG-XYL)**

In a 100 ml round-bottom flask xylene diamine (1.11 g, 8.16 mmol) was taken and dissolved with 5 ml of dry DMSO. In a separate 100 ml RB, Trt-C11-TEG-XYL^[5]^ (compound 1, 280 mg, 0.39 mmol) was dissolved in 2 ml of dry DMSO. Next, the DMSO solution of Compound 1 was added to the xylene diamine solution (in DMSO) slowly in a dropwise manner. The reaction mixture was stirred at 70^0^C for 48 h. After that, the reaction mixture was dissolved in DCM and washed with water (30 ml × 2), and brine (30 ml × 2), dried over Na_2_SO_4_ and concentrated under reduced pressure. Next, the reaction mixture was purified using reverse-phase flash column chromatography (C-18 20 g column used) with water/acetonitrile as eluent. The purified product was characterized by LCMS and dried by lyophilization. Compound 2 was thus obtained as a yellowish-brown oil (265 mg, Yield 89.5 %). ^1^H–NMR (600 MHz, DMSO-d^6^): δ 8.32 (s, 2H), 7.51 (q, 4H) 7.35 – 7.23 (m, 15H), 4.19 (s, 2H), 4.06 (s, 2H), 3.55 (s, 2H), 3.33 – 3.50 (m, 16H), 2.06 (t, 2H), 1.45 (m, 2H), 1.06-1.28 (m, 18H). LCMS (ESI-MS): calculated 741.56 [M+H]^+^; obtained 741.72 [M+H]^+^. The ^1^H–NMR of Compound 2 has been shown in figure S37.

**12.2. Synthesis of Compound 3 (HS-C11-TEG-XYL)**

Compound 2 (100 mg, 0.13 mmol) was taken with trifluoroacetic acid (TFA, 2.6 mmol) and triisopropylsilane (TIPS, 0.39 mmol) in dry DCM (3ml). During the addition of TFA, the solution slowly turned yellow, which after the addition of TIPS resulted in a colorless solution. The reaction mixture was stirred at RT for 5 h under an N_2_ atmosphere. After the completion of the reaction, the volatile components (solvent, TFA, TIPS) were removed under reduced pressure and the obtained residue was washed with hexane (4 times). The compound 3 obtained was dried (64.0 mg, yield 65.7 %) and characterized by ­^1^H NMR. ^1^H NMR (400 MHz, DMSO-d^6^): δ 9.03 (s, 2H), 8.27 (s, 2H), 7.55 – 7.49 (m, 4H), 4.20 (s, 2H), 4.07 (s, 2H), 3.69 (t, *J* = 5.2 Hz, 2H), 3.57 – 3.34 (m, 14H), 3.08 (d, *J* = 7.2 Hz, 2H), 1.61 – 1.34 (m, 4H), 1.25 – 1.21 (s, 14H). HRMS (ESI-MS): calculated 499.3564 [M+H]^+^; obtained 499.4186 [M+H]^+^. The ^1^H–NMR of Compound 3 has been shown in figure S38.

**12.3. Surface functionalization of AuNP (Compound 4)**

To fabricate AuNPs, we followed a two-step procedure. Firstly, we synthesized 1-Pentanethiol-coated gold nanoparticles (d ~ 2 nm) using the literature procedure.^[6]^ Xylene functionalized gold nanoparticles (XYL–AuNP) were prepared by place exchange of pentanethiol capped ~2 nm gold nanoparticles (Au-C_5_).

In a 20 ml vial Au-C_5_ (3.2 mg) was dissolved in nitrogen-purged dry DCM (1 ml). In another vial xylene diamine thiol ligand (compound 3, 15.6 mg) was dissolved in nitrogen-purged dry DCM (1 ml) + methanol (0.1 ml) and transferred to the first vial. The reaction was stirred at RT for 48 h under an N_2_ atmosphere. The solvent was evaporated under reduced pressure and the nanoparticles were washed 3 times with hexane. Nanoparticles were recovered by centrifugation and the supernatant was discarded. After that, the nanoparticles were redispersed in water and then dialyzed for 48h using Snake Skin Dialysis Membrane 10K to get the surface functionalized XyL-AuNP. Then, the surface functionalized XyL-AuNP was characterized by MALDI (Figure S39). The concentration of the functionalized AuNPs was measured to be 9.13 µM. The MALDI spectrum showed a peak at m/z = 499.072 (observed) which corresponds to the calculated mass value (m/z = 499.356) of the thiol ligand of xylene diamine moiety.

**13. Other experimental protocols and supporting results**

**13.1. Protocol for fluorescence quenching study of CB7–FLs with XYL–Qs**

The fluorescence quenching titrations of CB7–FLs vs XYL–Qs were performed using a fluorescence microplate reader. In this experiment, CB7–FLs were taken in a black 96 well microplate (Corning, non–binding plate) with a final concentration of 1 μM in 200 µl PBS buffer. Fluorescence spectra were recorded at respective excitation and emission wavelength scans listed in the table below (Table S5a). After this, 0.25 eq of XYL–Qs (final concentration 0.25 μM in 200 µl) were added to the system followed by the recording of fluorescence spectra. The addition of XYL–Qs (0.25 eq each) was subsequently continued followed by spectra recording for the respective CB7–FLs to achieve maximum quenching.

Similar protocols have been followed for quenching studies of CB7–FL against EtA–Q.

**13.2. Protocol for fluorogenic response study from CB7–XYL quenched complex**

The CB7–FLs were taken in a black 96 well microplate (Corning, non–binding plate) with a final concentration of 1 μM in 200 µl PBS buffer. The respective equivalent of XYL–Qs were added to each well of CB7–FLs to achieve maximum quenching of fluorophores (Table S6). After this, ADA–NH_2_ (final concentration 0.25 µM, 0.25 eq) was added to the quenched [CB7–FL∙XYL–Q] systems and fluorescence recovery kinetics were recorded for 10 min with a 30 s interval. After 10 min, another 0.25 eq ADA–NH_2_ was added, and recovery kinetics was again recorded. This protocol was followed for 4 successive additions of ADA (total 1 eq, final concentration 1 µM). The excitation and emission wavelengths that are used for kinetics measurements are listed in Table S5b.

**Table S5a**: Table for excitation and emission wavelength of CB7–FL used in fluorogenic response experiment.

| **CB7–FL** | **XYL–Q/EtA–Q** | **Excitation wavelength**  **(λ_ex_)** | **Emission range**  **(λ_em_)** |
| --- | --- | --- | --- |
| Coumarin | Dabcyl | 350 nm | (425-525) nm |
| Fluorescein | BHQ1 | 465 nm | (500-600) nm |
| BODIPY | BHQ1 | 465 nm | (500-600) nm |
| Cy3 | BHQ2 | 520 nm | (550-650) nm |
| TAMRA | BHQ2 | 520 nm | (550-650) nm |
| Cy5 | BHQ3 | 620 nm | (650-750) nm |
| SiR | BHQ3 | 620 nm | (650-750) nm |

**Table S5b**: Table for excitation and emission wavelength of CB7–FL used in fluorescence recovery experiment.

| **CB7–FL** | **XYL–Q** | **Excitation wavelength**  **(λ_ex_)** | **Emission wavelength**  **(λ_em_)** |
| --- | --- | --- | --- |
| Coumarin | Dabcyl | 350 nm | 450 nm |
| Fluorescein | BHQ1 | 465 nm | 520 nm |
| BODIPY | BHQ1 | 465 nm | 515 nm |
| Cy3 | BHQ2 | 510 nm | 565 nm |
| TAMRA | BHQ2 | 510 nm | 585 nm |

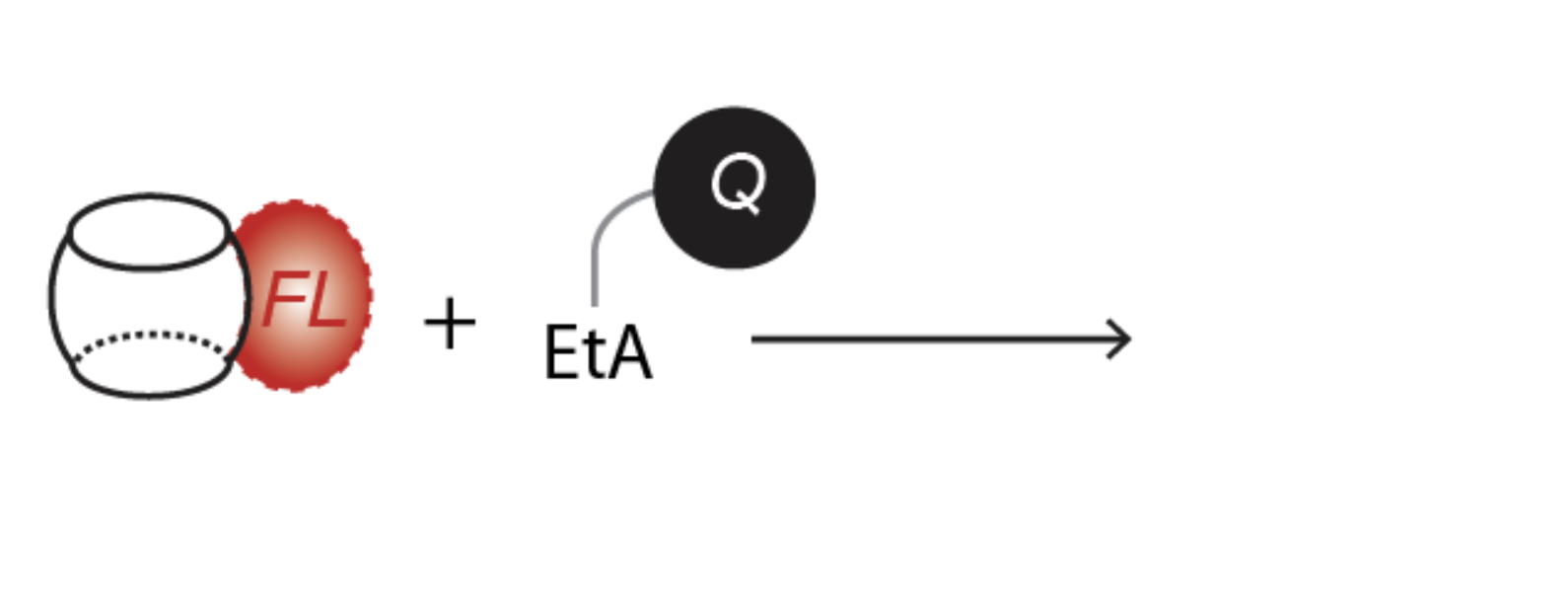

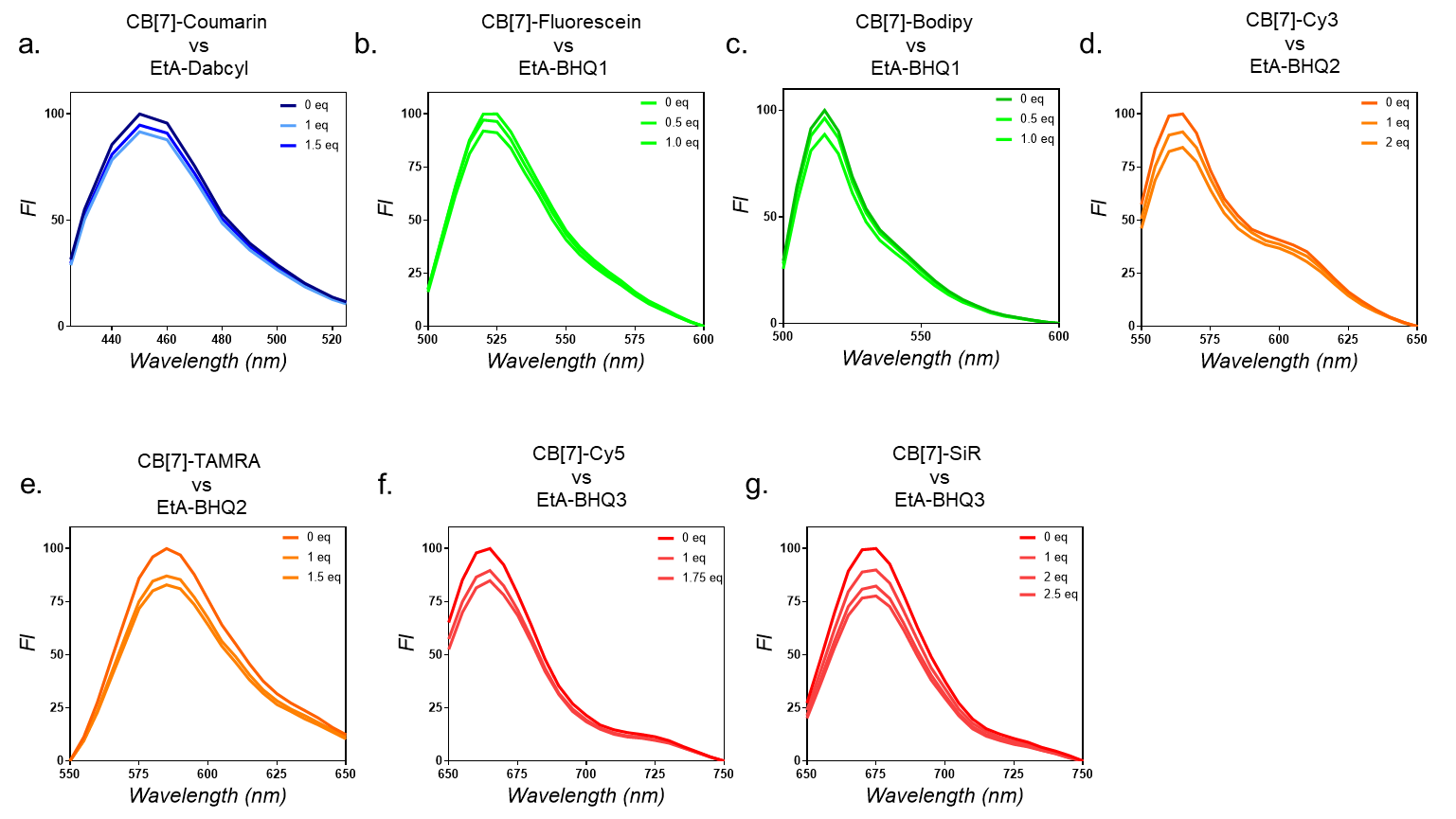

**Figure S1:** Fluorescence titration data for CB7–FL vs Ethanolamine (EtA–Q). Negligible fluorescence quenching from the non–interacting pair demonstrated that the fluorescence quenching requires specific recognition of CB7 host and XYL guest moiety.

**Table S6. The concentration of XYL-Q quencher used with respect to CB7-FL**

| SL No | CB7-FL | XYL-Q |
| --- | --- | --- |
| 1 | CB7-Coumarin (1.0 µM) | XYL-Dabcyl (1.5 µM) |
| 2 | CB7-Fluorescein (1.0 µM) | XYL-BHQ1 (1.0 µM) |
| 3 | CB7-Bodipy (1.0 µM) | XYL-BHQ1 (1.0 µM) |
| 4 | CB7-Cy3 (1.0 µM) | XYL-BHQ2 (2.0-2.5 µM) |
| 5 | CB7-TAMRA (1.0 µM) | XYL-BHQ2 (1.5 µM) |
| 6 | CB7-Cy5 (1.0 µM) | XYL-BHQ3 (1.5-2.5 µM) |
| 7 | CB7-SiR (1.0 µM) | XYL-BHQ3 (2.5 µM) |

**13.3. Protocol for MALDI-MS analysis**

I. The CB7-FL (FL = Coumarin, TAMRA, SiR, and Cy5) was taken in 1 µM concentration for the study. Xyl-Q was taken in concentration with respect to the type of quencher [Xyl-Q, Q = Dabcyl (1 µM), BHQ2 (2 µM), BHQ3 (3 µM)]. The respective quencher and fluorophore were combined together in appropriate concentrations in Mili Q water [CB-Cy5(1 µM) + Xyl-BHQ3(3 µM), CB-SiR(1 µM) + Xyl-BHQ3(3 µM), CB-TAMRA(1 µM) + Xyl-BHQ2(2 µM), CB-Coumarin(1 µM) + Xyl-Dabcyl(1 µM)]. The resulting solution was incubated for 30 min at RT. Next, the same solution (2 µL) was combined with 2 µL of matrix solution of α-cyano-4-hydroxycinnamic acid (solvent – 70:30 Acetonitrile/Water) for MALDI-MS analysis.

II. The displacement of the Xyl-Q quencher from CB-FL·Xyl-Q complex (made using above protocol I) was analyzed after adding 1-Adamantylamine hydrochloride (ADA) to the solution. The concentration of ADA is used the same as the respective quencher concentration. The resulting solution was incubated for 30 min before performing MALDI-MS analysis (matrix same as protocol I).

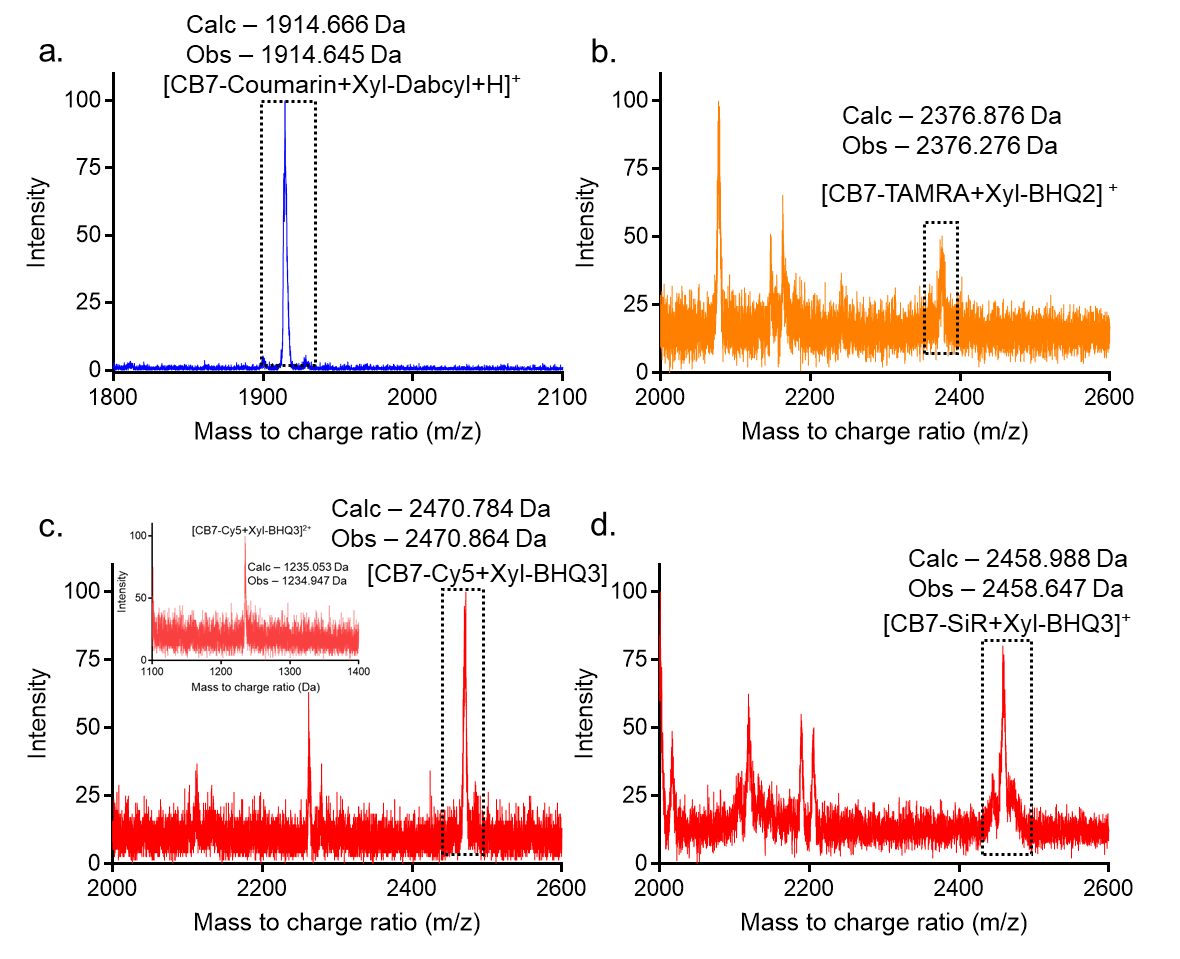

**Figure S2:** MALDI-MS spectroscopic data showing the detection of CB7-FL∙Xyl-Q monovalent complex. a) CB7-Coumarin∙Xyl-Dabcyl b) CB7-TAMRA∙Xyl-BHQ2 c) CB7-Cy5∙Xyl-BHQ3 d) CB7-SiR∙Xyl-BHQ3.

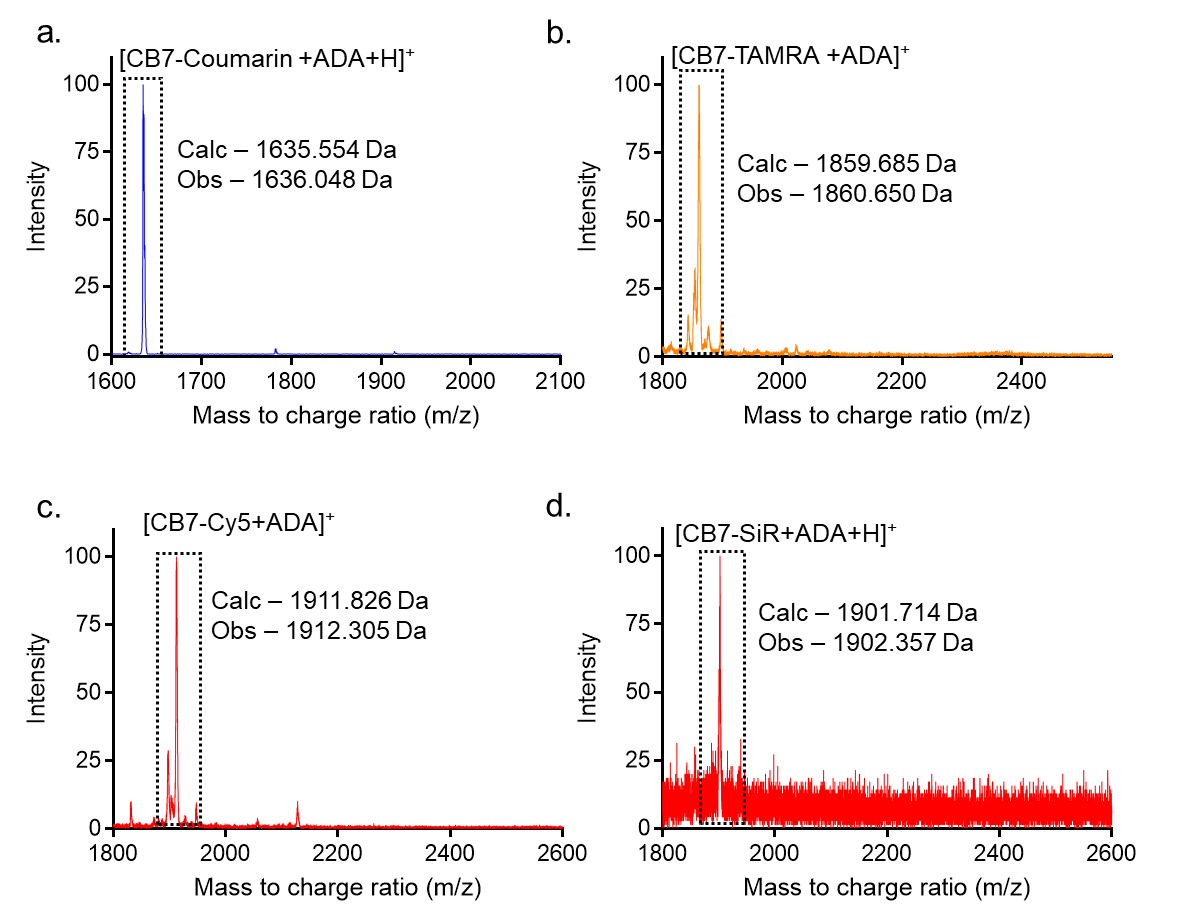

**Figure S3:** MALDI-MS analysis of ADA mediated displacement reaction (CB7-FL∙Xyl-Q + ADA) a) CB7-Coumarin∙Xyl-Dabcyl + ADA b) CB7-TAMRA∙Xyl-BHQ2 + ADA c) CB7-Cy5∙Xyl-BHQ3 + ADA d) CB7-SiR∙Xyl-BHQ3 + ADA. The complete disappearance of CB7-FL∙Xyl-Q complex mass and observation of CB7-FL∙ADA complex mass indicated a quantitative displacement reaction.

**13.4. Protocol for fluorescence quenching study of CB7–TAMRA with XYL–AuNP**

The fluorescence quenching titrations of CB7–TAMRA vs XYL–AuNP were performed using a fluorescence microplate reader. In this experiment, CB7– TAMRA was taken in a black 96-well microplate (Corning, non–binding plate) with a final concentration of 1 μM in 200 µl PBS buffer. Fluorescence spectra were recorded at respective excitation and emission wavelength scans listed in the table (Table S5a). After this, 0.005 eq of XYL– AuNP (final concentration 5.0 nM in 200 µl) was added to the system followed by the recording of fluorescence spectra. The addition of XYL– AuNP (0.005 eq in the first four steps, 0.01 eq in the subsequent three steps) was subsequently continued followed by spectra recording for the respective CB7– TAMRA to achieve maximum quenching. In the last two steps, the equivalent of XYL–AuNP is increased to reach the final concentration of 0.1 µM and 0.2 µM subsequently.

Similar protocols have been followed for quenching studies of EtA-TAMRA against XYL–AuNP (Figure S13 a).

**13.5. Protocol for fluorogenic response study from CB7–TAMRA∙XYL–AuNP quenched complex**

The CB7–TAMRA was taken in a black 96 well microplate (Corning, non–binding plate) with a final concentration of 1 μM in 200 µl PBS buffer. About 0.2 equivalent of XYL– AuNP was added to CB7–TAMRA to achieve maximum quenching of the fluorophore. After this, ADA–NH_2_ (final concentration 20.0 µM, 20.0 eq) was added to the quenched [CB7–TAMRA∙XYL–AuNP] systems, and fluorescence recovery was recorded after 10 minutes. The excitation and emission wavelengths used are listed in Table S5a.

**13.6. *In vitro* preparation and labeling of microtubules (MTs)**

The *in vitro* synthesized microtubules were inherently labeled with Alexa568 and biotin to examine the targeting specificity of the host-guest probe.

**Labeling reaction**

A warm Ti90 tube was filled with a 2.5 ml prewarmed high pH cushion. A polymerized MT mix was layered on top of the high pH cushion and spun at 43k rpm for 40 min at 35^o^C. The supernatant was then aspirated and the pellet was washed with warm labeling buffer. The pellet was then resuspended in 600 µl of warm labeling buffer (containing GTP). After subsequent washings, a total volume of 1 ml was transferred to a 2 ml centrifuge tube for labeling reaction. 20 µl of dye-NHS (Alexa568-NHS) was added to the resuspended MT mix and kept in a 37°C water bath. Mixed thoroughly by inversion. Vortexed gently and intermittently every 5 min for 20 min. Added remaining dye-NHS and repeated the above process. The reaction mix was then layered on 200 µl of prewarmed low pH cushion (containing GTP) in TLA 120.1 tubes and spun at 86k rpm for 20 min at 35°C. Pellet was washed with warm labeling buffer.

**Depolymerisation and polymerization cycle**

The pellet was resuspended in 100 µl of ice cold depolymerization buffer and placed on ice for 20 min. Spun at 80k rpm for 10 min at 4°C. To the supernatant, 5x BRB80, 2 mM MgCl_2,_ and 1mM GTP were added. It was then placed on ice for 5min. The polymerization was initiated by the addition of 350 µl prewarmed glycerol and it was allowed to polymerize for 30 min at 37°C. The polymerization mix was layered on 200 µl of prewarmed low pH cushion (with GTP) and spun at 80k rpm for 20 min at 35°C. Pellet was washed with warm BRB80 buffer. It was then resuspended in ice-cold BRB80 buffer. This was transferred to pre-cooled tubes and spun at 80k rpm for 10 min at 4°C. The supernatant was subsequently transferred to ice. The concentration of labeled tubulin was measured at 280 nm and aliquots were made. They were then snap-frozen and stored at -80^o^C until further use.

Buffers used:

1**.** BRB80 buffer

K- PIPES (pH – 6.8) – 80mM, MgCl_2_ – 2mM and EGTA – 1mM

2. 5x BRB80 buffer

K- PIPES (pH – 6.8) – 400mM, MgCl_2_ – 10mM and EGTA – 5mM

3. High pH cushion

HEPES (pH – 8.6) – 100mM, MgCl_2_ – 1mM, EGTA – 1mM, glycerol – 60%

4. Low pH cushion

5x BRB80 (pH – 6.8) – 1mM, glycerol – 60%

5. Labeling buffer

HEPES (pH – 8.6) – 100mM, MgCl_2_ – 1mM, EGTA – 1mM, glycerol – 40%

6. Depolymerisation buffer

K-glutamate (pH – 7.0) – 100mM, MgCl_2_ – 1mM

For the in-vitro synthesis of microtubules, 100 µM of tubulin heterodimers (α and β), 100 µM of labeled tubulin (Alexa568-NHS), 40 µM of biotin-conjugated tubulin, 1 mM of GMPCPP (nucleoside analog of GTP, with slower hydrolysis) were mixed in a microtubule polymerization buffer, BRB80 (80 mM PIPES buffer, 1 mM EGTA, 2 mM MgCl_2_, pH–6.8). This mix was then placed in ice for 5 min to depolymerize the tubulin oligomers if any. It was then spun down at 14k rpm for 10 min at 4^o^C to pellet down the protein aggregates. The supernatant which contains the unpolymerized tubulin was then placed for polymerization at 37^o^C for two hours. This was followed by pelleting of microtubules using a sucrose cushion which contains 50% sucrose in BRB80 buffer (spun down at 13k rpm for 10 min at 37^o^C to pellet down). The pellet was subsequently reconstituted in 1x BRB80 buffer and maintained at 37^o^C until further use.

**TIRF imaging protocol *of in vitro* prepared MTs**

For imaging of in vitro-microtubule labeling using CB[7]-ADA probe, a flow chamber was created using a clean microscopy slide, onto which two strips of double-sided adhesive tape is attached and a cover glass was placed on top. The flow cell was filled with 10 µl of BSA-Biotin (1 mg/ml) and incubated for 5 min, wherein BSA non-specifically interacts with the glass to facilitate immobilization of biotin on the surface. Washing was done to remove excess biotin by flowing in 1x BRB80 buffer twice through the chamber. Addition of 10 µl of streptavidin solution (0.5 mg/ml) for 5 min, which was then attached firmly to biotin on the surface. Unbound streptavidin in the channel was washed twice with 1x BRB80 buffer with casein (1.25 mg/ml) which blocks unbound sites and averts non-specific binding. Then, 20µl of an appropriate dilution of the MTs (with incorporated biotin-tubulin) with 5 µM tag ADA-taxol in 1x BRB80-casein buffer was flown through the chamber. This complex was incubated in the chamber for 10 min. Excess was washed off using 1x BRB80- casein buffer. CB[7]-Bodipy was diluted in 1x BRB80-casein buffer to a concentration of 500 nM. It was then flown (20µl) through the channel and incubated for 10 min. The final step involves washing the flow chamber using an anti-bleach mix of PCA, PCD, and Trolox in 1x BRB80-casein buffer (40µl). This was then imaged in the TIRF microscope under a 100x oil–immersion objective.

**13.7. Cell culture**

1. MEF/A431/3T3/HeLa cells were used for the experimental study.
2. The cells were cultured in a humidified atmosphere (5% CO_2_) at 37°C and grown in Dulbecco’s Modified Eagle’s Medium (DMEM, high glucose) supplemented with 10% fetal bovine serum (FBS) (Gibco, USA), 2 mM Glutamax (Invitrogen, USA) and 1% antibiotics (100 U/ml penicillin, 100 μg/ml streptomycin and 0.25 μg/ml amphotericin) (Gibco, USA).
3. At ~ 80% confluence, the cells were washed with DPBS (pH 7.3) (Gibco, USA), trypsinized, and suspended in a culture medium.
4. Cells were then counted and then in a typical experiment, ~10,000 cells/well/200µL media were plated in 96 well glass bottom plate (Eppendorf) / 35 mm glass-bottom cell imaging dish.
5. The cells were then maintained again in a humidified atmosphere (37°C, 5% CO_2_) for 24 h to reach ~60% confluence. Thereafter, cells were used for imaging experiments.

**13.8. Cell fixation, immunostaining and fluorogenic imaging protocol**

1. Culture media was removed from 96 well glass bottom plates and washed using 200 µL PBS two times.
2. Cells were fixed with chilled methanol for 7 minutes at –20°C followed by washing with PBS three times.
3. Blocking with 3% bovine serum albumin (BSA) in PBS at room temperature for 2 h.
4. Cells were incubated for 24 h at 4°C with primary antibody against microtubules (10 μg ml^−1^) diluted in PBS containing 3% bovine serum albumin.
5. Excess antibody was removed by three times washing with PBS (with 5 min incubation each time).
6. Cells were incubated with ADA conjugated secondary antibodies (10 μg ml^−1^) diluted in PBS containing 3% bovine serum albumin for 2 h.
7. Excess secondary antibody was removed by three times washing with PBS (with 5 min incubation each time).
8. [CB7–FL·XYL–Q] quenched probes (Final fluorophore concentration: 1 μM) were incubated with the ADA labeled cells and immediately proceeded for non–covalent fluorogenic imaging.

**13.9. Immunostaining and fluorogenic imaging protocol in live cell**

1. A431/3T3 cells (~10,000 in 200 μl culture media) were plated in the culture dish and allowed to reach up to 60% confluence.
2. Immunostaining of live cells was done after keeping the cells at ~4^°^C.
3. Culture media was removed carefully and cells were washed with 100 μl DPBS (pH 7.4) two times.
4. Cells were incubated with 100 μl primary antibody against EGFR (human, 10 μg ml^–1^) in DPBS for 30 min at 4^°^C.
5. Excess antibody was removed by washing the cells using 100 μl DPBS (pH 7.4) two times.
6. Then cells were incubated with 100 μl ADA conjugated donkey anti-human secondary antibody (10 μg.ml^–1^) in PBS for 30 min at 4^°^C.
7. Excess secondary antibody was removed by washing the cells using 100 μl DPBS (pH 7.4) two times.
8. [CB7–FL·XYL–Q] quenched probes (Final fluorophore concentration: 1 μM) were incubated with the ADA labeled cells and immediately proceeded for non–covalent fluorogenic imaging maintaining the live-cell imaging conditions.

**13.10. Protocol for dissection and labeling of actins in muscle tissue from *Drosophila melanogaster***

*Drosophila melanogaster* cultures maintained under 12 h light : 12 h dark cycles at 25^°^C were used. Adult flies were collected and kept on ice for around 15 minutes for anesthetizing.

**13.10.1. Thoracic muscle dissection**

1. After flies were anesthetized, they were submerged in PBS, placed dorsally on the dissection plate, and were pierced with insect pins on the abdomen region.
2. Using forceps, the exoskeleton of the thorax was incised carefully, peeled out gently and a bunch of clustered thoracic muscles was taken out.
3. Dissected muscle tissues were transferred in chilled PBS in labeled wells of the glass dish kept on ice and were allowed to settle.

**13.10.2. Fluorogenic imaging protocol for thoracic muscle tissue**

Tissues were fixed with 4% paraformaldehyde (PFA) at room temperature for 30 minutes with gentle shaking.

1. Tissues were washed at least thrice with 5 minutes of incubation using PBS containing 0.5% Triton X–100 (0.5% PBT).
2. Samples were then blocked using 10% horse serum in 0.5% PBT for 1 h at room temperature.
3. Phalloidin–ADA (2 µM) was added to each well and incubated for 2 h at room temperature.
4. Afterwards, samples were washed three times with 0.5% PBT for 5 min incubation each time.
5. [XYL–Q · CB7–FL] quenched probes (Final fluorophore concentration: 1 μM) were incubated with the ADA labeled tissues and immediately proceeded for non–covalent fluorogenic imaging.

**13.11. Dissection of intestine tissue from *Drosophila melanogaster* for live tissue imaging**

1. Third instar larvae of *Drosophila melanogaster* which were maintained in 12:12 hour light: dark and constant 25^ᵒ^C were scooped out from culture vials and kept on ice with PBS for around 15 minutes to anesthetize.

2. Individual larvae were submerged in PBS in a glass dish.

3. Using forceps, the mouth hooks of larvae were held firmly, while with the help of another pair of forceps the larva was held at about 3/4^th^ of the body length.

4. The posterior part of the body was gently pulled apart and separated from the anterior portion.

5. A small region of the gut corresponding to the midgut was isolated and placed in chilled PBS in a fresh well.

6. The gut was then incised with the help of fine scissors to expose and flatten the inner layer of tissue.

**13.12. Protocol for fluorogenic labeling of microtubules using ADA-taxol in live intestine tissue**

1. Dissected intestinal tissues were incubated with ADA-taxol conjugate (2 μM) in PBS medium for 90 min at room temperature.
2. Isolated tissues were washed twice with PBS with 1 min incubation each.
3. [XYL–Q · CB7–FL] quenched probes (Final fluorophore concentration: 1 μM in Schneider’s medium) were incubated with the ADA labeled tissues and immediately proceeded for non–covalent fluorogenic imaging using structured illumination microscopy method.

**13.13. Protocol for fluorogenic multiplexed imaging in fixed cells**

1. MEF cells were cultured and fixed using 4% PFA for 15 min at RT.
2. Fixature were removed and washed with 1x PBS three times.
3. Permeabilization using 0.25% (v/v) triton–X 100 in PBS 10 min at RT.
4. Cells were washed three times with 1x PBS.
5. Blocking using 3% BSA (w/v) in PBS for 2 h at RT.

4. Cells were incubated for 24 h at 4°C with primary antibody against microtubules (10 μg ml^−1^) diluted in PBS containing 3% bovine serum albumin and 0.1% Triton–X 100.
5. Excess antibody was removed by three times washing with PBS (with 5 min incubation each time).
6. Cells were incubated with ADA conjugated secondary antibodies (10 μg ml^−1^) diluted in PBS containing 3% bovine serum albumin and 0.5 μM Phalloidin–TCO conjugate for 2 h.
7. Excess secondary antibody and phalloidin were removed by three times washing with PBS (with 5 min incubation each time).
8. [CB7–TAMRA·Xyl-BHQ2] quenched probes (Final fluorophore concentration: 1 μM) and tetrazine–BODIPY probe (final concentration: 0.5 μM) were incubated with the ADA and TCO labeled cells for 5 min and then proceeded for fluorogenic multiplexed imaging of microtubules and actin.

**13.14. Protocol for fluorogenic labeling of microtubules using ADA-taxol in live HeLa cells**

1. HeLa cells (~10,000 in 200 μl culture media) were plated in the culture dish and allowed to reach up to 60% confluence.
2. Culture media was removed carefully and cells were washed with 100 µl DPBS (pH 7.4) two times.
3. Cells were incubated with ADA-taxol conjugate (10 µM) with verapamil hydrochloride (10 μM) for 4 h in complete cell culture DMEM media at 37^o^C.
4. Cells were then washed three times with DPBS (pH 7.4).
5. [CB7–TAMRA∙XYL–AuNP] quenched probes (CB7–TAMRA: 1 µM and XYL–AuNP: 200 nM) were incubated to the ADA labeled cells for 30 min in cell culture DMEM media (phenol red free) at 37^o^C.
6. After 30 min, non–covalent fluorogenic imaging is performed maintaining the live-cell imaging conditions (SIM and Confocal microscopy).

**13.15. Protocol for fluorogenic multiplexed imaging in live tissues**

1. After dissections, live intestine tissues were incubated with ADA-taxol (2 μM) and Jasplakinolide–TCO conjugate (0.5 μM) in PBS medium for 90 min at room temperature.
2. Isolated tissues were washed twice with PBS with 1 min incubation each.
3. [Xyl–BHQ2 · CB7–TAMRA] quenched probes (Final fluorophore concentration: 1 μM in Schneider’s medium) and tetrazine–BODIPY probe (final concentration: 0.5 μM) were incubated with the ADA and TCO labeled tissues for 5 min and then proceeded for fluorogenic multiplexed imaging.

**13.16. Protocol for control experiments in cells – Comparing ADA guest-modified targeting ligands to direct fluorophore-conjugated targeting agents**

**A. Microtubule target**

1. MEF cells were cultured and fixed using 4% PFA for 15 min at RT.
2. PFA was removed by washing with PBS three times.
3. Afterward, free aldehyde groups were reduced with NaBH_4_ (1 mg/ml) in PBS for 5 min.
4. After rinsing three times with PBS, cells were permeabilized using 0.25% (v/v) triton–X 100 in PBS for 10 min at RT.
5. Cells were washed three times with 1x PBS.
6. Blocking using 3% BSA (w/v) in PBS for 2 h at RT.

7. Cells were incubated for 24 h at 4°C with primary antibody against microtubules (10 μg ml^−1^) diluted in PBS containing 3% bovine serum albumin and 0.25 % Triton–X 100.
8. Excess antibody was removed by three times washing with PBS (with 5 min incubation each time).
9. Cells were incubated with a mixture of ADA conjugated secondary antibodies (10 μg ml^−1^) and Alexa647 conjugated secondary antibodies (3.33 μg ml^−1^) diluted in PBS containing 3% bovine serum albumin and 0.25 % Triton–X 100.
10. Excess secondary antibody was removed by three times washing with PBS (with 5 min incubation each time).
11. [CB7–Bodipy·XyL-BHQ1] quenched probe (Final fluorophore concentration: 1 μM) was incubated for 15 min and washed. After that, we proceeded with two-color SIM imaging of microtubules.

**B. Actin target**

1. MEF cells were fixed using the same above protocol (A).
2. Cells were incubated with a mixture of ADA-Phalloidin (1 μM) and Alexa488 conjugated phalloidin (0.33 µM) diluted in PBS containing 3% bovine serum albumin and 0.25 % Triton–X 100.
3. Excess Alexa488-Phalloidin and ADA-Phalloidin were removed by three times washing with PBS.
4. [CB7–Cy5·Xyl-BHQ3] quenched probe (Final fluorophore concentration: 1 μM) was incubated for 5 min and washed. After that, we proceeded with two-color SIM imaging of actin.

**14. Microscopy setup**

Epi–fluorescence microscopy via non–covalent fluorogenic labeling was carried out using an inverted microscope (Olympus) equipped with a cool–LED light source and a CMOS camera. Whereas, Structured illumination microscopy (SIM) was carried out using an inverted Zeiss ELYRA PS1 microscope equipped with 4 excitation lasers and an sCMOS camera. For epi–fluorescence microscopy of fixed cell and tissue samples, the imaging dishes were placed under the microscope, and images were captured in epi–fluorescence method using an Olympus oil–immersion objective (Plan–apochromat DIC 63x/1.40 Oil). Four LED sources have been used for excitation: 365 nm, 490 nm, 550 nm, and 635 nm for the respective excitation of fluorophores. Fluorescence light was spectrally filtered with – U-FUNA (Exciter filter BP 360-370 nm, Dichroic beamsplitter DM410 with barrier filter BA420-460) for excitation of 365nm, U-FBNA (Exciter filter BP 470-495 nm, Dichroic beamsplitter DM505 with barrier filter BA510-550) for excitation of 490 nm, U-FGWA (Exciter filter BP 530-550 nm, Dichroic beamsplitter DM570 with barrier filter BA575-625) for excitation of 550 nm, and Brightline Quad‐band “Pinkel” Filter set (LED‐DA/FI/TR/Cy5‐4X‐B‐000) for excitation of 635 nm. Imaging was carried out using an optiMOS sCMOS camera (QImaging).

Live tissue samples kept in microscope imaging dishes were placed under the microscope maintained at 37°C and 5% CO_2_ atmosphere and fluorescence microscopic images were captured by structured illumination method using an inverted Zeiss ELYRA PS1 microscope. Four lasers have been used for excitation: 405 nm (50 mW), 488 nm (200 mW), 561 nm (200 mW), and 642 nm (150 mW) for respective excitation of fluorophores. Details of imaging parameters for the individual experiments have been given in the table below. Imaging was performed using a Zeiss oil–immersion objective (alpha Plan–apochromat DIC 63x/1.40 Oil DIC M27, numerical aperture (NA) 1.40 oil). Fluorescence light was spectrally filtered with emission filters (MBS– 405+EF BP 420–495/LP 750 for laser line 405, MBS– 488+EF BP 495–570/LP 750 for laser line 488, MBS– 561+EF BP 570–650/LP 750 for laser line 561 and MBS–642+EF LP 655 for laser line 642) and imaged using a PCO edge sCMOS camera.

For confocal imaging, cells were imaged using a Leica TCS SP8 microscope. Two lasers were used for the experiments: 405 nm (source: 50 mW) and 552 nm (source: 20 mW). Imaging was performed using Leica oil–immersion objectives: HC PL APO CS2 63x with a numerical aperture (NA) 1.40. Excitation beam splitter – DD 488/552. Cells were imaged with a HyD detector (408nm-466 nm) – Gain 100 and a PMT detector (569nm-651nm) – Gain 750. Confocal images were processed using LAS X (Leica) and ImageJ software.

**Table S7**: Table for imaging parameters in SIM experiment.

| **Lasers** | **Fluorophores used** | **Exposure time (ms)** | **Laser power density (W.cm^–2^)** |
| --- | --- | --- | --- |
| 405 | Coumarin | 100 | 1.22 |
| 488 | Fluorescein, BODIPY | 100 | 6.51 |
| 561 | Cy3, TMR | 100 | 7.32 |
| 642 | Cy5, SiR | 100 | 2.93 |

**Table S8**: Table for imaging parameters of Epi-fluorescence microscopy for Figure 2 – 5.

| **LED Source** | **Fluorophores used** | **Exposure time (ms)** | **LED intensity (Percentage, %)** |
| --- | --- | --- | --- |
| 365 nm | Coumarin, DAPI | 100 | 25 |
| 490 nm | Fluorescein, BODIPY | 100 | 20 |
| 550 nm | Cy3, TMR | 100 | 20 |
| 635 nm | Cy5*, SiR | 100 | 20 |

* In the case of imaging using Cy5 and DAPI channel, the DAPI parameter used is 25% LED intensity and 40 ms exposure time.

**15. Image processing and data analysis**

Epi–fluorescence microscopy images of fixed cells were used as acquired after adjusting the brightness using ImageJ and data analysis was carried out using GraphPad prism. Other epifluorescence images from fixed tissues and live cells were subjected for deconvolution using cellsens software (Olympus) pre-installed with constrained iterative deconvolution mathematical algorithm function. Structured illumination image was reconstructed using a structured illumination analysis package for Zen 2.0 software (Zeiss). Additional software has been used for color adjustment (ImageJ) and data analysis (Origin 9.0 and GraphPad Prism).

**Table S9**: Experimental set-up for imaging of cells in Figures 2 & 3.

| Figures | Mode of imaging | Targeting molecules (TM) | | Cell/Tissue (Protocol) | Exposure time |
| --- | --- | --- | --- | --- | --- |
|  |  | TM – 1 | TM – 2 |  |  |
| Figure 2c & 3 | Epi-fluorescence microscopy | α-tubulin (rat) | ADA conjugated donkey anti-rat | MEF cell (SI – 13.8) | 100 ms |

**Table S10**: Experimental set-up for imaging of tissues in Figure 4.

| Figures | Mode of imaging | Targeting molecules (TM) | Cell/Tissue (Protocol) | Exposure time |
| --- | --- | --- | --- | --- |
| Figure 4a | SIM | ADA-Phalloidin | MEF cell (SI – 13.16) | 100 ms |
| Figure 4b | Epi-fluorescence microscopy | ADA-Phalloidin | Thoracic muscle tissue (SI – 13.10) | 100 ms |

**Table S11**: Experimental set-up for imaging of cells and tissues in Figure 5.

| Figures | Mode of imaging | Targeting molecules (TM) | | Cell/Tissue (Protocol) | Exposure time |
| --- | --- | --- | --- | --- | --- |
|  |  | TM – 1 | TM – 2 |  |  |
| Figure 5b | Epi-fluorescence microscopy | Human EGFR Antibody | ADA conjugated donkey anti-human | A431 cell (SI – 13.9) | 100 ms |
| Figure 5d | SIM | ADA-taxol | | In-vitro microtubules (SI – 13.6) | 100 ms |
| Figure 5g | Confocal | ADA-taxol | | Live cell microtubules (SI – 13.14) |  |
| Figure 5h | SIM | ADA-taxol | | Live intestine tissue (SI – 13.12) | 100 ms |

**Table S12**: Experimental set-up for imaging of cells and tissues in Figure 6.

| Figures | Mode of imaging | Targeting molecules (TM) | | Cell/Tissue (Protocol) | Exposure time |
| --- | --- | --- | --- | --- | --- |
|  |  | TM – 1 | TM – 2 |  |  |
| Figure 6b | SIM | 1. α-tubulin (rat) | ADA conjugated donkey anti-rat | MEF cell (SI – 13.13) | 100 ms |
|  |  | 2. Phalloidin-TCO | |  |  |
| Figure 6c | SIM | 1. ADA-taxol | 2. Jasplakinolide–TCO | Live intestine tissue (SI – 13.15) | 100 ms |

**15.1. Protocol for kinetic study in the cell using confocal imaging**

MEF cells were fixed and labeled with ADA-antibody using the protocol SI – 13.8. Two experiments were performed with – 1. Fixed MEF cells without any immunostaining and 2. Fixed MEF cells labeled with ADA-antibody. Next, cells were incubated with quenched probe [CB7-TAMRA (1 µM) + XYL-BHQ2 (1.5 µM)] in both cases. The imaging was performed using a confocal microscope with a time interval of 3 min for only MEF cells and 1 min for ADA-antibody labeled cells over a duration of 30 min (Figure S4 and S6).

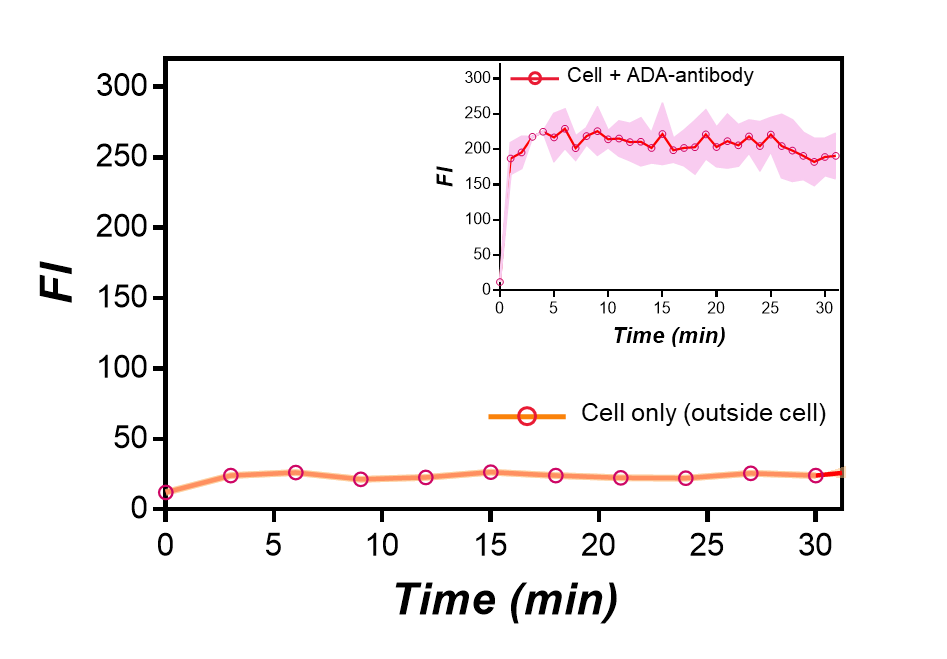

**Figure S4:** Time-lapse fluorescence study using only cells (without ADA labeling), which shows that the intensity of the cellular medium does not change upon incubation with the host-guest quenched ([CB7–TAMRA·XYL–BHQ2]) probe. This confirms the stability of the quenched probes in cellular conditions. As a comparison, the inset shows the kinetics of fluorogenic labeling using quenched probes [CB7–TAMRA·XYL–BHQ2] in ADA immunolabeled cells. The probe concentration was maintained as described in Table S6.

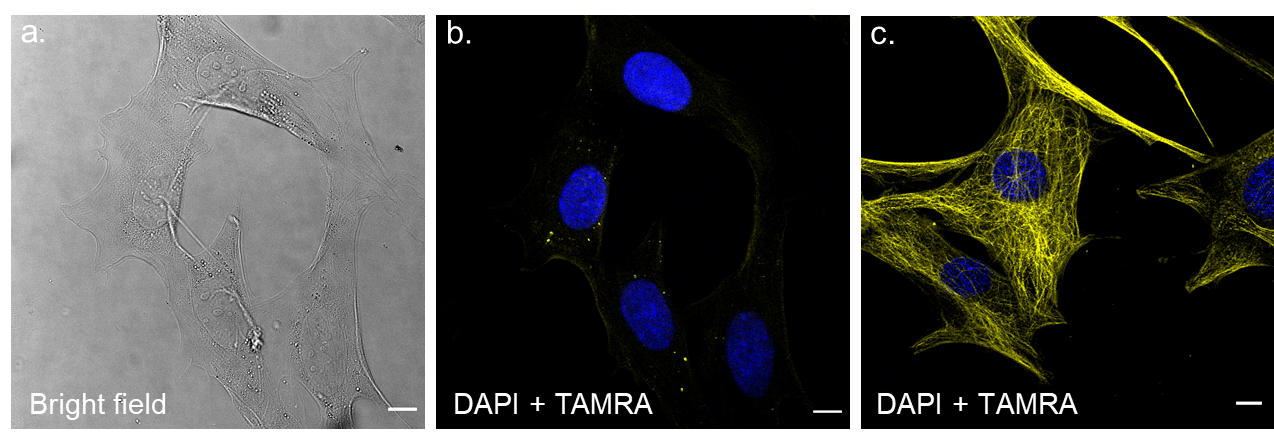

**Figure S5:** (a-b) Images of control cells (without ADA labeling) upon incubation with the host-guest quenched ([CB7–TAMRA·XYL–BHQ2]) probe. a) Brightfield and b) Fluorescence channel (TAMRA + DAPI). Negligible fluorescence signals from the cells indicated minimal off-target activation and labeling from the host-guest quenched probe. As a comparison, (c) shows an example of fluorogenic labeling using [CB7–TAMRA·XYL–BHQ2] in ADA labeled cells. MEF cells are labeled using protocol SI – 13.8. Imaging was performed using a confocal microscope (Source – 405 nm for DAPI and 552 nm for TAMRA). Gain – 100, Laser intensity – 2% for DAPI and Gain – 750, and Laser intensity – 1% for TAMRA. Scale bar – 10 µm. The probe concentration was maintained as described in Table S6.

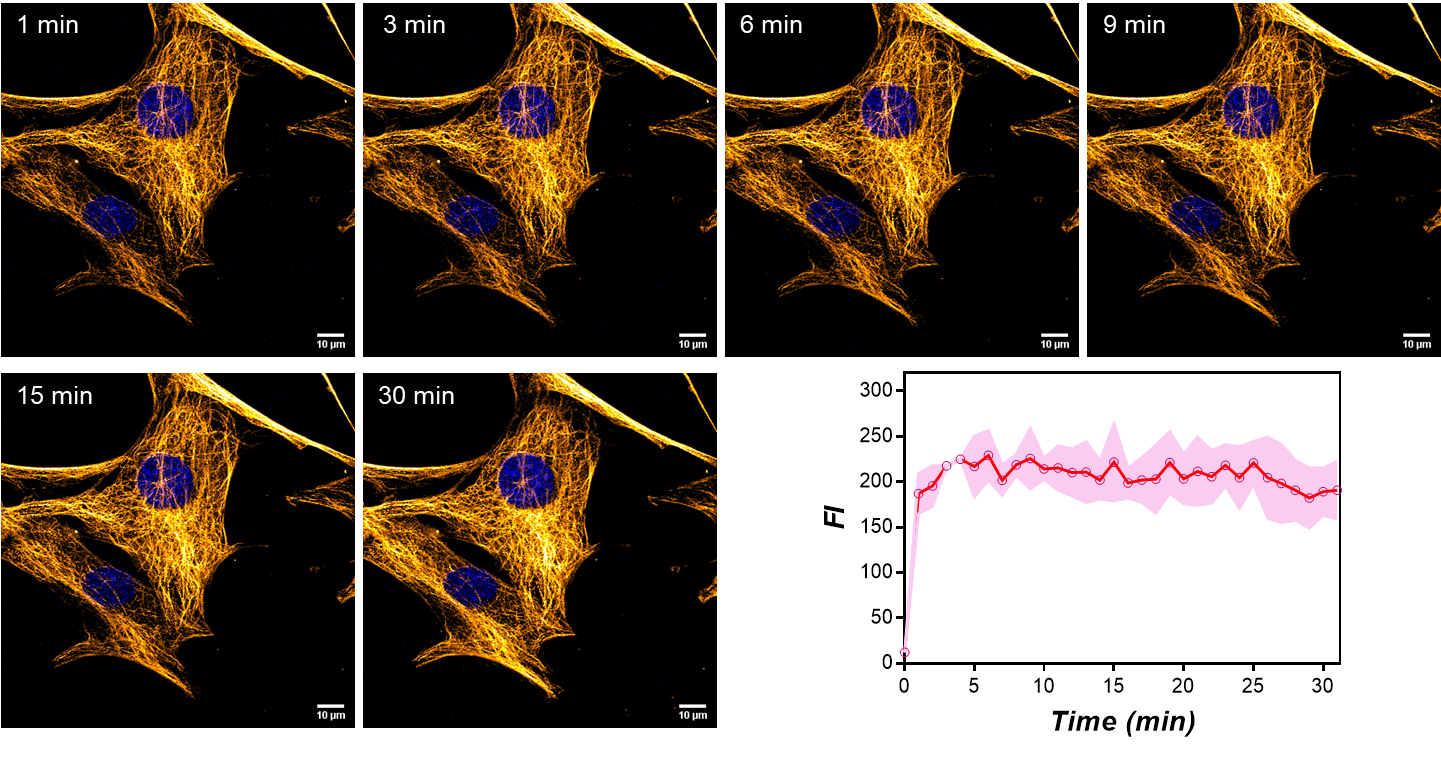

**Figure S6:** Kinetics of fluorogenic labeling using quenched probes in ADA labeled cells. Antibody-based ADA cells were treated with [CB7–TAMRA·XYL–BHQ2] quenched probe and time-lapsed images were immediately recorded to demonstrate the temporal intensity improvement from microtubule targets, which in turn exhibits the labeling kinetics on the imaging platform. The fluorescence intensity plot over time indicated the completion of the labeling within minutes. MEF cells are labeled using protocol SI – 13.8. Imaging was performed using a confocal microscope (Source – 405 nm for DAPI and 552 nm for TAMRA). Gain – 100, Laser intensity – 2% for DAPI and Gain – 750, and Laser intensity – 1% for TAMRA. Scale bar – 10 µm. The probe concentration was maintained as described in Table S6.

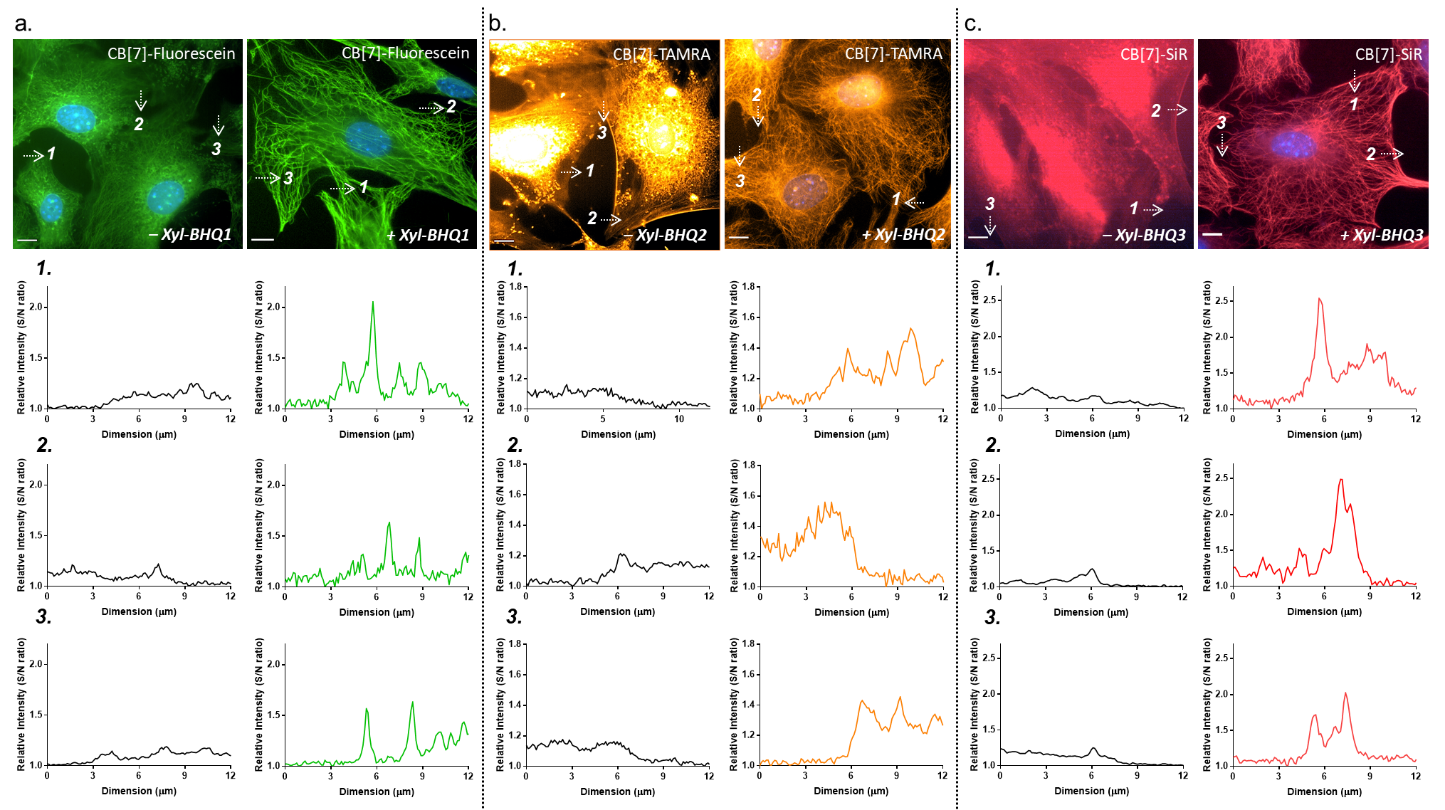

**Figure S7:** Intensity profile reporting single to noise (S/N) ratio for the fluorogenic imaging (CB7-FL·Xyl-Q) as compared to only fluorophore-based imaging (CB7-FL) in ADA-labeled cells. a) CB7-Fluorescein with and without Xyl-BHQ1, b) CB7-TAMRA with and without Xyl-BHQ2, and c) CB7-SiR with and without Xyl-BHQ3. The intensity profile was drawn along the arrows as shown in the images. Scale bar – 10 µm. MEF cells are labeled using protocol SI – 13.8 [The probe concentration was maintained as described in Table S6]. Imaging parameters are used as described in Table S8 (LED source – 365 nm for DAPI, 490 nm for Fluorescein, 550 nm for TAMRA, and 635 nm for SiR).

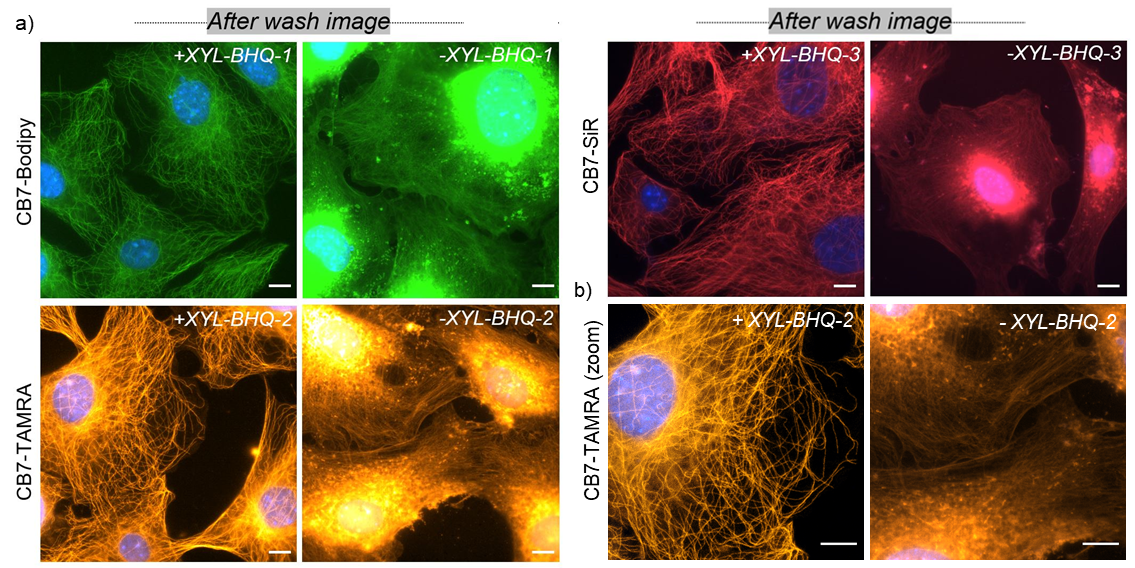

**Figure S8:** a) Fluorescence images acquired after washing excess probes demonstrate reduced off-target signal from the host-guest quenched probes due to its conditionally activable nature whereas, the absence of quenchers resulted in off-target binding of imaging probes. b) Zoomed image. Scale bar:10 μm (a–b). MEF cells are labeled using protocol SI – 13.8 [ The probe concentration was maintained as described in Table S6]. Imaging parameters are used as described in Table S8 (LED source – 365 nm for DAPI, 490 nm for Bodipy, 550 nm for TAMRA, and 635 nm for SiR).

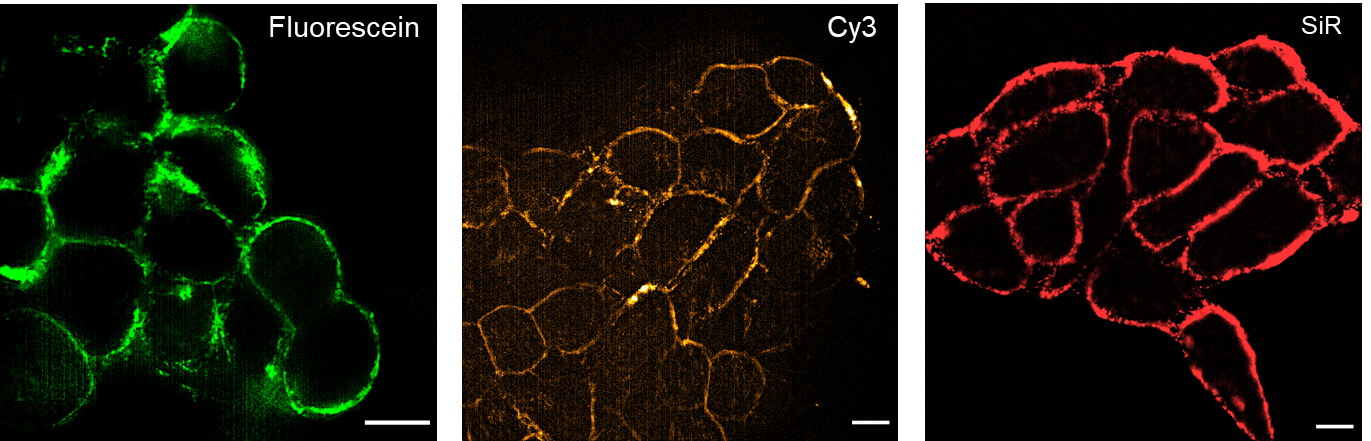

**Figure S9:** Additional images (with fluorescein, Cy3, and SiR host-guest probes) showing fluorogenic imaging of EGFR in live A431 cells. These images demonstrate the capability of fluorescein, Cy3, and SiR probes for supramolecular host-guest fluorogenic imaging in live cells. Scale bar – 10 µm. A431 cells are labeled using protocol SI – 13.9 [The probe concentration was maintained as described in Table S6]. Imaging parameters are used as described in Table S8 (LED source – 490 nm for Fluorescein, 550 nm for Cy3, and 635 nm for SiR).

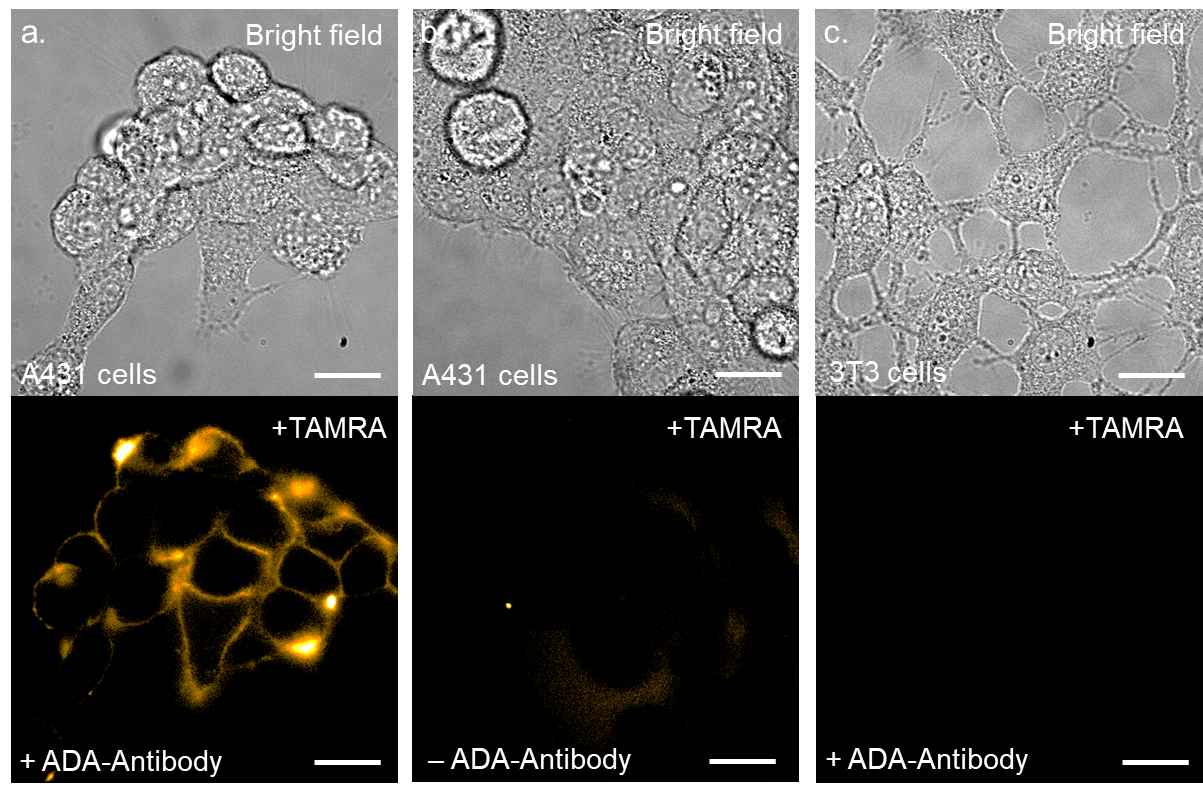

**Figure S10:** Fluorogenic imaging of EGFR and control experiments using [CB7–TAMRA·XYL–BHQ2] probe. a) EGFR labeling using ADA-Antibody in A431 cells. b) A431 cells without ADA-Antibody treatment show no EGFR labeling. c) EGFR negative 3T3 cells shows no fluorescence signal even with ADA-Antibody treatment [ CB7-TAMRA: 1.0 µM and Xyl-BHQ2 – 1.0 µM]. Scale bar – 20 µm. A431 and 3T3 cells are labeled using protocol SI – 13.9. Excitation used – 550 nm for TAMRA having 20% intensity (LED source) with 30 ms exposure time.

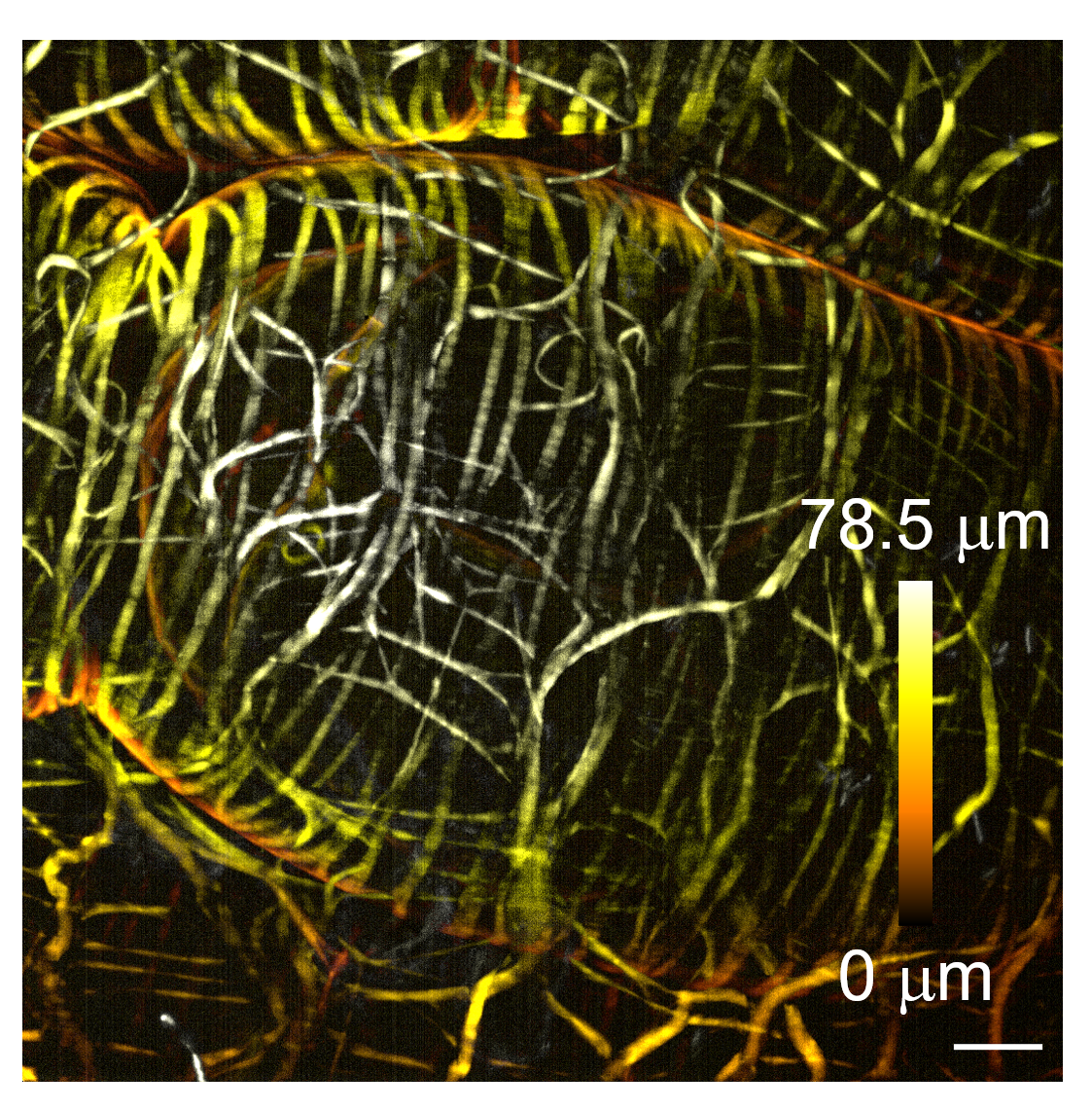

**Figure S11:** 3D fluorogenic imaging of actins over 80 µm Z depth from ovary tissues of *Drosophila melanogaster.* Scale bar (XY): 10 µm. The probe concentration for CB7-TAMRA·XyL-BHQ2 is maintained as described in Table S6.

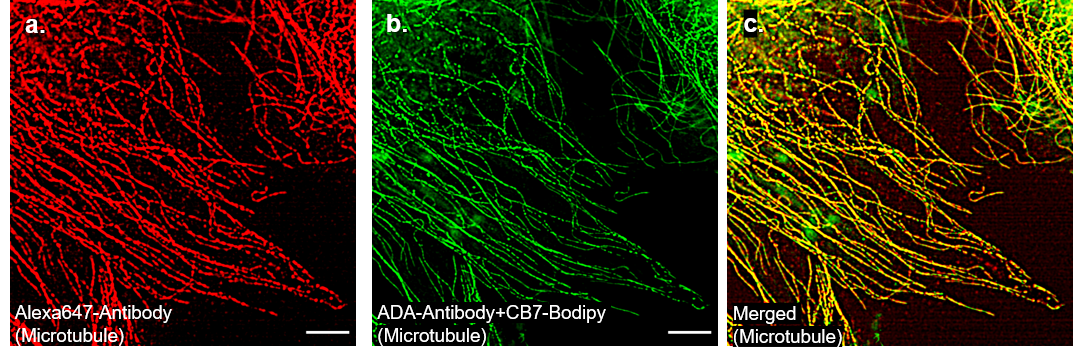

**Figure S12:** Control experiment comparing ADA guest-modified antibodies to direct fluorophore-conjugated antibodies. (a-c) Two-color SIM imaging of microtubules in MEF cells co-stained with two different antibody conjugates. a) SIM image of the microtubules targeted using Alexa647-antibody (642 nm channel). b) SIM image of the microtubules targeted using ADA-antibody. CB7–Bodipy·XYL–BHQ1 complex is used to carry out the fluorogenic imaging of microtubules in the 488 nm channel (after washing). c) Merged image of microtubules. Scale bar – 5 µm. MEF cells are fixed and labeled using protocol SI – 13.16.

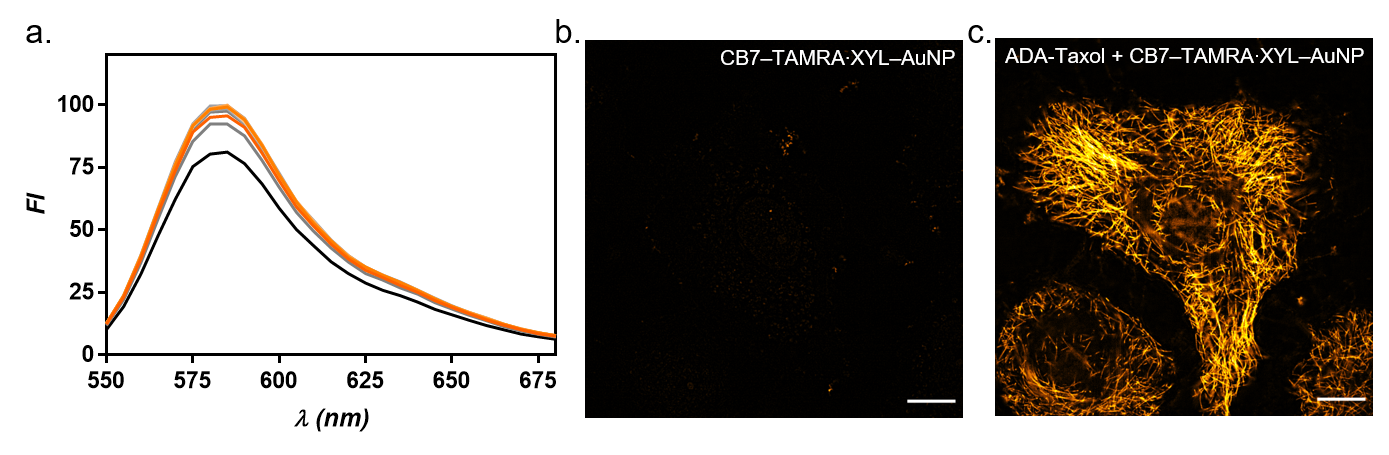

**Figure S13:** (a) Fluorescence titration data for Ethanolamine–TAMRA (EtA-TAMRA) vs XYL-AuNP. Negligible fluorescence quenching from the non–interacting pair demonstrated that the fluorescence quenching requires specific recognition of CB7 host and XYL guest moiety. (b) SIM imaging of microtubules in live HeLa cells using CB7-FL∙XYL-AuNP quenched probes without ADA-Taxol. (c) Fluorogenic SIM imaging of microtubules in live HeLa cells using ADA-Taxol and CB7-FL∙XYL-AuNP quenched probes. Scale bar: 10 µm.

**16. HPLC, MALDI, and NMR characterization data**

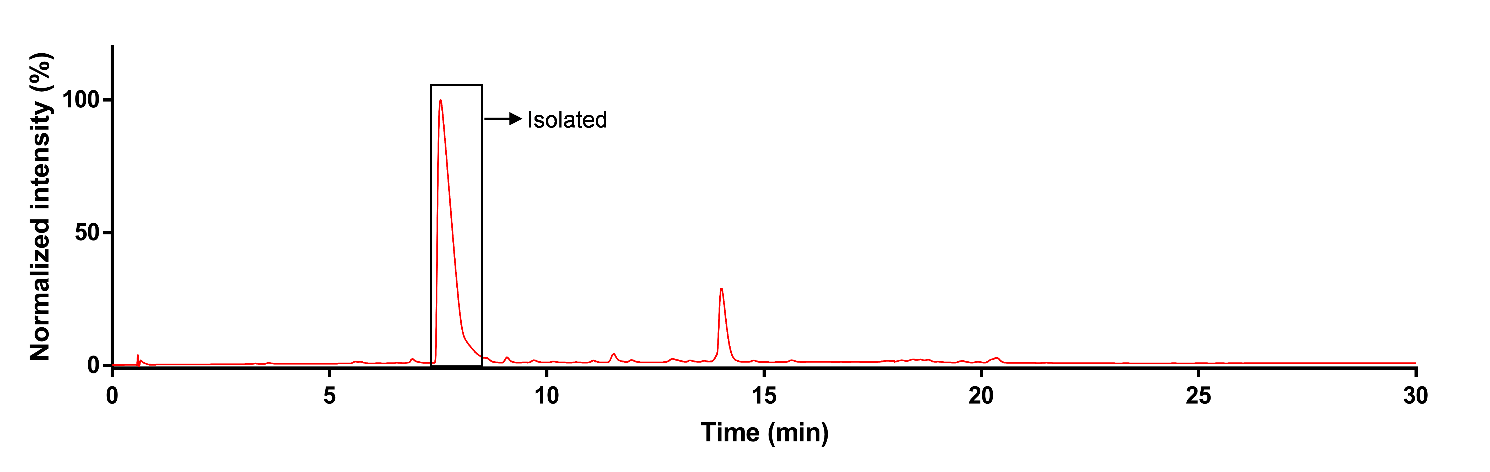

**Figure S14:** HPLC chromatogram of XYL–Dabcyl conjugate. The polarity of acetonitrile was varied from 5 to 100% in 30 min. The conjugated product was isolated at retention time (R_t_) 8.1 min.

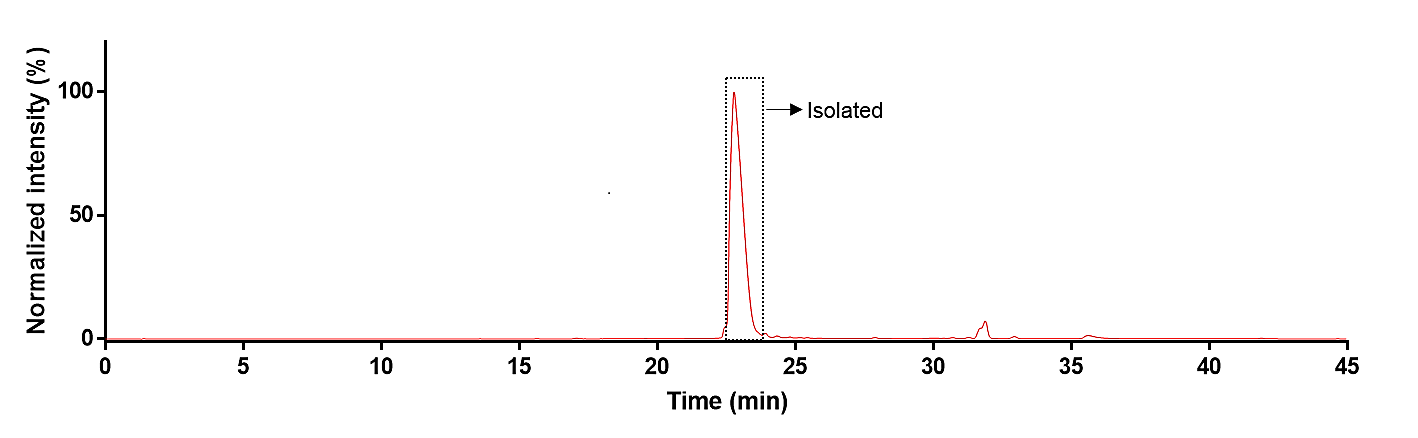

**Figure S15:** HPLC chromatogram of XYL–BHQ1 conjugate. The polarity of acetonitrile was varied from 5 to 100% in 45 min. The conjugated product was isolated at retention time (R_t_) 23.6 min.

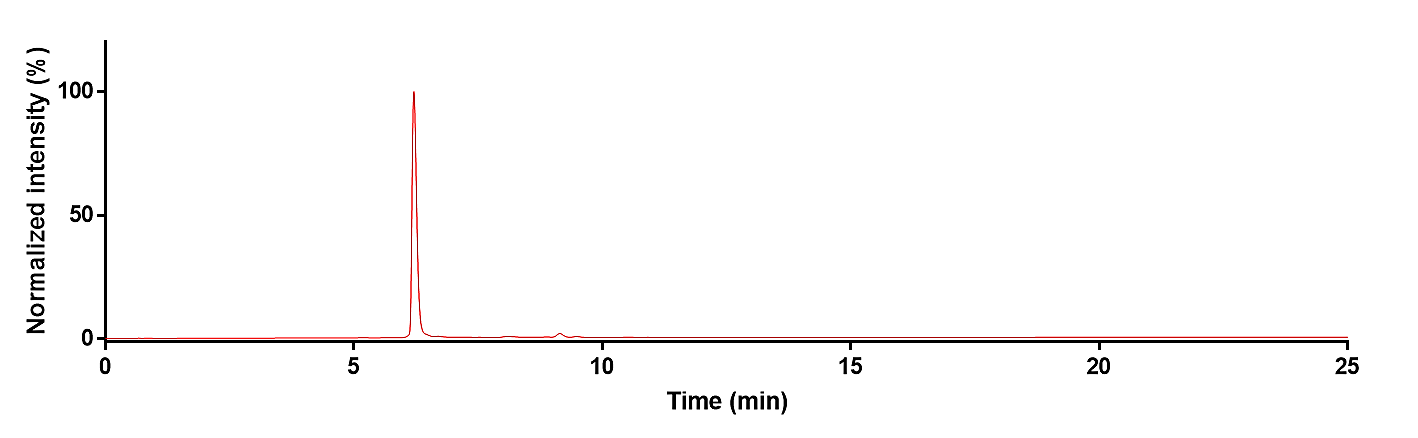

**Figure S16:** HPLC chromatogram of XYL–BHQ2 conjugate. The polarity of acetonitrile was varied from 5 to 100% in 25 min. The conjugated product was isolated at retention time (R_t_) 6.9 min.

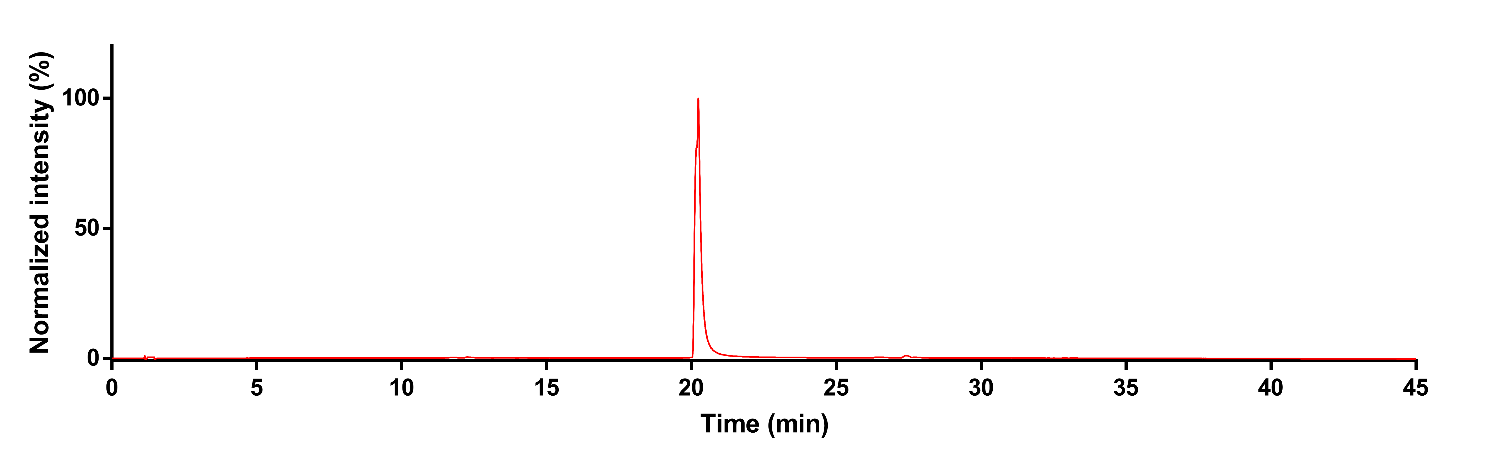

**Figure S17:** HPLC chromatogram of XYL–BHQ3 conjugate. The polarity of acetonitrile was varied from 5 to 100% in 45 min. The conjugated product was isolated at retention time (R_t_) 20.5 min.

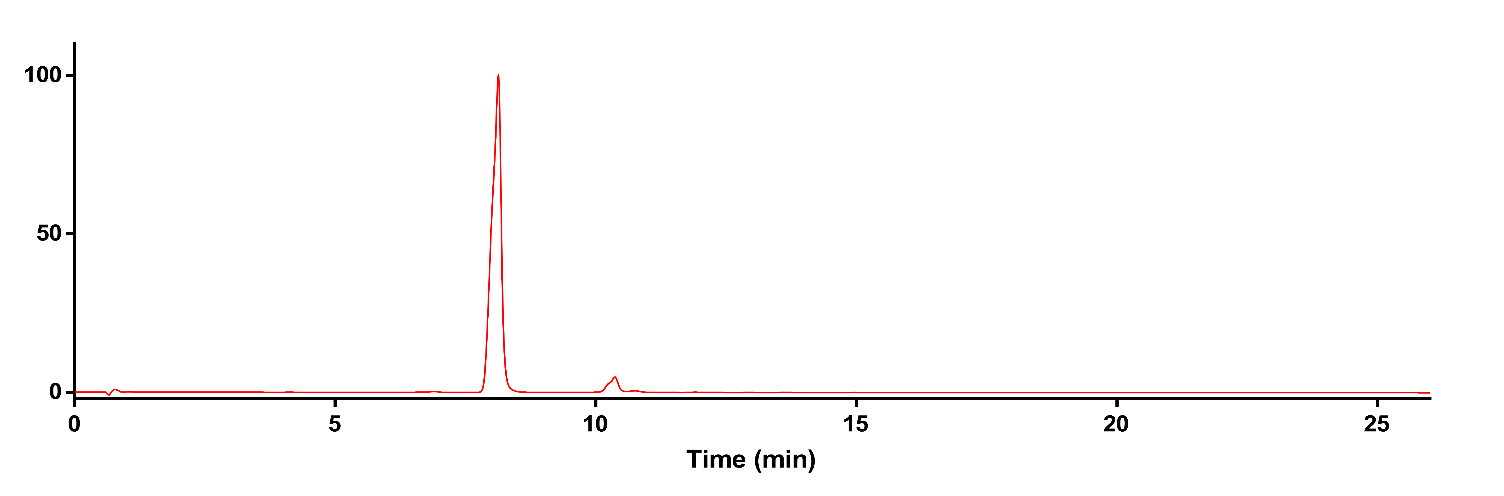

**Figure S18:** HPLC chromatogram of EtA–Dabcyl conjugate. The polarity of acetonitrile was varied from 5 to 80% in 25 min and then upto 100% in 26 min. The conjugated product was isolated at retention time (R_t_) 8.1 min.

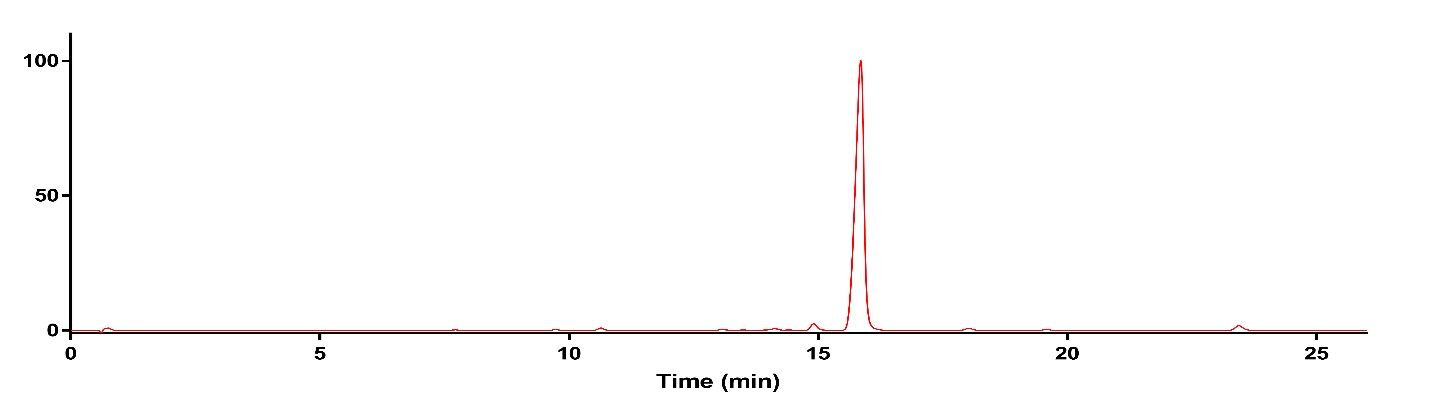

**Figure S19:** HPLC chromatogram of EtA–BHQ1 conjugate. The polarity of acetonitrile was varied from 5 to 80% in 25 min and then upto 100% in 26 min. The conjugated product was isolated at retention time (R_t_) 15.5 min.

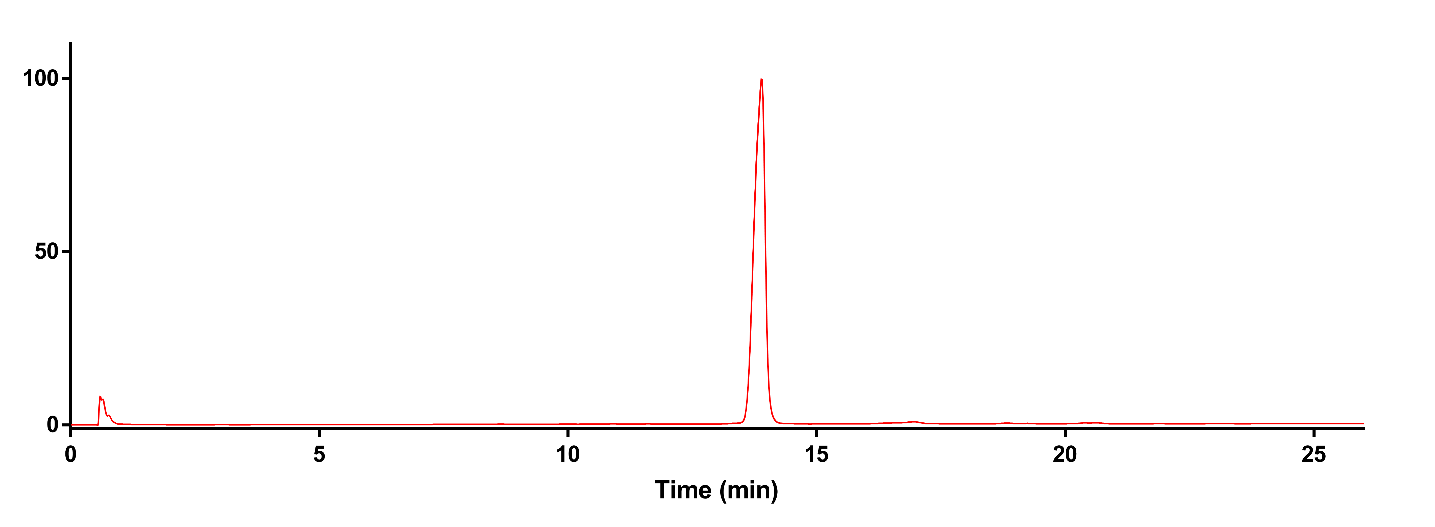

**Figure S20:** HPLC chromatogram of EtA–BHQ2 conjugate. The polarity of acetonitrile was varied from 5 to 80% in 25 min and then upto 100% in 26 min. The conjugated product was isolated at retention time (R_t_) 14.6 min.

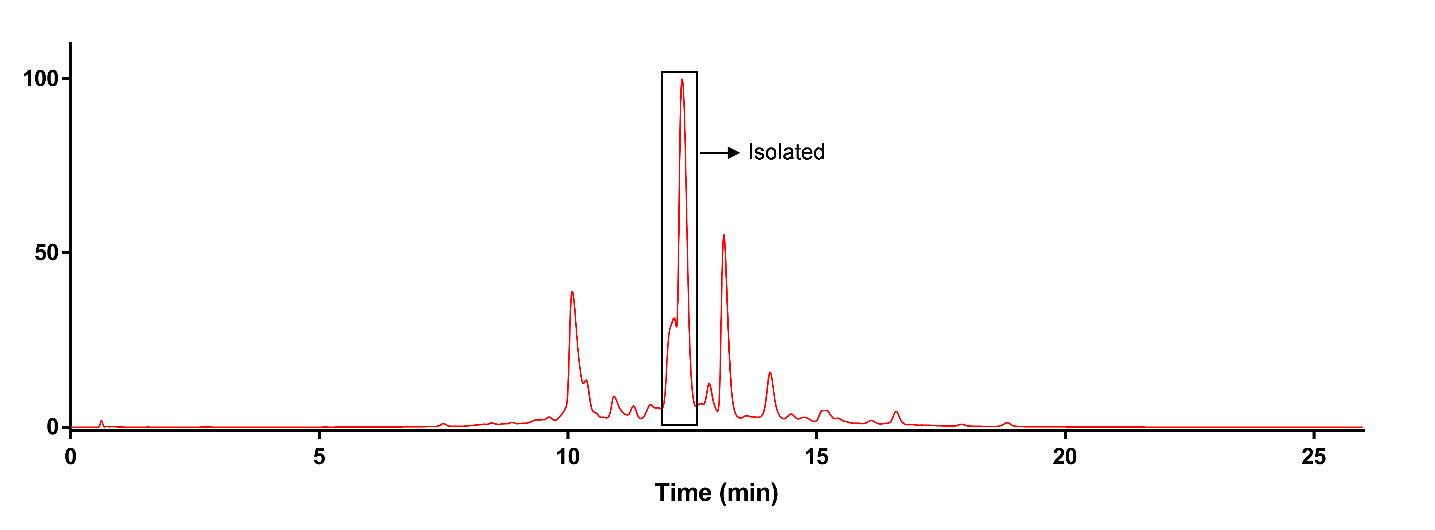

**Figure S21:** HPLC chromatogram of EtA–BHQ3 conjugate. The polarity of acetonitrile was varied from 5 to 80% in 25 min and then upto 100% in 26 min. The conjugated product was isolated at retention time (R_t_) 12.6 min.

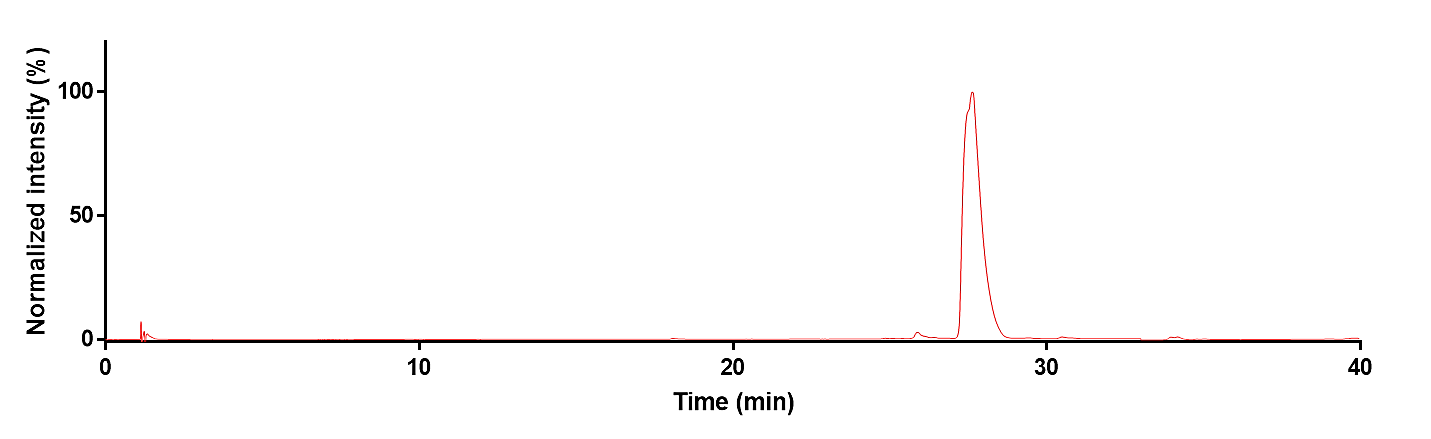

**Figure S22:** HPLC chromatogram of ADAPc–PEG–NHS ester. The polarity of acetonitrile was varied from 5 to 100% in 40 min. The conjugated product was isolated at retention time (R_t_) 28.5 min.

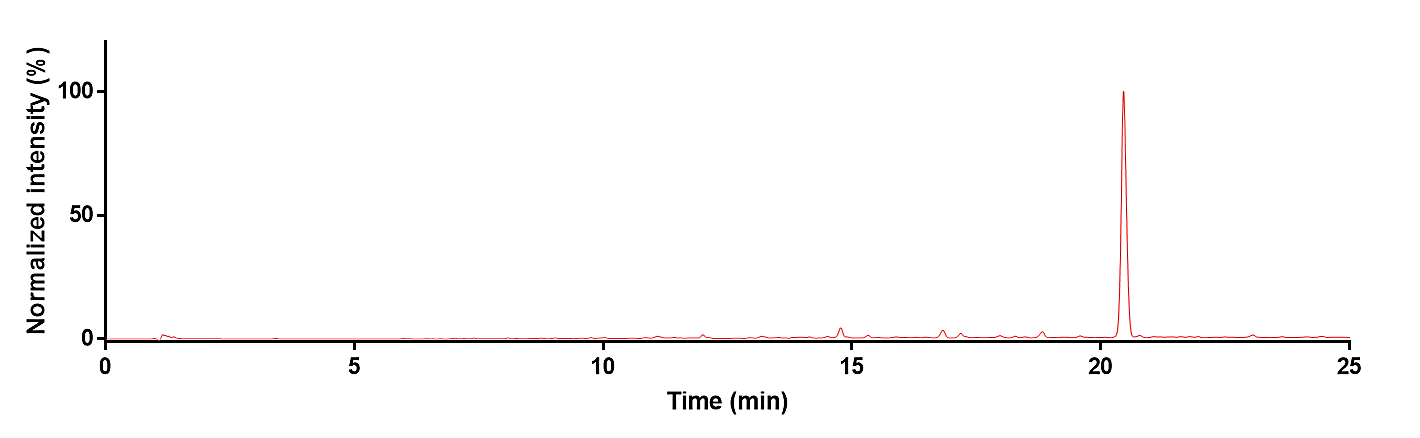

**Figure S23:** HPLC chromatogram of ADA conjugated taxol. The polarity of acetonitrile was varied from 5 to 100% in 25 min. The conjugated product was isolated at retention time (R_t_) 20.5 min.

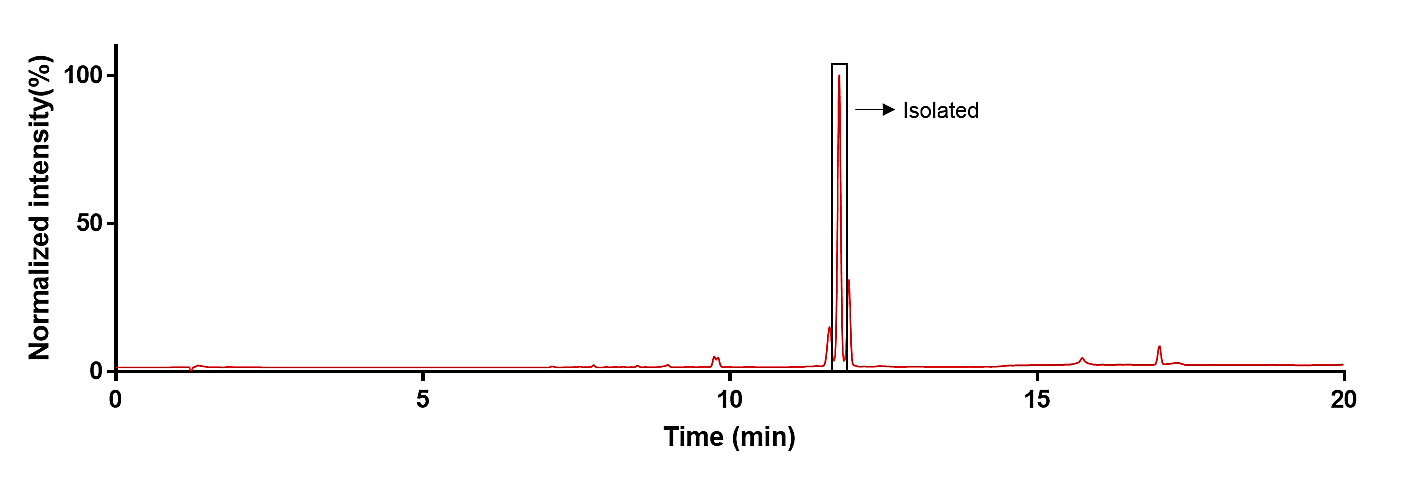

**Figure S24:** HPLC chromatogram of Phalloidin–TCO conjugate. The polarity of acetonitrile was changed from 5% to 50% in 15 min and then to 100% in 20 min. The conjugated product was isolated at retention time (R_t_) 13.5 min.

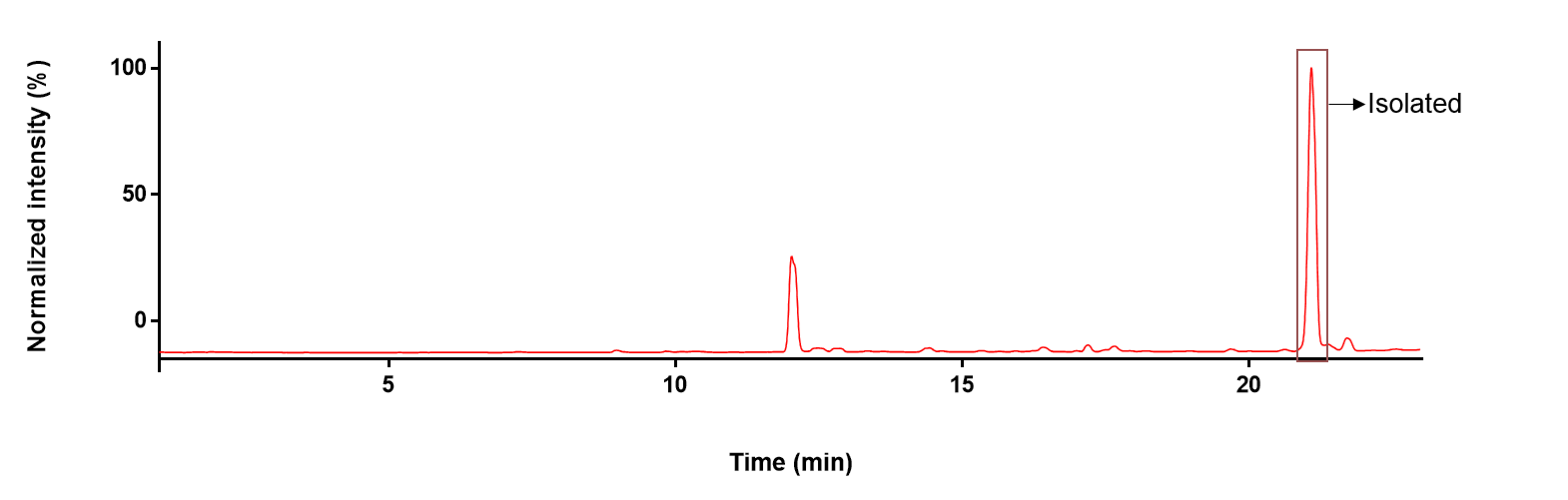

**Figure S25:** HPLC chromatogram of Jasplakinolide–TCO conjugate. The polarity of acetonitrile was changed from 5% to 70% in 20 min and then to 100% in 22 min.

**16.1. Protocol for MALDI-MS analysis of antibody**

Antibodies (1 µL) after desalting through Zeba spin column (~ 1 mg.mL^–1^ in MiliQ water) were taken in a microcentrifuge tube containing 1 µL of sinapinic acid [10 mg.mL^–1^ in 50:50 water (0.1% TFA/acetonitrile)]. The mixed solution was placed onto a MALDI plate and allowed to dry at room temperature for analysis.

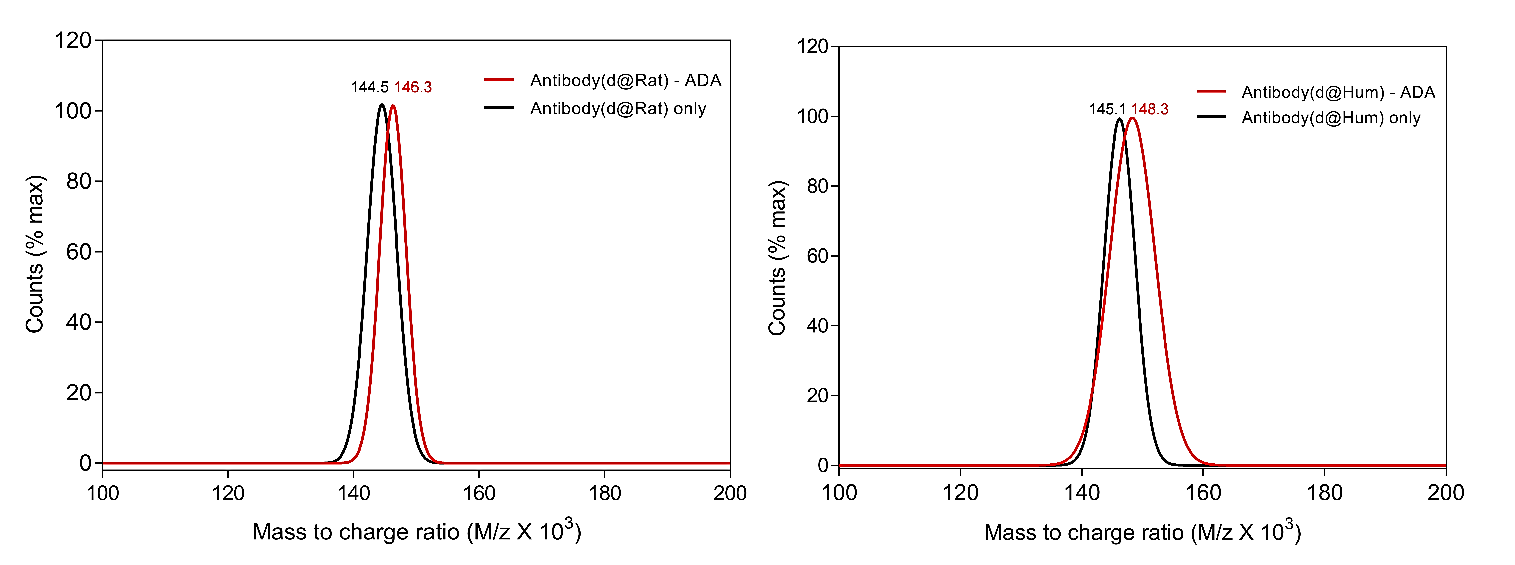

**Figure S26:** MALDI-MS analysis of ADA conjugated rat and human secondary antibodies. The number of ADA moieties anchoring to the antibodies was calculated to be two and three for rat and human antibodies respectively.

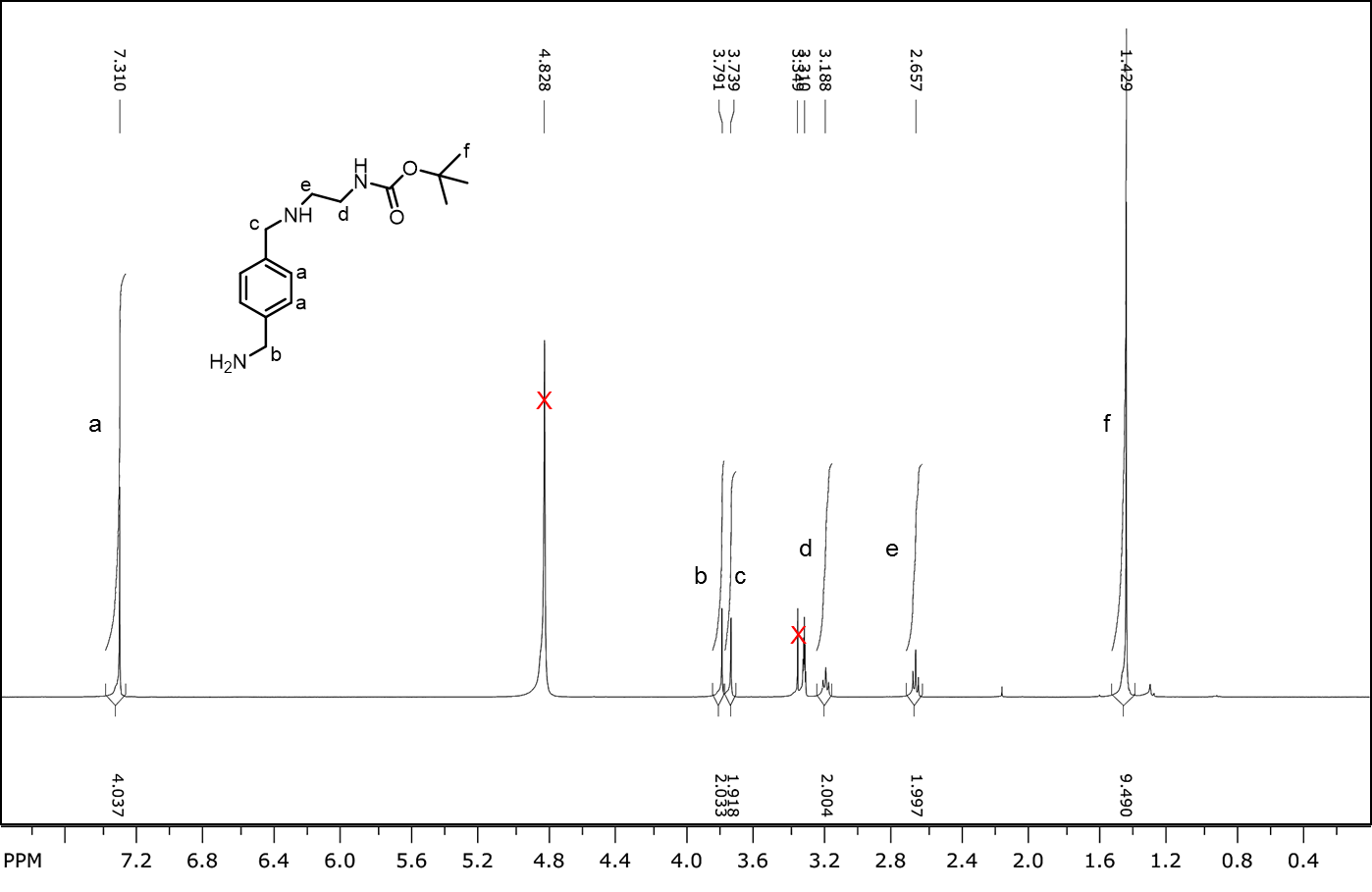

**Figure S27:** ^1^H NMR of compound 2 (400 MHz, CD_3_OD).

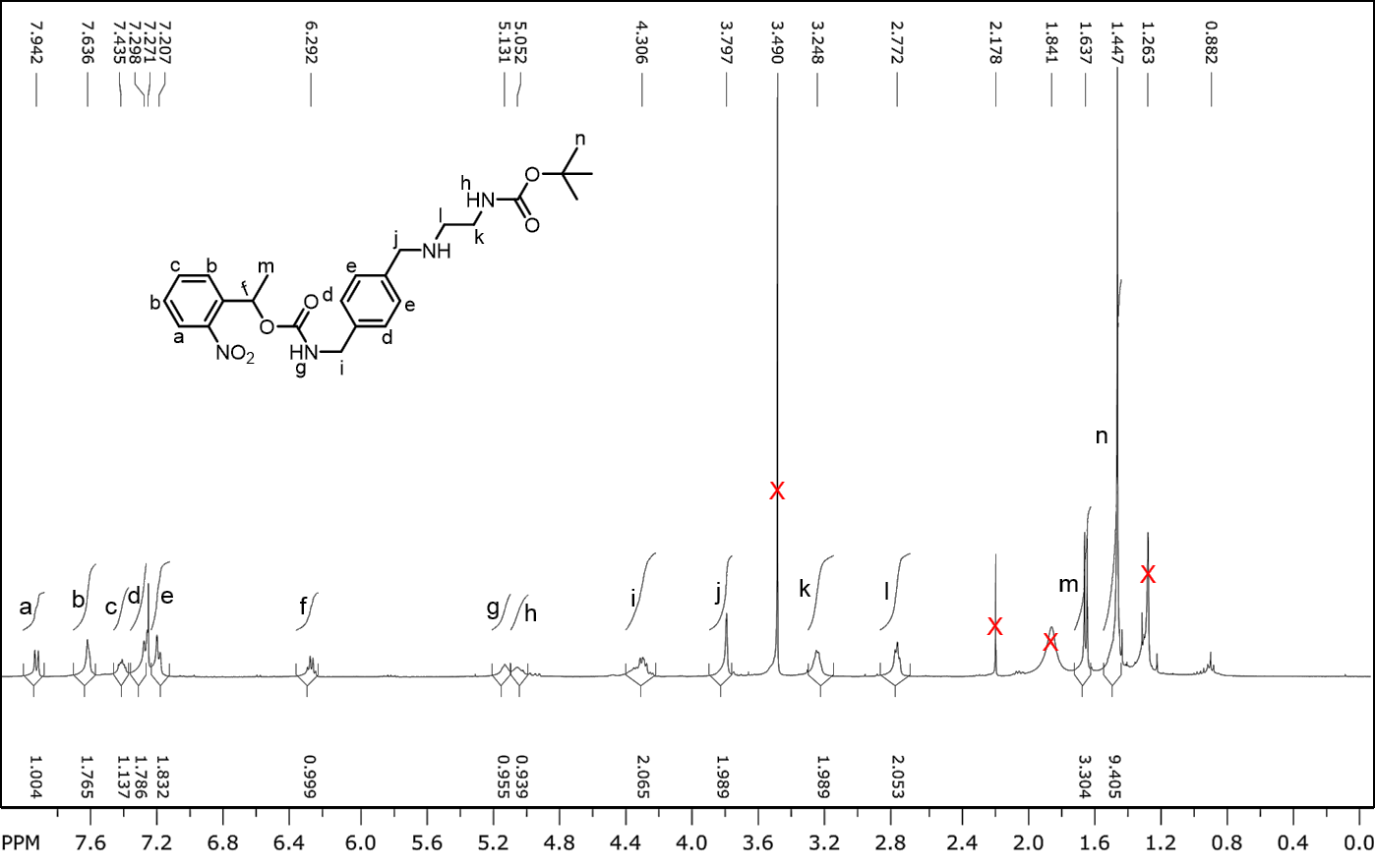

**Figure S28:** ^1^H NMR of compound 4 (400 MHz, CDCl_3_).

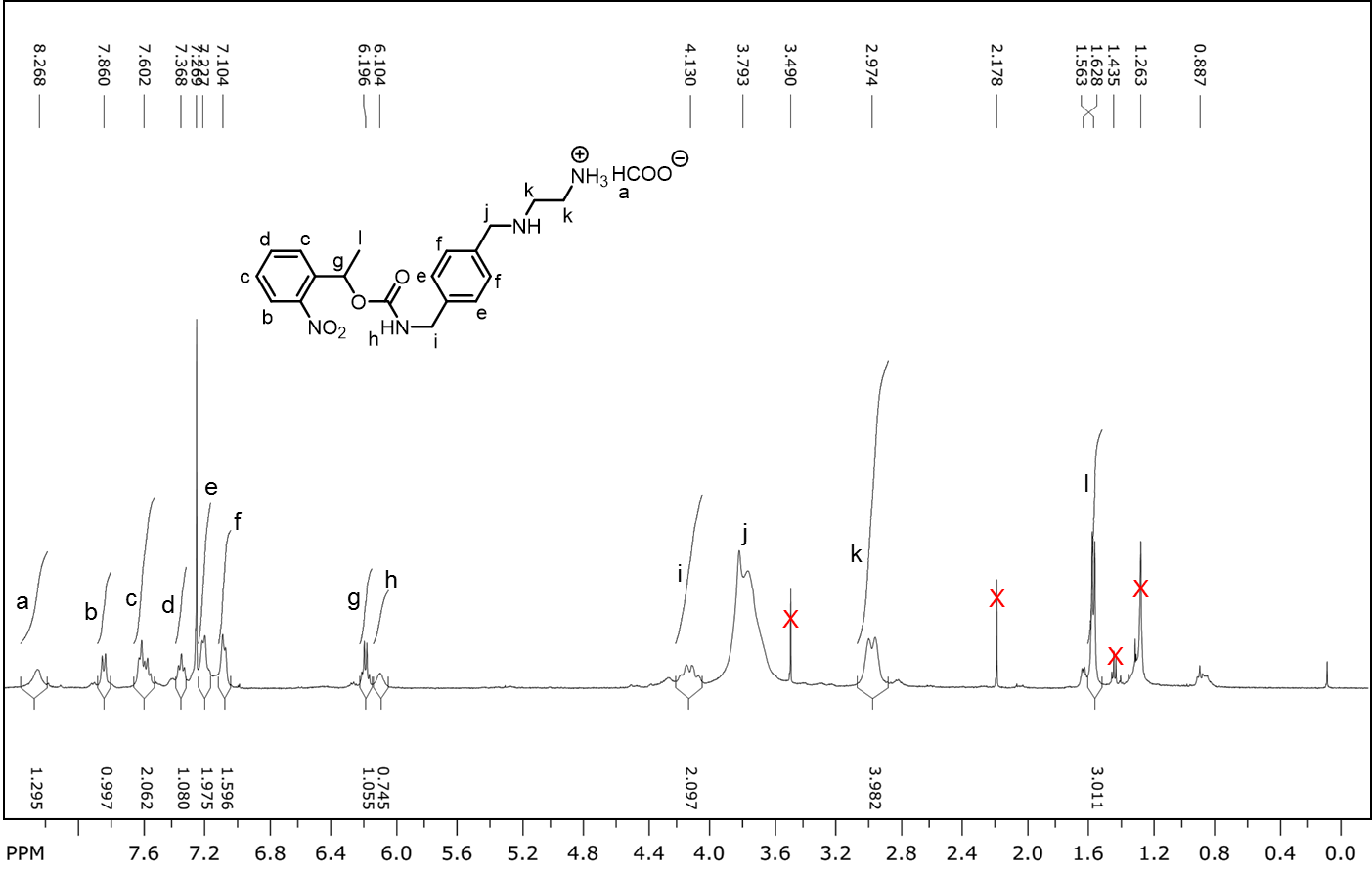

**Figure S29:** ^1^H NMR of compound 5 (400 MHz, CDCl_3_).

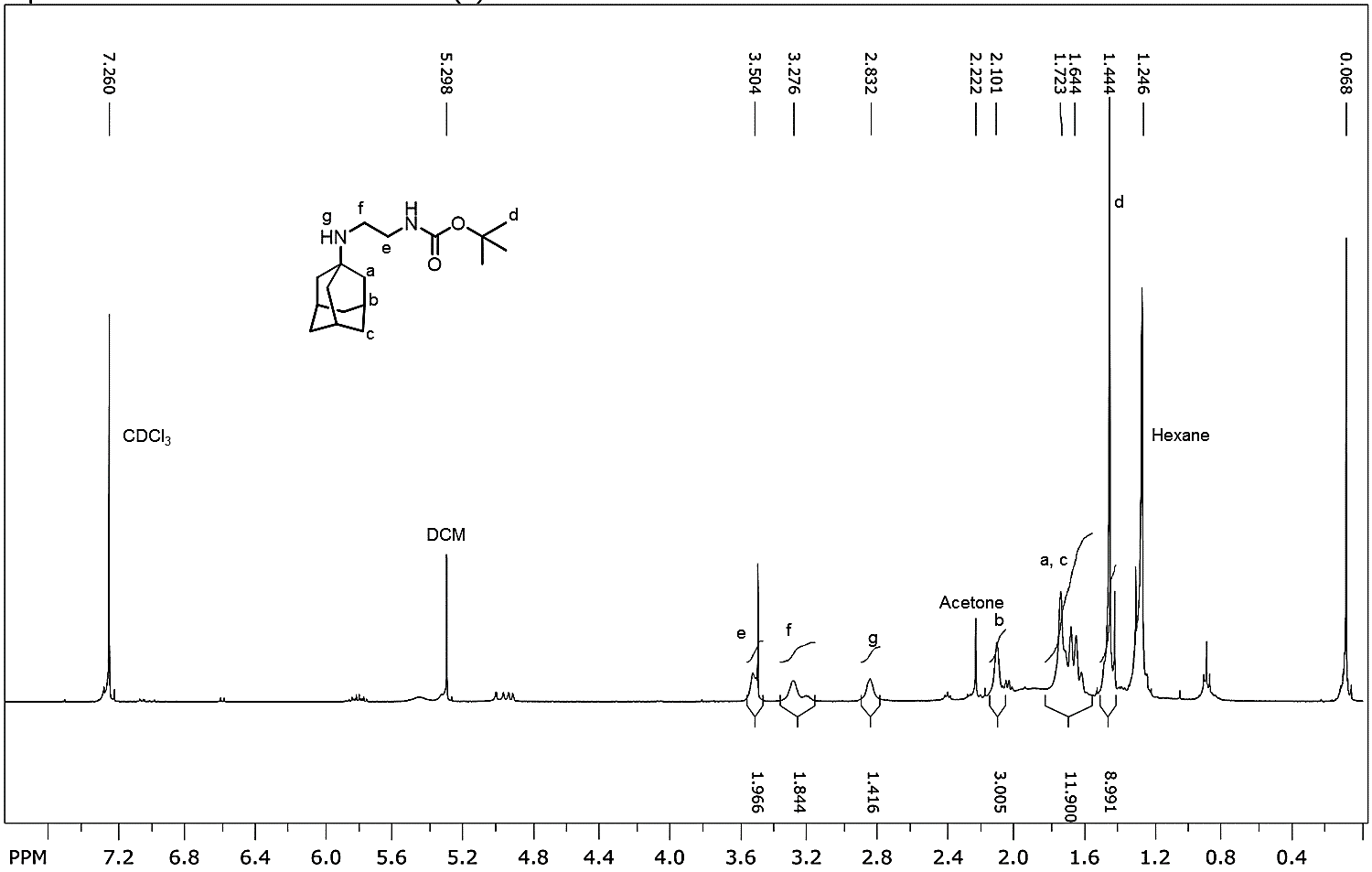

**Figure S30:** ^1^H NMR of compound 6 (400 MHz, CDCl_3_).

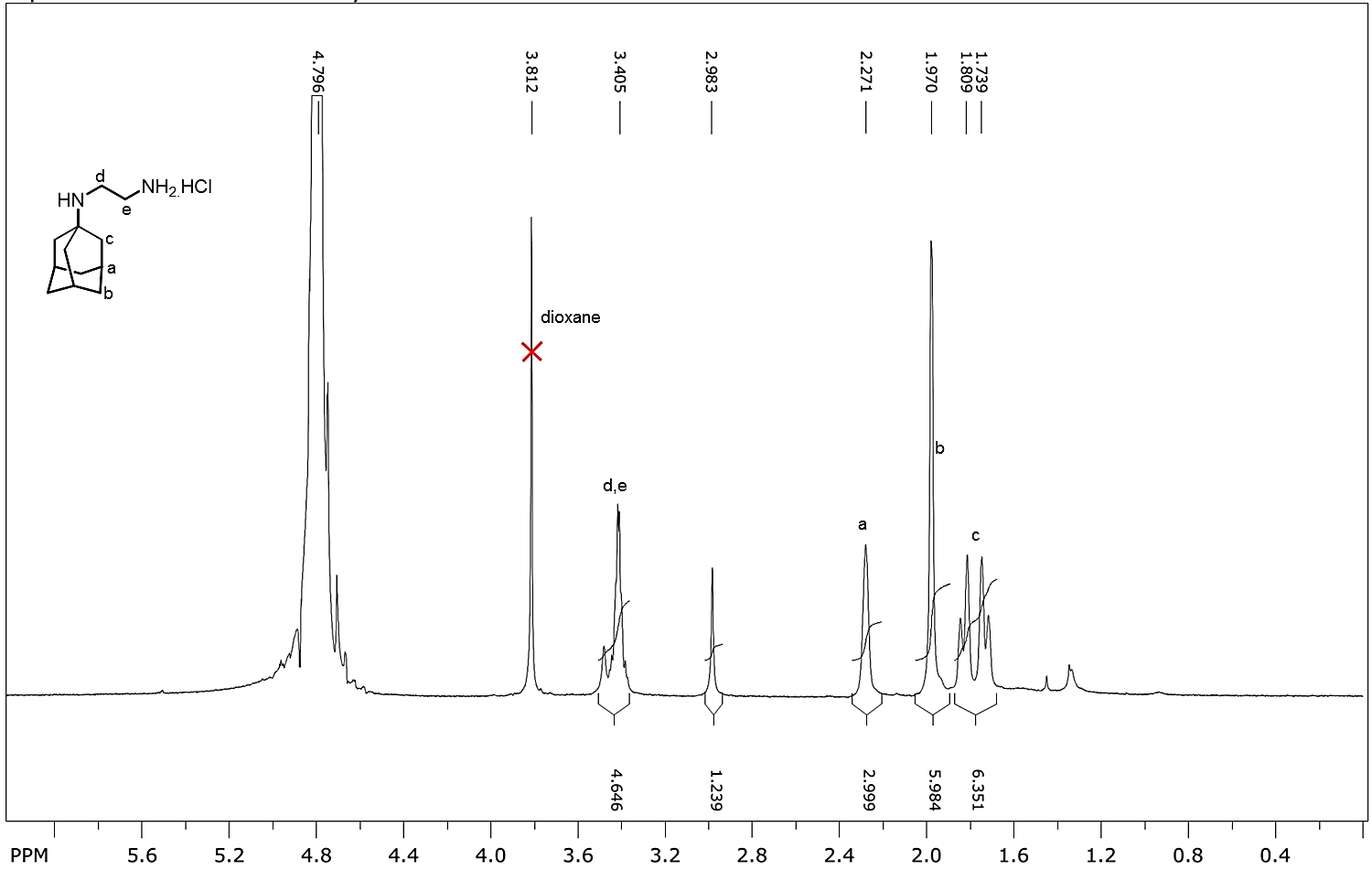

**Figure S31:** ^1^H NMR of compound 7 (400 MHz, D_2_O).

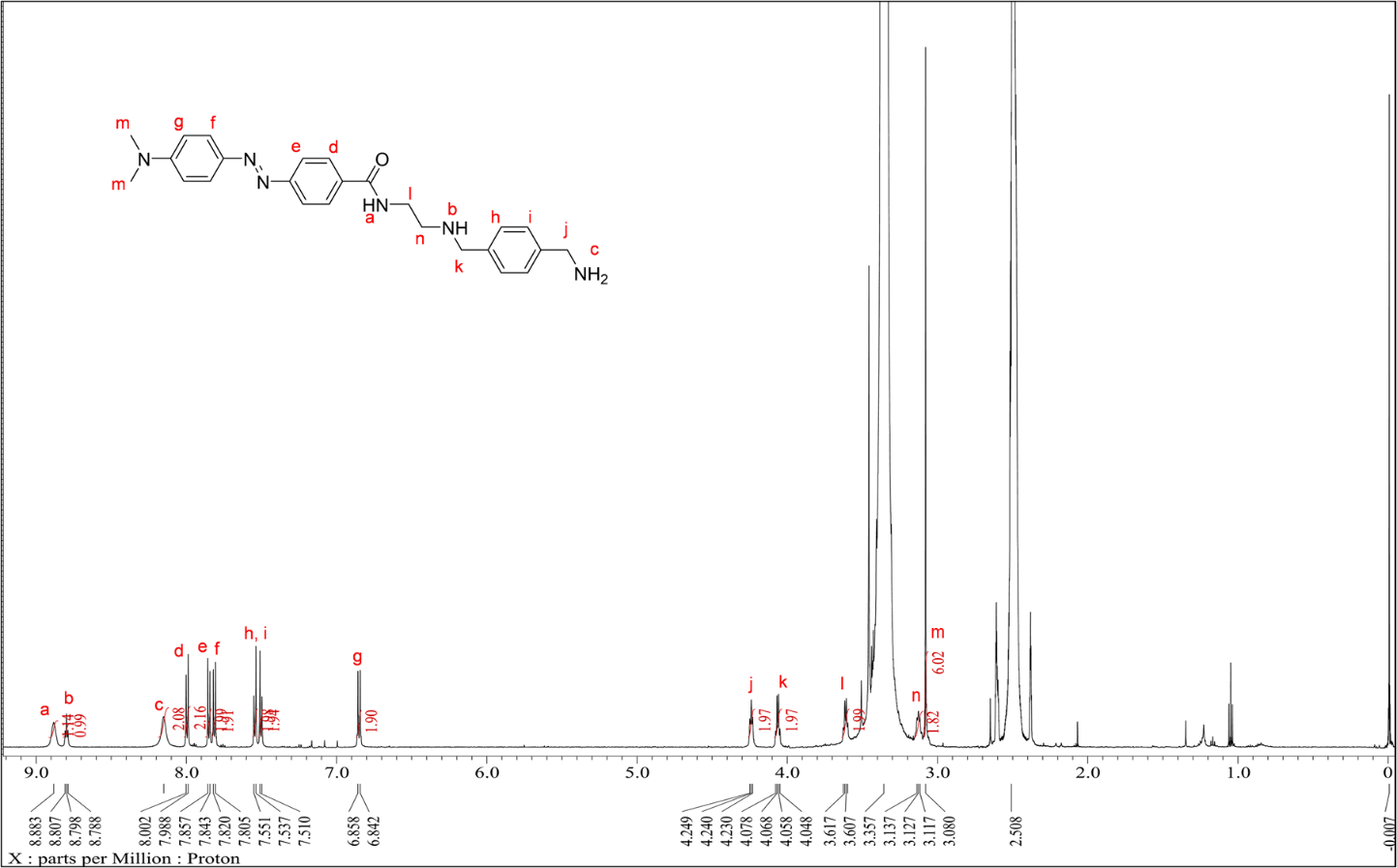

**Figure S32:** ^1^H–NMR spectrum of Xyl–Dabcyl conjugate (600 MHz, DMSO–d_6_).

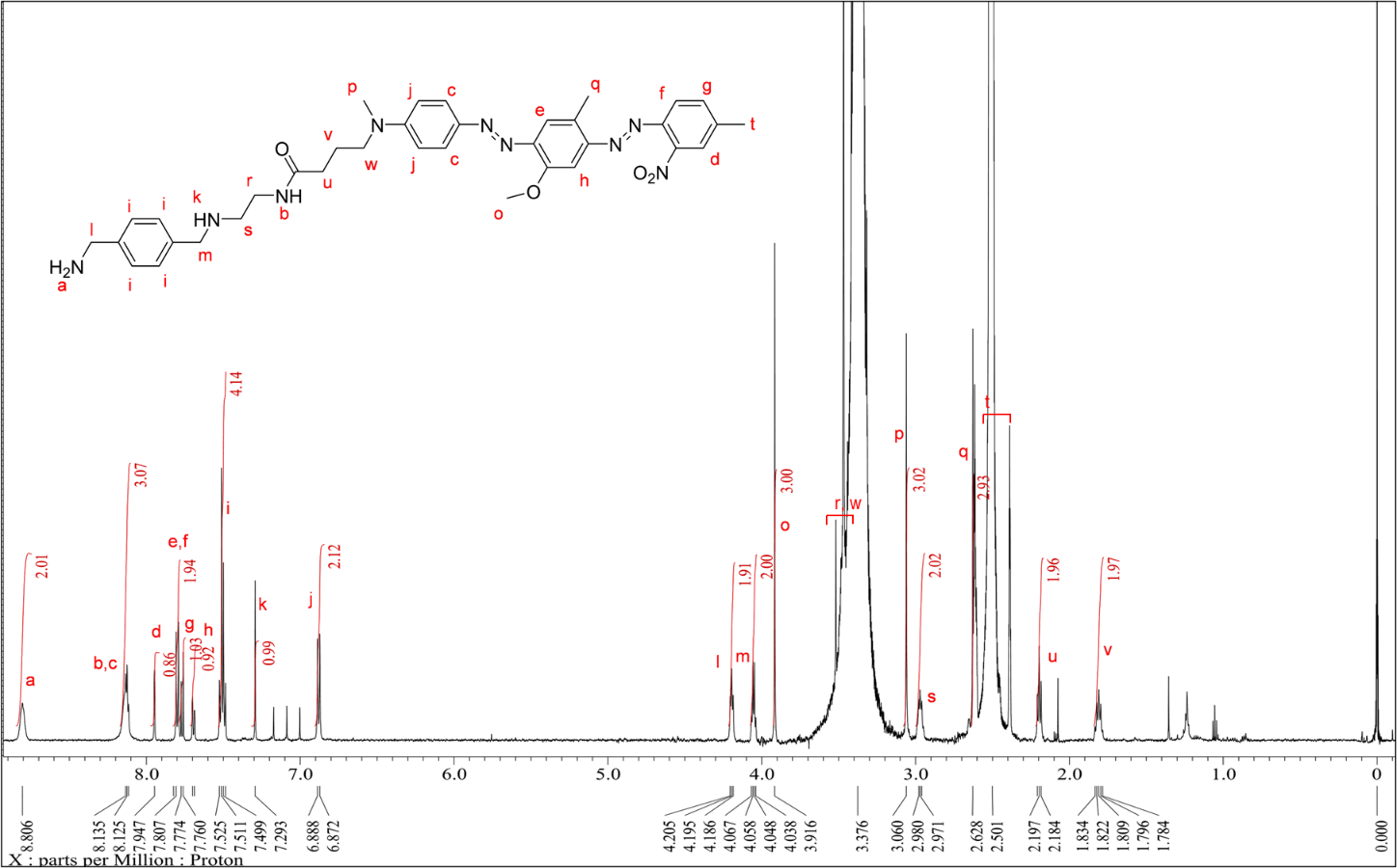

**Figure S33:** ^1^H–NMR spectrum of Xyl–BHQ1 conjugate (600 MHz, DMSO–d_6_).

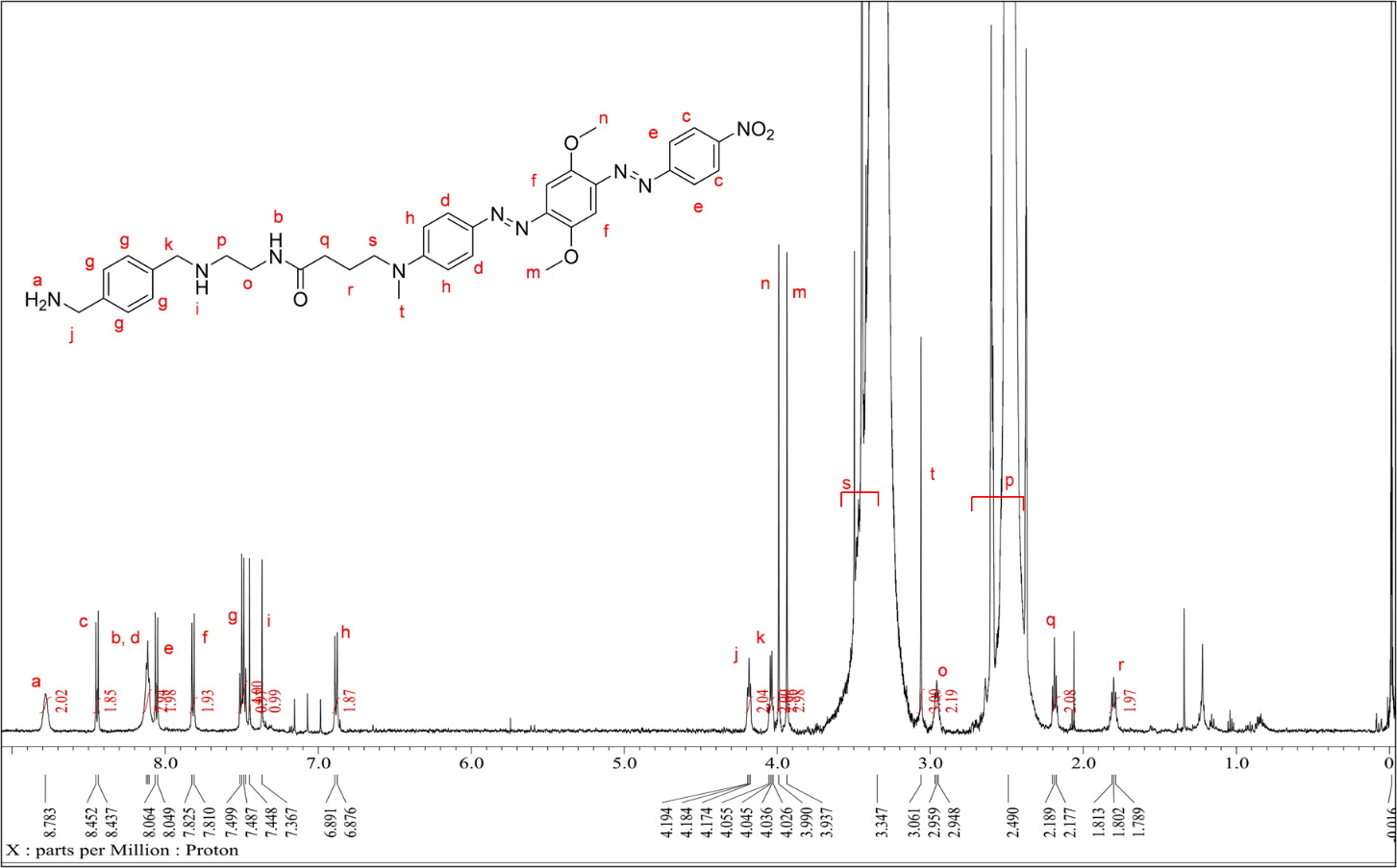

**Figure S34:** ^1^H–NMR spectrum of Xyl–BHQ2 conjugate (600 MHz, DMSO–d_6_).

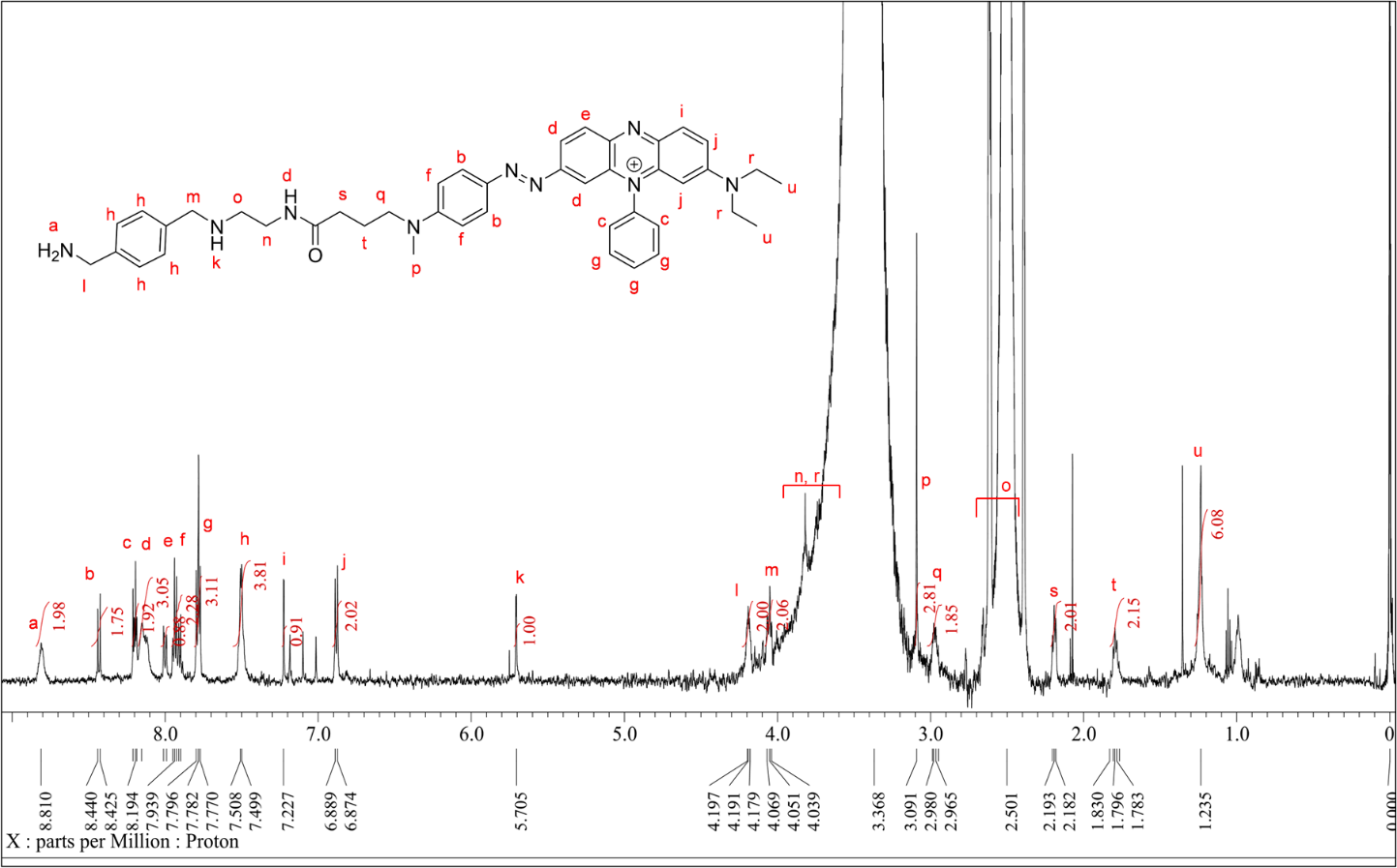

**Figure S35:** ^1^H–NMR spectrum of Xyl–BHQ3 conjugate (600 MHz, DMSO–d_6_).

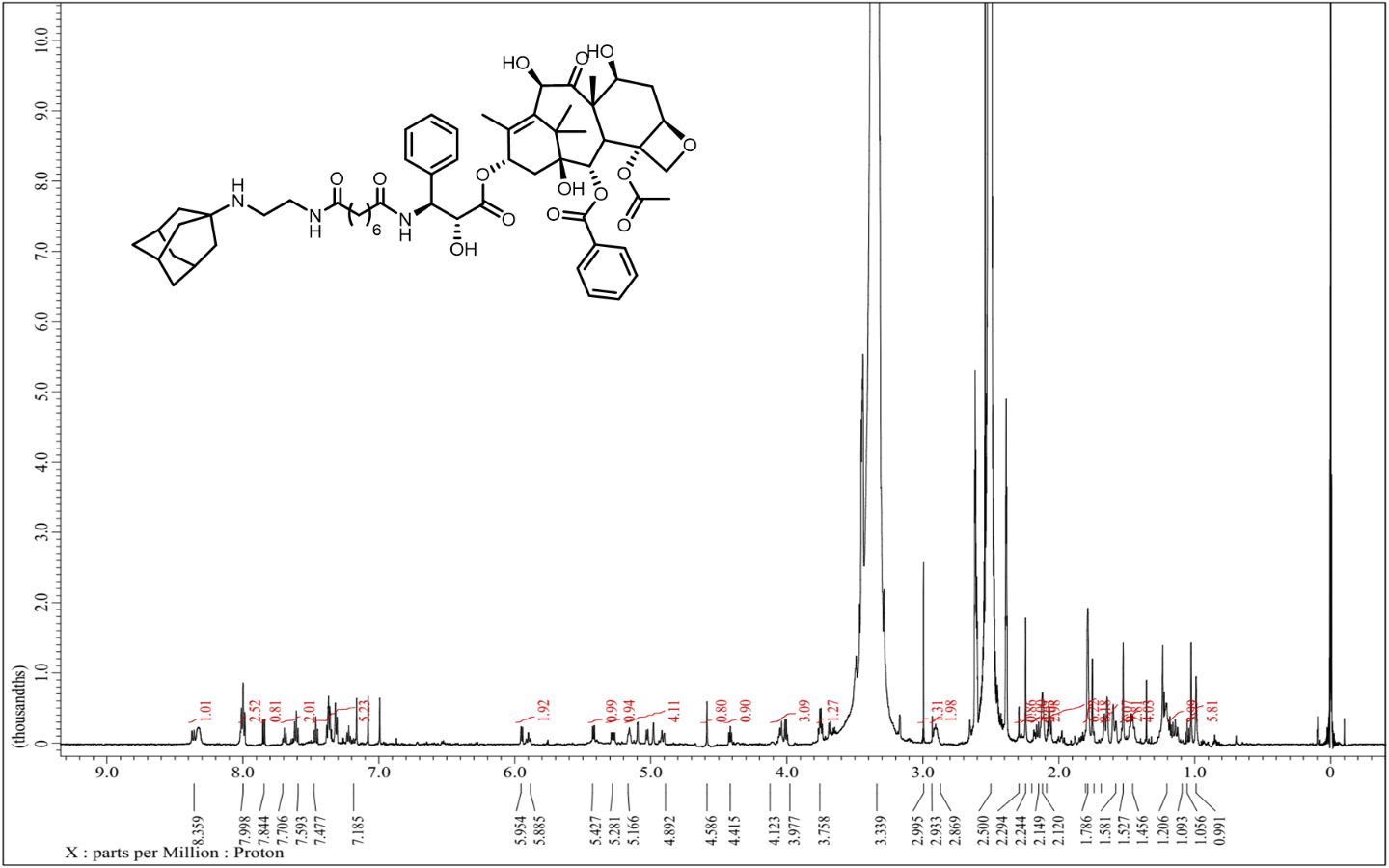

**Figure S36:** ^1^H–NMR spectrum of ADA conjugated taxol (600 MHz, DMSO–d_6_).

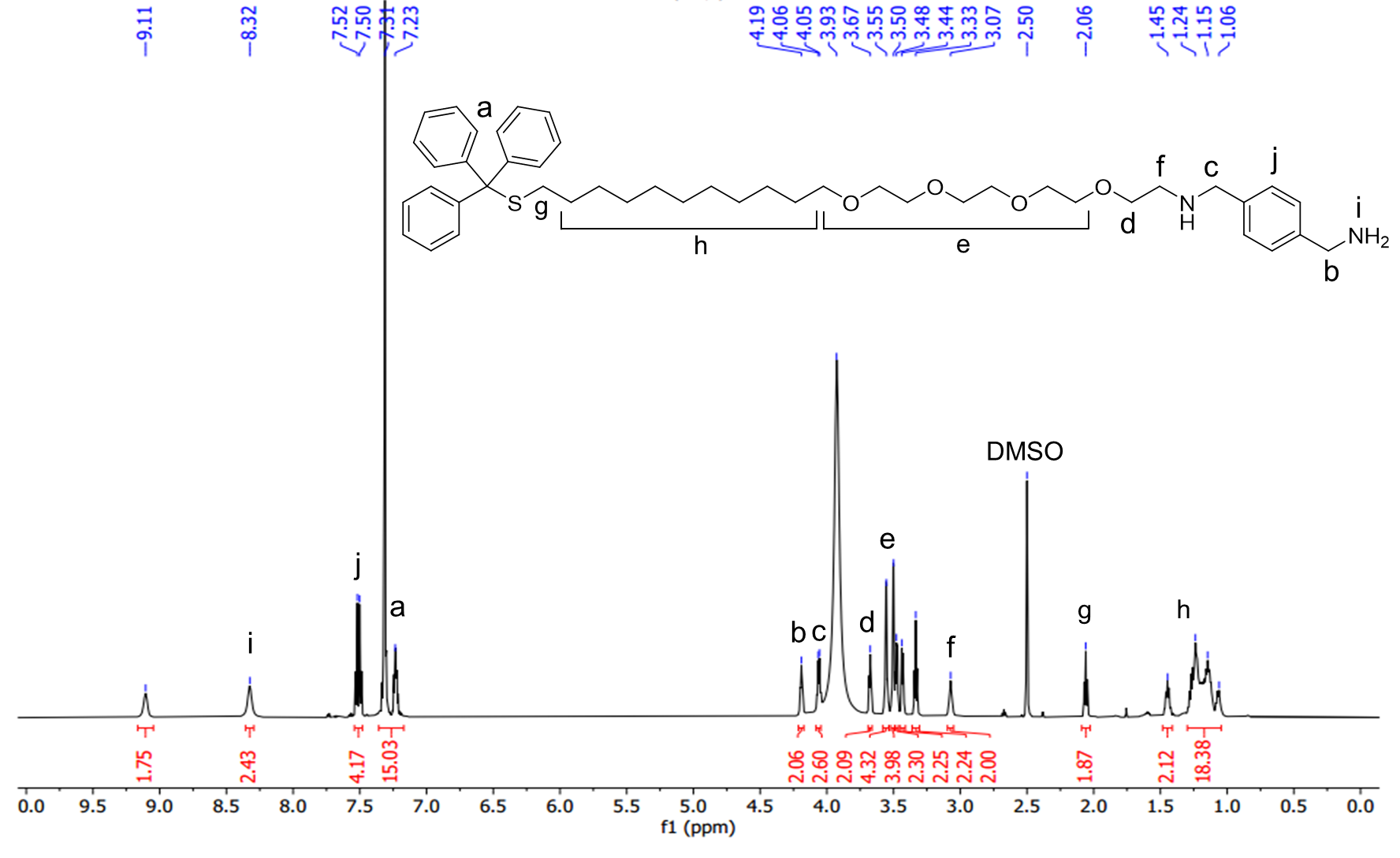

**Figure S37:** ^1^H–NMR spectrum of Compound 2**:** Trt-C11-TEG-XYL (600 MHz, DMSO–d_6_).

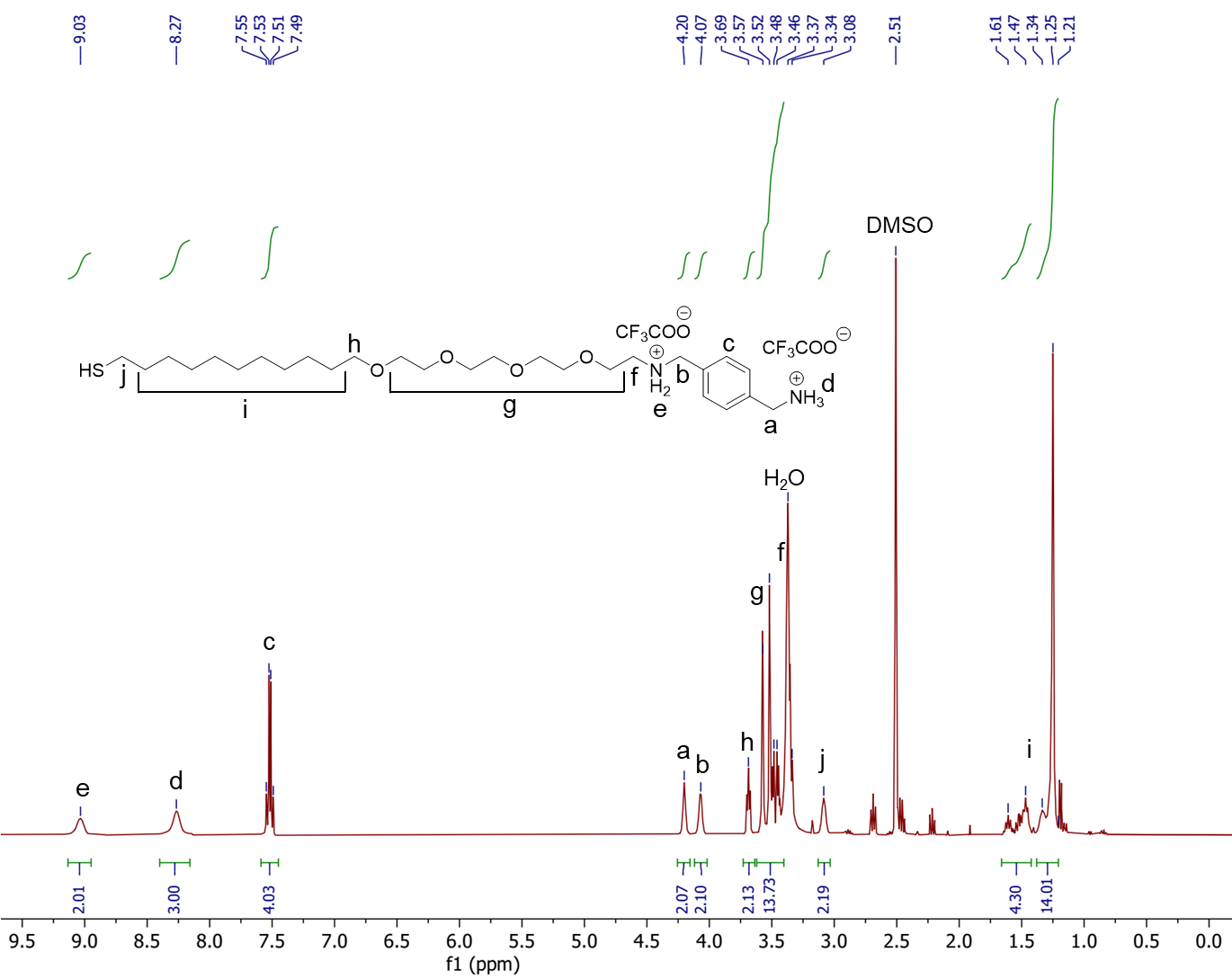

**Figure S38:** ^1^H–NMR spectrum of Compound 3: HS-C11-TEG-XYL (400 MHz, DMSO–d_6_).

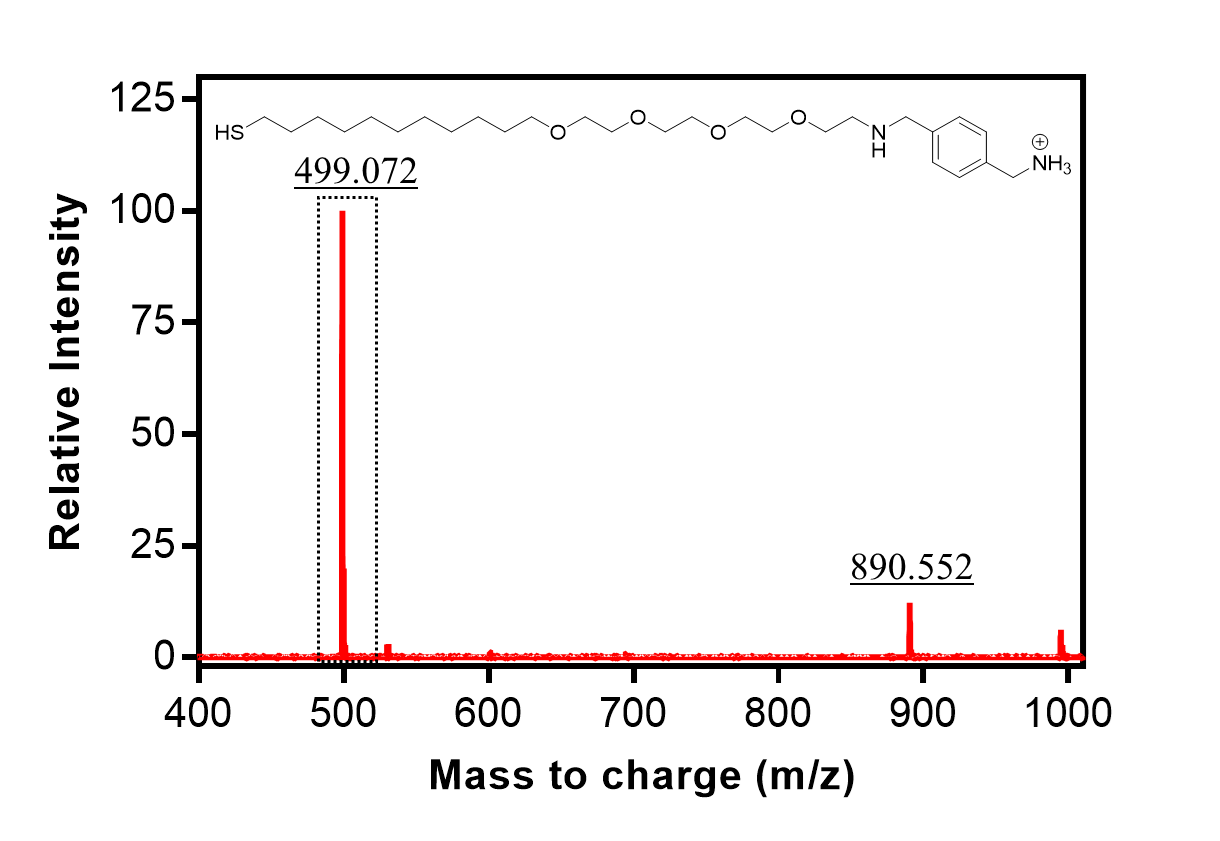

**Figure S39**: MALDI spectrum of XyL-AuNP showing a peak at m/z = 499.072 (observed) which corresponds to the theoretical mass value (m/z = 499.356) of thiol ligand of xylene diamine moiety.

**Table S13**: List of CB7 conjugated fluorophores that are used in this study^[7]^

| **Fluorophores** | **Synthesis references** |
| --- | --- |
| CB7-Coumarin | 7 |
| CB7-Fluorescein | 4 |
| CB7-BODIPY | 7 |
| CB7-TAMRA | 4 |
| CB7-Cy3 | 4 |
| CB7-Cy5 | 7 |
| CB7-SiR | 7 |
